# Computable Early *C. elegans* Embryo with a Data-driven Phase Field Model

**DOI:** 10.1101/2020.12.13.422560

**Authors:** Xiangyu Kuang, Guoye Guan, Ming-Kin Wong, Lu-Yan Chan, Zhongying Zhao, Chao Tang, Lei Zhang

**Affiliations:** Center for Quantitative Biology, Peking University, Beijing 100871, China; Department of Biology, Hong Kong Baptist University, Hong Kong 999077, China; State Key Laboratory of Environmental and Biological Analysis, Hong Kong Baptist University, Hong Kong 999077, China; Peking-Tsinghua Center for Life Sciences, Peking University, Beijing 100871, China; School of Physics, Peking University, Beijing 100871, China; Beijing International Center for Mathematical Research, Peking University, Beijing 100871, China

**Keywords:** morphogenesis, embryogenesis, *C. elegans*, phase field model, cell morphology, mechanoconnectome

## Abstract

Morphogenesis is a precise and robust dynamic process during metazoan embryogenesis, consisting of both cell proliferation and cell migration. Despite the fact that much is known about specific regulations at the molecular level, how cell proliferation and migration together drive the morphogenesis at the cellular and organismic levels is not well understood. Here, using *Caenorhabditis elegans* as the model animal, we present a data-driven phase field model to compute the early embryonic morphogenesis within a confined eggshell. By using three-dimensional time-lapse cellular morphological information generated by imaging experiments to set the model parameters, we can not only reproduce the precise evolution of cell location, cell shape and cell-cell contact relationship *in vivo*, but also reveal the critical roles of cell division and cellcell attraction in governing the early development of *C. elegans* embryo. In brief, we provide a generic approach to compute the embryonic morphogenesis and decipher the underlying mechanisms.

## INTRODUCTION

Development of metazoan embryo consists of cell proliferation, cell migration and cell differentiation, which is robust against perturbations and reproducible among individuals (Sulston et al., 1983; Jelier et al., 2016; Guan et al., 2019; Nie et al., 2020). The system evolves from a fertilized zygote to a multicellular structure with hundreds to thousands of cells and forms stereotypic three-dimensional spatial patterns (Sulston et al., 1983; Knust and Müller, 1998; Deneke et al., 2016). This dynamic process of morphogenesis is usually achieved by the precise control on cell divisions, mechanical interactions between cells (e.g., repulsion and attraction) (Lejeune and Linder, 2017; Lejeune et al., 2019) and other molecular-level regulations like cortical myosin flow, inhomogeneous cytomembrane adhesion and active actomyosin contractility (Naganathan et al., 2014; Sugioka and Bowerman, 2018; Nance et al., 2003; Lee et al., 2006; An et al., 2017). Thus, a fundamental question in developmental biology is “Will the egg be computable?” (Wolpert, 1994).

The eutelic nematode *Caenorhabditis elegans* has invariant developmental procedure at the cellular level, i.e., each cell can be identified based on its lineage history and has highly reproducible division timing, migration trajectory and cell fate among individual embryos (Sulston et al., 1983; Guan et al., 2019). Morphogenesis in *C. elegans* starts as early as fertilization and the cell specification happens intensively afterwards. For example, there are 4 cell types at 4-cell stage and 6 cell types at 8-cell stage – the cell types are diversified by consecutive asymmetric divisions of the germline stem cell (i.e., P0, P1, P2 and P3) and contact-based cell-cell signaling transduction (e.g., Wnt and Notch) (Gönczy and Rose, 2005; Hubatsch et al., 2019; Thorpe et al., 1997; Priess, 2005). These cells of different sizes, shapes and fates migrate, communicate and interact with each other, making the correctness of their positions and contact relationships extremely momentous.

Extensive theoretical studies have been carried out on *C. elegans* as a model organism for developmental biology and made great efforts to computationally rebuild the morphogenetic behaviors *in silico*, with an ultimate goal of permitting virtual experiments with various assignments on all kinds of physical parameters to facilitate in-depth interpretation and understanding on embryonic morphogenesis. To list a few, a multi-particle model was designed to analyze the structure evolution at 4-cell stage using groups of interactive particles representing cell membranes, which however, contained dozens of system parameters that were difficult to measure or fit (Kajita et al., 2002; Kajita et al., 2003). Later, several coarse-grained models were proposed to reproduce cell positions at up to ~50-cell stage, which simplified the cells into mass particles and ignored most of the morphological features as well as the morphogenetic processes (Fickentscher et al., 2013; Fickentscher et al., 2016; Yamamoto and Kimura, 2017; Tian et al., 2020; Guan et al., 2020). Recently, deep-learning methods were applied to extract information from 4-dimensional cell motion data, which by itself did not address the physical and biophysical mechanistic questions (Wang et al., 2018; Wang et al., 2019). On the other hand, phase-field methods have been widely applied to simulate single-cell dynamics including cell morphology and motility (Wang et al., 2017; Aras et al., 2018; Cao et al., 2019; Tao et al., 2020), and to mimic the evolution of multicellular systems on the phenomenological level, such as early development in sea cucumber and nematode embryos (Nonomura, 2012; Löber et al., 2015; Akiyama et al., 2018; Jiang et al., 2019).

In this paper, we present a data-driven phase field approach to reconstruct the cell-packing dynamics in early *C. elegans* embryogenesis and investigate the strategies/principles accounting for the stereotypic morphogenetic patterns, by combining both *in vivo* cell morphology data and *in silico* phase field model. We first collect cell-resolved morphological data from our previous work (Cao et al., 2019), including cell location, cell shape, cell volume, cell surface area, and cellcell contact relationship and area, and eggshell shape, which were originally acquired from three-dimensional (3D) timelapse imaging on 4 wild-type embryos. Next, we develop a data-driven phase field model to reproduce the morphogenetic transformation from 1- to 4-cell stage within a confined eggshell by considering minimal mechanical constrictions, setting the cell volume and division orientation, and fitting the parameters according to experimental data. We predict the asymmetry of attractions among the 5 contacted cell pairs in the diamond-shaped 4-cell structure, which is in line with the previous knowledge about non-uniform distribution of cell-cell adhesive protein HMR-1, and validate the prediction by fluorescent imaging experiment. We further simulate the cell division, deformation and motion from 4- to 8-cell stage by introducing self-determined mechanisms on cell-cell attraction matrix and cell division timing to guide the spatial development. A frequently-occurring defective phenotype called “planarization” is found in the compressed embryo when developmental programs are disturbed, which is further verified by laser-ablated and RNAi-treated embryos. Lastly, we identify three key physical factors critical for the precise and robust evolution of the 3D structure and the cell-cell contact map: cell division timing, cell division orientation and cell-cell attraction matrix.

## RESULTS

### *In Vivo* Data Collection and *In Silico* Framework Establishment

A total of 18 wild-type embryo samples imaged by 3D time-lapse confocal microscopy are collected from datasets produced previously, along with their outputs of membrane segmentation, nucleus tracing and cell lineaging (Table S1) (Guan et al., 2019; Cao et al., 2019), providing multi-dimensional cell-level developmental properties from 1- to 8-cell stages (Figure 1A; see STAR Methods). The cell division orientation, cell volume and eggshell shape are quantified and directly inputted into simulation as predetermined parameters (Figure 1B; Table S2), while the cell morphology including cell location, cell shape, cell surface area, and cell-cell contact relationship and area, is used to verify our model by comparison to the simulation results (Tables S3-S6). It should be pointing out that the early *C. elegans* development can be divided into separate stages with exact cell numbers, which are achieved by an invariant division sequence highly conserved within *Caenorhabditis* species (Zhao et al., 2008; Memar et al., 2019).

**Figure 1.**
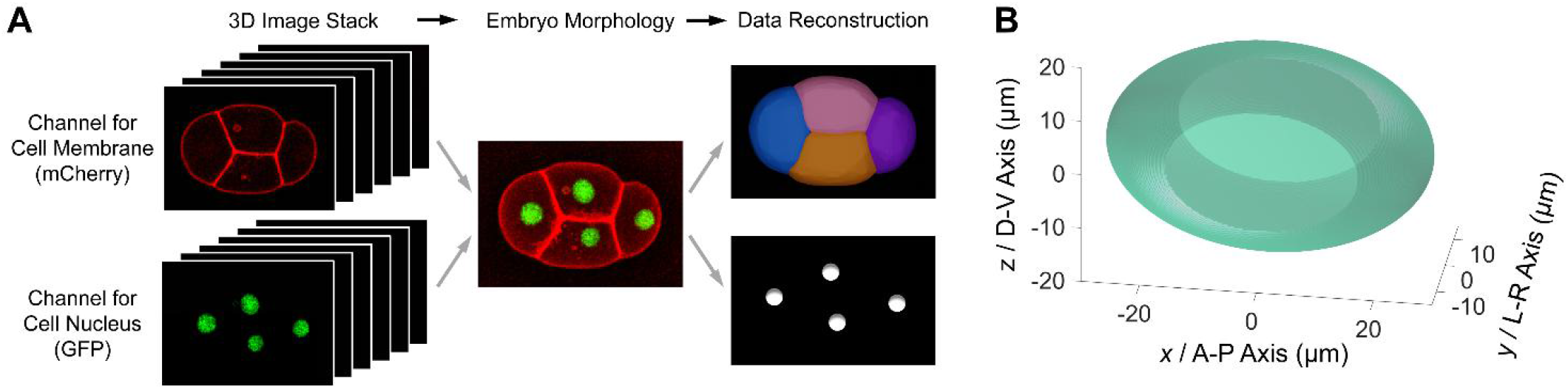
Reconstruction of *In Vivo* Morphological Information. (A) *In vivo* imaging experiment and quantification of 3D cell morphology and cell position, by identifying fluorescent signal in cell membrane (mCherry, red) and cell nucleus (GFP, green) (illustrated with strain ZZY0535; Chen et al., 2018). (B) An ideally rebuilt eggshell under compression, based on size measurements on 4 wild-type embryos (Cao et al., 2019).

To simulate the morphological and morphogenetic dynamics of a multicellular system, we develop a phase field model which uses 3D phase field *ϕ_i_* to describe each individual cell *i* (*i* = 1,…, *N*), where *N* is the cell number, subjected to multiple forces, including cell surface tension ***F***_ten_, cell-eggshell and cell-cell repulsion ***F***_rep_, cell-cell attraction ***F***_atr_ and cell volume constriction ***F***_vol_ i.e.,

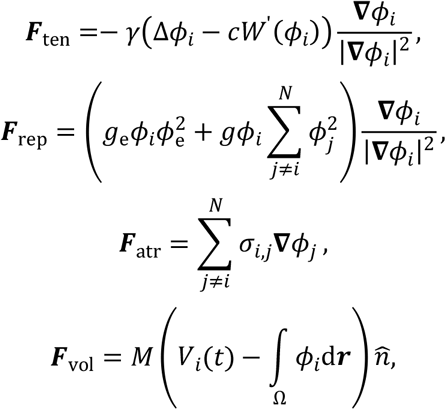

where *γ* is the cell surface tension and *c* is a positive coefficient related to the thickness of cell boundary between interior and exterior of cells; *W*(*ϕ*) = *ϕ*^2^(*ϕ* – 1)^2^ is a double-well potential with minima at *ϕ* = 1 and *ϕ* = 0; *g*_e_ and *g* are positive coefficients, denoting the strength of cell-eggshell and cell-cell repulsive energy respectively; *σ_i,j_* is a non-negative coefficient and positively associated with the attraction intensity between the *i*-th and *j*-th cells; *M* is a positive coefficient which denotes the volume constraint strength and 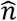 is the unit normal vector at interface which orients inward; *V_i_*(*t*) denotes the prescribed volume for the *i*-th cell.

Cell division is implemented as instantaneous bisection of phase field *ϕ_i_* by a splitting plane, whose direction and location are determined by cell volume segregation direction and ratio obtained from experimental measurements (see STAR Methods). The whole system evolves over developmental time *t* inside an overdamped environment as follows:

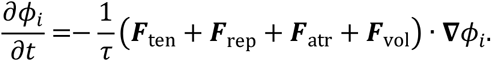

where *τ* is the viscosity coefficient of embryo’s internal environment. To build up a minimal model that has the least physical constraints but outlines the most significant characteristics of a developing embryo, we will modify the system progressively according to the *in vivo* cell morphology data. We initially set the intercellular attraction *σ_i,j_* = 0 for all the contacted cell pairs, which would be investigated in depth later as a high-dimensional factor to diversify the path of morphogenesis (hereafter referred to as “developmental path”). Additionally, the independent physical coefficients *γ, c*, and *g*_e_ are optimized and fixed by fitting the structural characteristics observed in experiment from 1- to 4-cell stages (see STAR Methods).

### The Morphogenetic Procedure from 1- to 4-Cell Stages can be Reconstructed Only Using Cell Surface Tension, Cell-Eggshell and Cell-Cell Repulsion and Cell Volume Constriction

We start with reconstructing the embryo morphology from 1- to 4-cell stages. The zygote P0 first divides asymmetrically under polarization induced by sperm entry, generating somatic founder cell AB (with larger size) and germline stem cell P1 (with smaller size). Next, AB proceeds symmetric division into ABa (in anterior) and ABp (in posterior), while about 2.5 minutes later P1 proceeds asymmetric division again to generate somatic founder cell EMS (with larger size) and germline stem cell P2 (with smaller size) (Sulston et al.,1983; Hubatsch et al., 2019; Brauchle et al., 2003). Owing to cell polarization and signaling transduction (i.e., Wnt from P2 to EMS and Notch from P2 to ABp), the 4 cells at this moment already acquire differentiated fates (Gönczy and Rose, 2005; Priess, 2005; Thorpe et al., 1997). Thus, correct formation of the diamond-shape 4-cell structure, especially for the cell-cell contacts with sufficient stability and area, is crucial for intercellular signaling and lays a foundation for further embryogenesis.

In the simulations of 1- to 4-cell stages, cell division orientation (i.e., cell volume segregation direction and ratio) during cytokinesis are all in accordance to those measured in experiments (Table S2). To set up a minimal model in the very beginning, cell-cell attraction is ignored in all the cell-cell contact pairs (i.e., *σ_i,j_* = 0), resulting in the 1- to 4-body system only controlled by cell surface tension, cell-eggshell and cell-cell repulsion and cell volume constriction. Notably, the simulated embryo morphology is highly similar to the one *in vivo*, for all the 2-, 3- and 4-cell stages (Figure 2A; Movie S1). The first cell division P0 → AB & P1 proceeds along the major axis of eggshell and establishes the anterior-posterior (*x* → A-P) axis in *C. elegans* embryo (Gönczy and Rose, 2005). The second cell division AB → ABa & ABp has spindle orientation initially perpendicular to but quickly reoriented to A-P axis, depending on the direct contact between AB and P1 and the mechanosensitive redirection of cortical myosin flow (Sugioka and Bowerman, 2018). The bias in AB division orientation contributes to the establishment of dorsal-ventral (*z* / D-V) axis, and leads to a transient three-body structure together with P1. At 3-cell stage, P1 is elongated and connected to both ABa and ABp, which is also reproduced in the simulation (Figure 2A). The last cell division P1 → EMS & P2 proceeds approximately parallel to A-P axis to place the somatic founder cell EMS anteriorly and germline stem cell P2 posteriorly, making the latter cell contacted with both ABp and EMS but not with ABa, for accurate fate induction (Priess, 2005; Thorpe et al., 1997).

**Figure 2.**
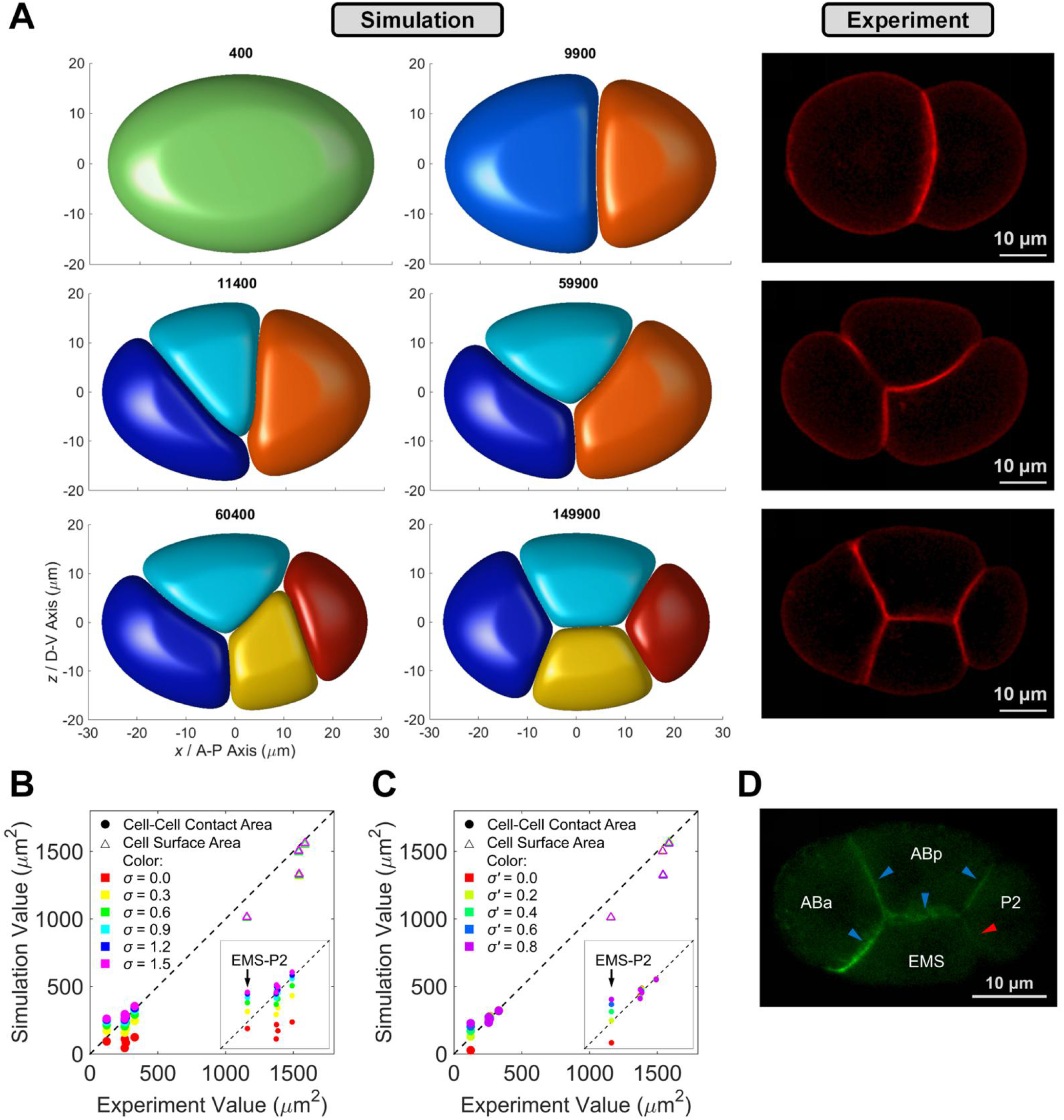
Embryo Morphology Reconstruction from 1- to 4-Cell Stages. (A) Comparison of embryo morphology between simulation and experiment from 1- to 4-cell stages (view from *y I* leftright axis); the 1^st^ and 2^nd^ columns, cell-arrangement progression in phase-field simulation; the 3^rd^ column, a live embryo with mCherry fluorescence on cell membrane (strain ZZY0535; Chen et al., 2018); scale bar, 10 μm. (B) Comparison of cell surface area and cell-cell contact area between simulation and experiment at 4-cell stage, with globally symmetric attraction *σ* applied on all the cell-cell contacts (*σ* = 0.0, 0.3, 0.6, 0.9, 1.2 and 1.5); inset, range from 0 to 500 μm^2^ in both coordinates. (C) Comparison of cell surface area and cell-cell contact area between simulation and experiment at 4-cell stage, with globally symmetric attraction *σ* =0.9 applied on cell-cell contacts of ABa-ABp, ABa-EMS, ABp-EMS and ABp-P2, and different asymmetric attraction *σ*’ on contact of EMS-P2 (*σ*’ = 0.0, 0.2, 0.4, 0.6 and 0.8); inset, range from 0 to 500 μm^2^ in both coordinates; minimal average relative deviation ≈ 0.06 when *σ*’ = 0.2. (D) Distribution of membrane-attached E-cadherin HMR-1 at 4-cell stage, with substantially higher accumulation in cellcell contacts of ABa-ABp, ABa-EMS, ABp-EMS, ABp-P2 (pointed by blue arrows) than that in EMS-P2 contact (pointed by red arrow) (strain LP172; Marston et al., 2016); scale bar, 10 μm.

There are two notable differences between simulation and experiment. First, the contact interface between AB and P1 at 2-cell stage is slightly protruding toward the larger cell AB in simulation, but is in opposite direction in experiment (Figure 2A). This distinction is raised by asymmetry of surface tension between cells (i.e., *γ*_AB_ and *γ*_P1_), in other words, the cell surface tension of AB is plausibly stronger than that of P1 in real embryo so that AB can maintain more spherical instead of P1, however, we treat all the cells with totally same level of cell surface tension (*γ*_AB_ = *γ*_P1_ = 0.25) for simplicity. Both the model and simulation show that the increasement in *γ* can strengthen a cell’s surface tension and its ability to maintain spherical (Figure S1; see STAR Methods). Whether AB has stronger cell surface tension than P1 needs to be verified by further experiment, for example, by fluorescent imaging on cytoskeleton density or direct measurement on surface tension (Xia et al., 2014). The second difference is the distinguishable hollows at junction points among cells, which can be found from 3- to 4-cell stages and are generated by cooperation of relatively strong cell surface tension and weak cell-cell attraction, implying potential limitations in the current model such as quantification of cell-cell contact area (Figure 2A). Influenced by cell surface tension, cell shape tends to be spherical to minimize the surface energy as well as surface area, weakening the cells’ deformation and generating openings among them. Nevertheless, this defect can be overcome when cell-cell attraction is globally introduced (Figure S2).

### Introduction of Asymmetric Cell-Cell Attraction can both Rebuild the Contact Area Between Specific Cells and Infer the Inhomogeneous Distribution of Adhesive Protein in Real Embryo

Since attractive force always exists between contacted cells (e.g., generated by intrinsic adhesion of cell membrane) and the global attraction has been proved to be essential for eliminating the hollows mentioned above (Figure S2), we next take this interaction into account and compare both cell surface area and cell-cell contact area in simulation with their corresponding values measured experimentally. Here, we first set up the global attraction coefficient *σ* = 0.0, 0.3, 0.6, 0.9, 1.2 and 1.5, and simulate their resultant 4-cell structures respectively. The systems are allowed to relax until reaching mechanical equilibrium (duration = 10000 time steps). Interestingly, the cell surface area is not sensitive to the global attraction for that it’s dominantly determined by a cell’s prescribed volume (all changes of relative deviation < 0.02), while the cell-cell contact area rises with the increase of global attraction (all changes of relative deviation > 0.87) (Figure 2B; Table S7). We then compare both the cell surface area and cell-cell contact area between simulation and experiment, to determine the optimal value of *σ* by minimizing the average relative deviation from experiment to simulation. All the surfaces and interfaces acquire area that moderately fits the experimentally measured value when *σ* = 0.9 (all relative deviations < 0.14), except the contact area between EMS and P2 (relative deviation > 0.91). The EMS-P2 contact has substantially larger area than the actual one, indicating that the attraction between them should be much weaker than the other 4 cell pairs. *In vivo* imaging on E-cadherin HMR-1 at 4-cell stage reveals a significant lower localized accumulation in EMS-P2 interface than that in ABa-ABp, ABa-EMS, ABp-EMS and ABp-P2 interfaces, in accordance with a recent report (Figure 2D) (Yamamoto and Kimura, 2017). This asymmetry of cell-cell attraction was proposed to affect the final spatial pattern (i.e., “H”, “I”, “T” and diamond shapes in Figure S3 and Movies S2-S4), contributing to stability and diversity of the 4-cell structure.

Considering the known function of attraction asymmetry and the oversized area in EMS-P2 interface, we further screen a smaller attraction coefficient *σ*’, which is specifically assigned to the EMS-P2 contact, and eventually obtain the optimal combination *σ* = 0.9 and *σ*’ = 0.2 with an area of EMS-P2 contact close to its actual value (average relative deviation < 0.07, all relative deviations < 0.14) (Figure 2C; Table S8). Hence, the attraction asymmetry is necessary for comprehensive reconstruction of embryo morphology, especially for the physical contact between cells. The area of direct contact is critical for intercellular communication which widely exists during embryogenesis and functions in regulation of both fate specification and division orientation (Thorpe et al., 1997; Priess, 2005; Walston et al., 2004). The cell-cell contact area needs to be mediated accurately to transduct signaling at an appropriate range of intensity, for example, to reach a responsive threshold (Chen et al., 2018; Shaya et al., 2017). The weak attraction on EMS-P2 interface significantly controls their contact area and probably ensures the precision of Wnt signaling between them, which coordinates the division orientation of EMS and induce two cell fates in its progenies (i.e., endoderm for E cells and mesoderm for MS cells) (Thorpe et al., 1997). In a word, with asymmetric cell-cell attraction introduced, we successfully reproduce the whole evolution of embryo morphology as well as cell-cell contact area from 1- to 4-cell stages.

### A Self-Determined Binarized Cell-Cell Attraction Matrix is Used for Later Developmental Stages

Before gastrulation onset (i.e., 26-cell stage), *C. elegans* embryo always undergoes a series of sequential cleavage, obeying invariant distinct order as P0 → AB → P1 → AB2 → EMS → P2 → AB4 → MS & E → C → AB8 & P3 → MS2 (Guan et al., 2019). On the basis of the minimal model including cell surface tension, cell-eggshell and cell-cell repulsion, cell-cell attraction and cell volume constriction, we next attempt to reproduce the cellular migratory behaviors observed at those later stages, so as to test the performance of the phase field model and uncover the underlying strategies/principles for precise and robust embryonic morphogenesis. Since the attraction between specific cells is already asymmetric at 4-cell stage and proved to influence the cell-arrangement pattern substantially (Yamamoto and Kimura, 2017; Guan et al., 2020), here we use two simplified assumptions to introduce the setting of cell-cell attraction matrix and make the multi-stage simulation self-driven.

First, the cell-cell attraction matrix is binarized. As the attractive force exhibits relatively stronger or weaker intensity between different contacted cell pairs (Figure 2D) (Yamamoto and Kimura, 2017), for simplicity, we binarize these two states using the parameter value acquired from 4-cell stage (i.e., *σ* = 0.9 and *σ*’ = 0.2), which represent strong and weak attraction respectively. This approximation reduces the dimension of cell-cell attraction matrix and limits the possibility of developmental path, so as to achieve an affordable computational cost. The developmental path is consequently determined by a combination of binarized states in all the contacted cell pairs. Moreover, for the later stages, we only examine the cellcell contact relationship but not the exact area of contact interface because fitting this quantitative property needs delicate and massive work both experimentally and computationally (Figures 2B, 2C and S2; Tables S7 and S8).

Second, the attraction between non-sister cells and between sister cells is regarded as strong and weak respectively. It was previously found that some sister cells can be separated transiently (contact area = 0) during collective motion, suggesting that the attraction between them is not enough to keep them connected continuously, even though they are already adjacent to each other initially due to their sisterhood. For instance, ABpl and ABpr couldn’t maintain their physical contact in 2 out of the 4 wild-type embryo samples (Cao et al., 2019). This weak interaction is likely caused by postponed recovery of attraction-related proteins (e.g., HMR-1) during cytokinesis, which is distributed on cell membrane and has a time delay when the membrane grows and ingresses (Figure S4). Besides, the membrane segregation of a cell has very little impact on its contacted neighbors, namely, the shared membrane interface seems to be intact. A notable example is that the HMR-1 accumulation in those unperturbed interfaces is preserved during cytokinesis, but there is nearly no accumulation in the membranes between newborn sister cells, supporting the assumed attraction asymmetry between non-sister cells and between sister cells (Figure S4).

Apart from the autogenic cell-cell attraction matrix default above, extra value assignment on the attraction coefficient *σ* between specific cell pair is permitted (hereafter referred to as “attraction motif”), in consideration of other programmed active regulations such as the persistently low accumulation of HMR-1 in EMS-P2 contact. In principle, if the real system needs to be reconstructed more strictly, one can insert both membrane marker and lineaging marker into the HMR-1::GFP strain to facilitate identification of cell morphology and cell position simultaneously, and then quantify the spatio-temporal localization of HMR-1 or other proteins for more accurate distribution profiling and value assignment.

### All the Conserved Cell-Cell Contacts and Non-Contacts from 4- to 8-Cell Stages can be Established under Proper Cell-Cell Attraction Matrix

In combination of both the minimal model and the cell-cell attraction matrix, cell motions from 4- to 8-cell stages are simulated according to division order and division orientation (i.e., volume segregation direction and ratio) measured by experiments (Figures 3A–3C, 4A, S5, S6, S7A and S7B; Table S2; Movies S5-S8). The cell division is initiated at the time point when the motional embryo reaches a quasi-steady state with temporally minimal kinetic energy, i.e., when the embryo moves with the slowest velocity. It should be pointing out that, in simulation, both 6- and 7-cell stages end at their first quasi-steady state and the 8-cell stage ends at its second quasi-steady state, for the reason that the duration of 8-cell stage is over twice of those of 6- and 7-cell stages in real embryo (Guan et al., 2019). To establish the ground truth for verification of the simulation results, we describe the embryo morphology with the cell-cell contact relationships reproducible among individual. A cell pair is defined as “conserved contact” for a specific stage if it exists in all the 4 embryo samples (*N*_contact_ = 4). If a cell pair doesn’t contact in any of them, it’s defined as “conserved non-contact” (*N*_contact_ = 0), then the other cases are “unconserved contact” (1 ≤ *N*_contact_ ≤ 3) (Tables S4-S6). These three situations make up a cell-cell contact map with potential information that which contact/non-contact is significant and which one matters little. In the following comparisons between simulation and experiment, we only focus on the conserved contacts and non-contacts which are assumed to be important properties for a developing *C. elegans* embryo.

**Figure 3.**
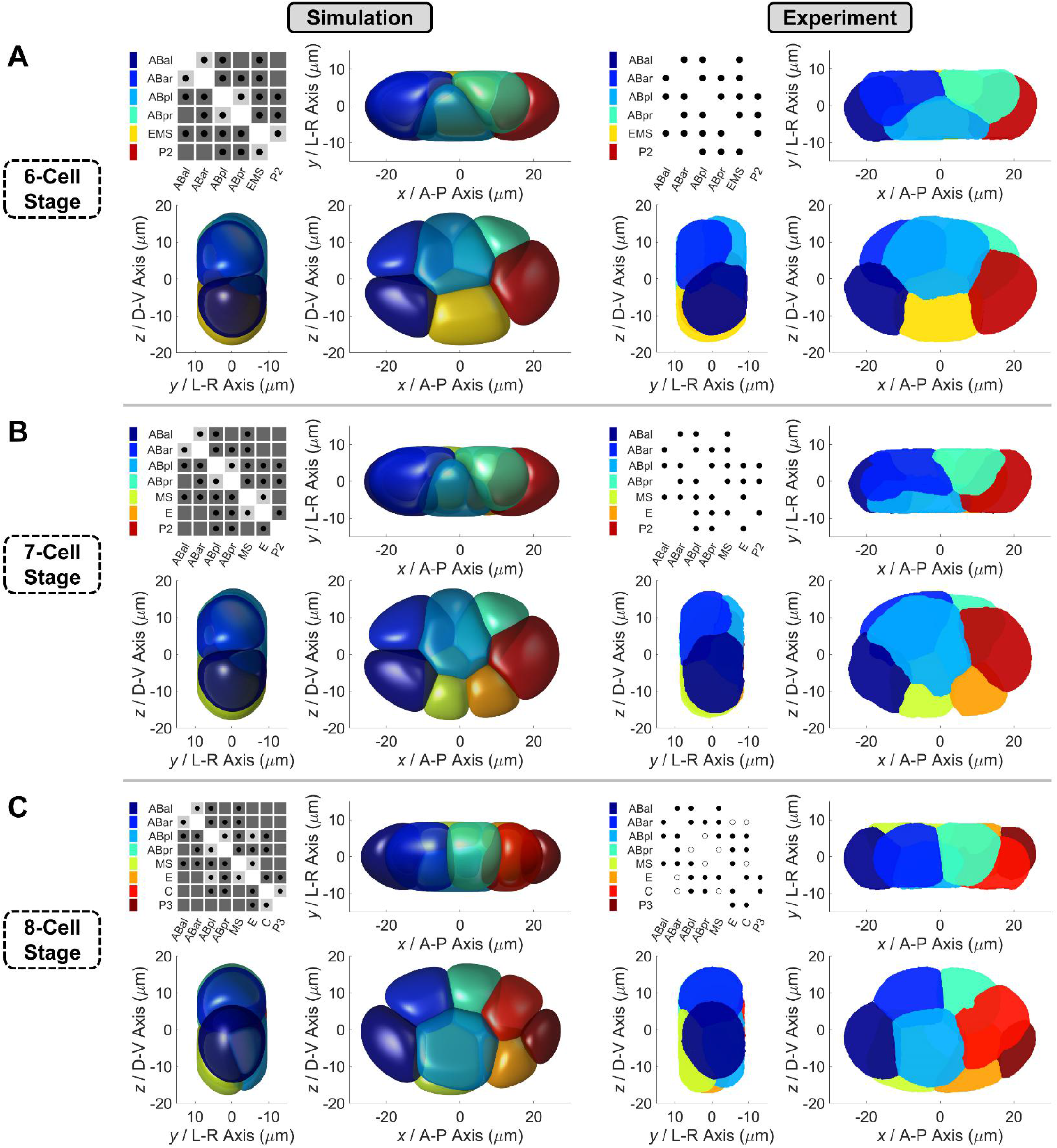
Comparison of Compressed Embryo Morphology between Simulation and Experiment from 6- to 8-Cell Stages. (A) The upper panel, 6-cell structure. (B) The middle panel, 7-cell structure. (C) The lower panel, 8-cell structure (with attraction motif on ABpl-E contact, i.e., *σ*’_ABpl-E_ = 0.2). In each panel, embryo morphology in simulation and experiment is respectively illustrated on the left and right in three orthogonal observation directions, while a cell-cell contact map is placed in the top left conners. About the map in simulation, dark gray and light gray shades denote strong attraction (*σ* =0.9) and weak attraction (*σ*’ = 0.2) respectively, while black dots represent the contacted cell pairs. About the map in experiment, black dots represent the conserved contacted cell pairs which are reproducible in all the 4 embryo samples, while empty circles represent the unconserved contacted cell pairs which exist in at least one embryo sample, but not all of them (Cao et al., 2019). The relationship between cell identity and color is listed next to the contact maps.

**Figure 4.**
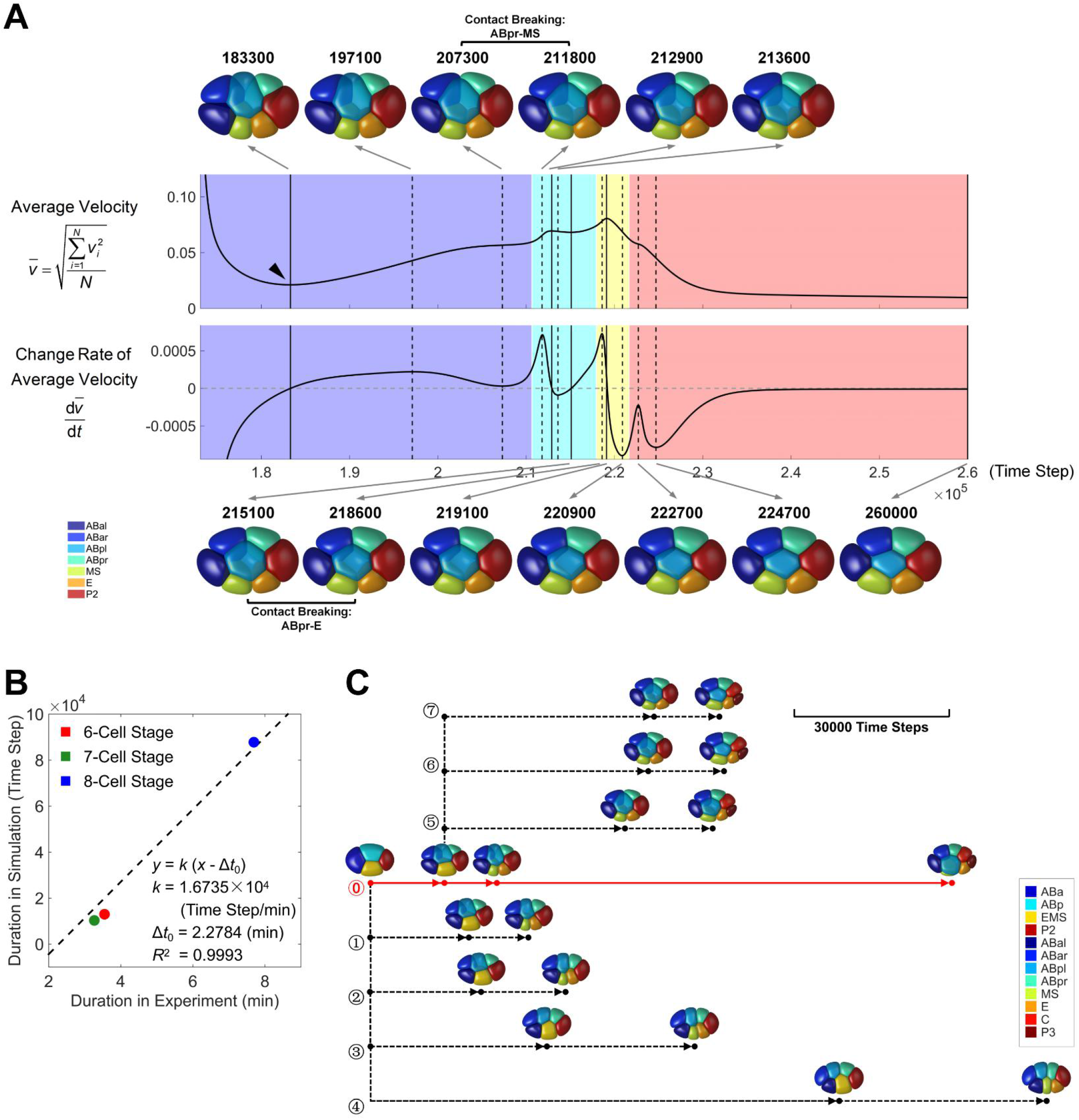
The Role of Timely Cell Division to Protect the 3D Embryo Structure and Cell-Cell Contact Map. (A) Morphological evolution of a compressed embryo during 7-cell stage, with simulation time long enough for the whole system to reach mechanical equilibrium. The curves of average velocity (upper) and its change rate (lower) are illustrated side by side. The solid and dashed vertical black lines denote the extreme points in the curves of average velocity and its change rate, respectively. The time point of the first quasi-steady state is indicated by a black triangle. The 3D structures at those time points in both curves are illustrated on top and bottom, pointed by gray arrows originating from their corresponding lines. The last structure in the bottom right is the system’s terminal state approaching mechanical equilibrium. The change of cell-cell contact map is illustrated by different colors in the background, while the detail is written between two consecutive structures. The relationship between cell identity and color is listed in the bottom left corner. (B) Linear fitting of time scale between simulation and experiment systems. The durations in simulation are obtained from the quasi-steady states (Figures 4A, S6 and S7B), while the ones in experiment are obtained from 222 wild-type embryo samples in previous dataset (Guan et al., 2019). The intercept is predetermined as – Δ*t*_0_ =– 2.2784 min, obtained from 4 wild-type embryo samples in previous dataset (Cao et al., 2019). (C) A tree composed of developmental paths diversified by different cell division timing. The branch of normal developmental path is plotted with red solid line while the ones with disturbed cell division timing are plotted with dashed black lines. The cell division timing is denoted with solid point. The perturbed cell division timing is set at the critical time points (i.e., extreme points) in the curves of average velocity as well as its change rate (Figures 4A and S6), and only the ones with developmental path differentiated from the others are plotted. The final state is determined by the first and second quasi-steady state for 7- and 8-cell stage, respectively. The 3D structures at the time points with cell divisions activated are illustrated near the corresponding points. A scale bar representing *in silico* developmental time is placed in the top right conner. The relationship between cell identity and color is listed in the bottom right corner. The terminal embryo morphology and cell-cell contact map of branches **⓪** ~ ⑦ can be found in Figure S9.

For both 6- and 7-cell stages, the model and assumptions on cell-cell attraction matrix (default) straightforwardly reproduce all the cell-cell contacts and non-contacts conserved in the 4 embryo samples. Apart, cell location and cell morphology in simulation and experiment are highly similar to each other, indicating that the simulation has successfully recaptured the physical state of each cell as well as the mechanical interactions between them (Figures 3A and 3B; Movies S5 and S6). The ring-like structure at 7-cell stage, which is formed by 6 cells contacting and surrounding ABpl, was recently reported to play a decisive role in polarity redistribution and axis establishment (Figure 3B; Dutta et al., 2019). Our simulation suggests that this characteristic pattern is set up by cooperative cell division orientations of ABa, ABp and EMS, which together make ABpl the geometric center of the whole embryo. This may explain why ABpl has so unique motive behavior compared to the others, including the most severe deformation and the longest travel distance (Pohl and Bao, 2010; Guan et al., 2019).

Even though the cell location, cell morphology and cell-cell contact relationship are reproduced up to 7-cell stage, the simulation fails in the upcoming 8-cell stage when the self-determined attraction matrix is applied. Although the initial 8-cell structure resembles the real one, ABpl would ingress inward the embryo along L-R axis instead of migrating in anterior-ventral direction, resulting in a flattened structure with the conserved ABpr-MS contact broken immediately, which is twodimensional in AP-DV plane eventually (Figures S5 and S7A; Movie S7). A recent work reported the discovery of significantly higher E-cadherin HMR-1 accumulation in ABpl’s anterior contacts than those in posterior, and whether this asymmetric enrichment contributes to ABpl’s directional migration is yet to be answered (Dutta et al., 2019). Regarding the attraction matrix default, all the three cells anterior to ABpl have strong attraction (i.e., *σ*_ABal-ABpl_ = 0.9, *σ*_ABar-ABpl_ = 0.9, *σ*_ABpl-MS_ = 0.9), however, the contacts with posterior cells E and C are also strong while the one with ABpr is weak (i.e., *σ*_ABpl-E_ = 0.9, *σ*_ABpl-C_ = 0.9, *σ*’_ABpl-ABpr_ = 0.2) (Figure S5). Here, we firstly take the attraction motif into account, which symbolizes a change between strong and weak attraction to modify the binarized cell-cell attraction matrix. With the knowledge of anterior-posterior adhesion asymmetry on ABpl, there are two kinds of single motifs that may need to be added, namely, *σ*’_ABpl-E_ = 0.2 and *σ*’_ABpl-C_ = 0.2. Hence, we revise the cell-cell attraction matrix with these two motifs and perform simulation respectively. Surprisingly, the remarkable movement of ABpl as well as a persistent 3D embryo structure are completely rescued by the ABpl-E motif, but not by the ABpl-C motif, indicating that the relatively low accumulation of E-cadherin HMR-1 or weak attraction in posterior, especially in ABpl-E interface, is significant for *C. elegans* morphogenesis at 8-cell stage (Figures 3C and S7B; Movie S8).

After weakening the attraction in ABpl-E contact, ABpl could proceed a continuous long-range movement in anterior and ventral directions, matching the observation *in vivo*. In this case, ABpl acquires more asymmetric forces from outside as E only exerts slight pulling force to it (i.e., *σ*’_ABpl-E_ = 0.2). Fascinatingly, the positions and contacts of the 8 cells exhibit outstanding mirror symmetry about the AP-DV plane, with respect to both the geometric and mechanical topologies. Briefly, ABal, ABar, ABpr, C, P3 and E cells are sequentially located near the AP-DV plane, while ABpl and MS are evenly distributed on both sides of the AP-DV plane (Figure 3C). This spatial symmetry contributes to the structural stability, therefore, ABpl as well as its force field play a pivotal role in early morphogenesis.

It’s worth pointing out that, the binarized asymmetric attraction matrix itself appears to help preserve the established cell-cell contacts and leave longer time for each stage to stabilize, compared with the globally strong (*σ* = 0.9, *σ*’ = 0.9) and weak (*σ* = 0.2, *σ*’ = 0.2) attraction matrices (Figure S8). It’s possible that an asymmetric attraction matrix can improve both compactness and flexibility of cell population, which may be a natural way to build up a robust multicellular structure and utilized by embryos undergoing rapid proliferation. Qualitatively, the embryo with globally weak attraction lacks of internal force to link the cells together (e.g., breaking of ABar-MS contact), while the one with globally strong attraction may have intensive dragging force at some locations (e.g., breaking of ABar-ABpr contact), both consequently generating openings between cells (Figure S8).

### The Cell Division Timing Controlled by Quasi-Steady State is in Line with Experimental Measurements

In simulations for all the three stages (i.e., 6-, 7- and 8-cell stages), we design an autonomic time-selecting rule that cell division should be initiated when the embryo reaches a quasi-steady state with temporally minimal kinetic energy (Figures 4A, S6, S7A and S7B). We bring up this because 1) theoretically, activating cell division at the least motional state can minimize the structural variation raised by intrinsic noise in cell division timing, for example, the duration of 8-cell stage is 7.69 ± 0.93 min *in vivo*; 2) according to simulations for 6- and 7-cell stages, the embryo may collapse into defective flattened structure over time, which has lower system energy and higher structural stability (Figures 4A and S6); 3) the simulated cell morphologies accord well with the experimental observations, when cell division timing is selected by quasisteady state (Figures 3A–3C). To validate whether our simulation could represent the real situation with respect to time scale, we align the time scale of our model (time step) to the one in reality (min). To this end, we perform proportional linear fitting on the durations of all 6-, 7- and 8-cell stages between model and experiment (least square method), revealing conversion ratio *k* = 1.6735 × 10^4^ time step/min and considerable goodness of fit *R*^2^ = 0.9993 (Figure 4B). An intercept is predetermined as – Δ*t*_0_ = – 2.2784 min, which is the experimental duration between cell nucleus separation and cell membrane separation, for the reason that the process of elongation as well as cytokinesis during cell division is ignored in our simulation; instead, we split the two daughter cells into individuals and let them interact with each other instantly when executing cell division. The genetically programmed length relation Δ*t*_7–Cell_ < Δ*t*_6-Cell_ < Δ*t*_8–Cell_ is also recaptured by this time-selecting rule. Sufficient time is required for 8-cell stage compared to those of 6- and 7-cell stages, allowing the long-range migration of ABpl as well as the global rotation of embryo. In conclusion, the autonomic time-selecting rule based on quasi-steady state captures the cell division timing *in vivo* and the phase field model is capable of characterizing the morphogenesis in a live embryo, from both perspectives of space and time.

In nematode species like *C. elegans* and *C. briggsae*, cell divisions before gastrulation are coordinated in an invariant sequence, separating the developmental procedure into stages with specific cell numbers (Zhao et al., 2008; Memar et al., 2019). For each stage, the multicellular structure starts from a group of cell division(s) and ends in another. Induced by shape-changing during cytokinesis, the static state of an embryo would be intensively perturbed at first, and then relaxes and stabilizes into an approximately close-packing structure under mechanical interactions, which was previously found in both physical simulations and experimental observations (Fickentscher et al., 2013; Fickentscher et al., 2016; Guan et al., 2019). The next cell division(s) needs to be activated in a steady and noise-limited environment, so as to confirm the precision and robustness of spatial development. The motional intensity will transform the intrinsic noise in cell division timing into the spatial noise in cell-arrangement progression, leading to unavoidable structural variability at the moment when cell division(s) occurs. This may explain why the cell division timing in early *C. elegans* embryo is elaborately programmed in an ordered sequence and matches the one predicted with quasi-steady state.

Given that the time scale *in silico* can represent the one *in vivo*, next we explore how different physical factors affect the developmental path of embryonic morphogenesis. We only focus on 4- to 8-cell stages for that the embryo evolves from 2D to 3D structure and establishes its body axes during those stages, which is also well reproduced by the intercellular cytomechanics set up in our computational system (Figures 3A–3C). Here, we select the three key factors inputted into the phase field model, i.e., cell division timing, cell division orientation, cell-cell attraction matrix, and seek to uncover their roles for *C. elegans* spatial development.

### A Timely Cell Division Can Protect the Established 3D Embryo Structure and Cell-Cell Contact Map

In *C. elegans* early embryogenesis, various underlying mechanisms have been identified to coordinate cell division timings, including asymmetric cell division (Brauchle et al., 2003; Benkemoun et al., 2014; Guan et al., 2020), cell volume reduction (Arata et al. 2015; Fickentscher et al., 2018), cell-cell signaling (Rocheleau et al., 1997; Boeck et al., 2011) and so forth, which maintain the orders and durations between consecutive division events. Previous studies suggest that the cell division timing has a non-ignorable effect on how cells migrate and how an embryo evolves, by simulating the system with coarse-grained model and perturbed cell division timing (Fickentscher et al., 2016; Tian et al., 2020). In our model, we hypothesize that cell divisions are activated when the moving embryo reaches a quasi-steady state with the overall slowest velocity. This hypothesis is reasonable not only because the derived cell division timing is close to the ones *in vivo*, but also for the comprehensive reconstruction of embryo morphology. Notably, a trend of “planarization” in embryo structure is found in simulations for both 6- and 7-cell stages when cell division timing is postponed, in which a lot of cell-cell contacts are broken and all the cells are located within AP-DV plane in the end (Figures 4A and S6). An accurate control on cell division timing seems to be necessary for protecting the system from losing the functional 3D structure as well as its cellcell contact map. We notice that, for 6- to 8-cell stages, all the conserved cell-cell contacts and non-contacts are initially established by the directed cell divisions and persistently remained until the divisions of those cells, no matter in simulation or in experiment. As previous studies proposed that the early embryo can achieve accurate cell migrations and cell-cell contacts merely by basic mechanical coordination (Fickentscher et al., 2016; Tian et al., 2020), setting timely cell divisions may be a strategy/principle to provide stable and continuous contacts to establish cell-cell communications with active regulations as least as possible, which are valuable resource for multiple biological processes including fate specification (e.g., Notch signaling from MS to ABalp and ABara) (Priess, 2005; Neves and Priess, 2005) and division orientation (e.g., Wnt signaling from P2 to EMS and from C to ABar) (Thorpe et al., 1997; Walston et al., 2004).

To further validate this idea and search the developmental paths diversified by cell division timing, we disturb the division timing of EMS and P2, and observe what will happen in the later morphogenesis. We independently initiate cell division at the critical time points (i.e., extreme points) in the curves of average velocity as well as its change rate at 6- and 7-cell stages (Figures 4A and S6), and then push forward the structural evolution using completely same cell division orientation and cell-cell attraction matrix. A tree composed of 8 developmental paths as branches are identified based on their distinguishable final cell-cell contact maps (Figures 4C and S9). Interestingly, if EMS divides a little bit later (≥ 5500 time steps) than the time point of the first quasi-steady state, the ABar-ABpr contact conserved in real embryos would be broken immediately at 7-cell stage. For the excessively delayed timing in P2 division (≥ 24000 time steps), although the ABpl-E motif is remained, the terminal embryo structure fails to be three-dimensional. In summary, an opportune cell division timing is critical for maintaining both the 3D structure and cell-cell contact map of a compressed embryo.

### Both Volume Segregation Direction and Ratio during Cytokinesis have Selective Impact on the Developmental Path

Here, we emphasize the definition of “cell division orientation” as a combination of direction and ratio of cell volume segregation during cytokinesis. By molecular and genetic approaches, cell division orientation has been found to be regulated by redirected myosin flow (Naganathan et al., 2014; Sugioka and Bowerman, 2018; Pimpale et al., 2020) and cellcell signaling (Thorpe et al., 1997; Walston et al., 2004; Langenhan et al., 2009), where cell-cell contact serves as a common basis for both mechanisms. To investigate how cell division orientation is associated with the selection of developmental path, we first align the division orientations of ABa, ABp, EMS and P2 onto the three orthogonal body axes respectively and perform simulations, constructing an even more simplified scenario based on geometry instead of merely experimental measurement (Figure 5A). The dominant orientation of ABa and ABp divisions should be parallel to D-V and L-R axes respectively, so that the four AB descendents can relax inside the compressed eggshell and establish a tetrahedron-like compact cell-cell contact map. As expected, the division orientation of EMS should be along A-P axis, owing to the Wnt signaling from P2 in posterior (Thorpe et al., 1997). It’s worth noting that all the divisions of ABa, ABp and EMS are almost symmetric in volume segregation (Table S2), and only the trials with dominant directions mentioned above can rebuild the embryo morphology and cell-cell contact map resembling the experimental ones. On the other side, for the asymmetric division of P2, C acquires a volume around twice of that of P3 (Table S2). Briefly, there are a total of 3 × 3 = 9 possible combinations of volume segregation direction and ratio (i.e., direction along A-P, L-R or D-V axis, *V*_C_: *V*_P3_ = 2:1, 1:1 or 1:2). It’s surprised that the volume segregation direction of P3 has to be along D-V axis and a larger cell (C) has to be placed on top, otherwise an allosteric 3D structure will be generated (Figure 5A). This observation suggests that, the spatial development at 8-cell stage is highly sensitive to P2 division, which may serve as a structural regulator by special volume segregation direction and ratio.

**Figure 5.**
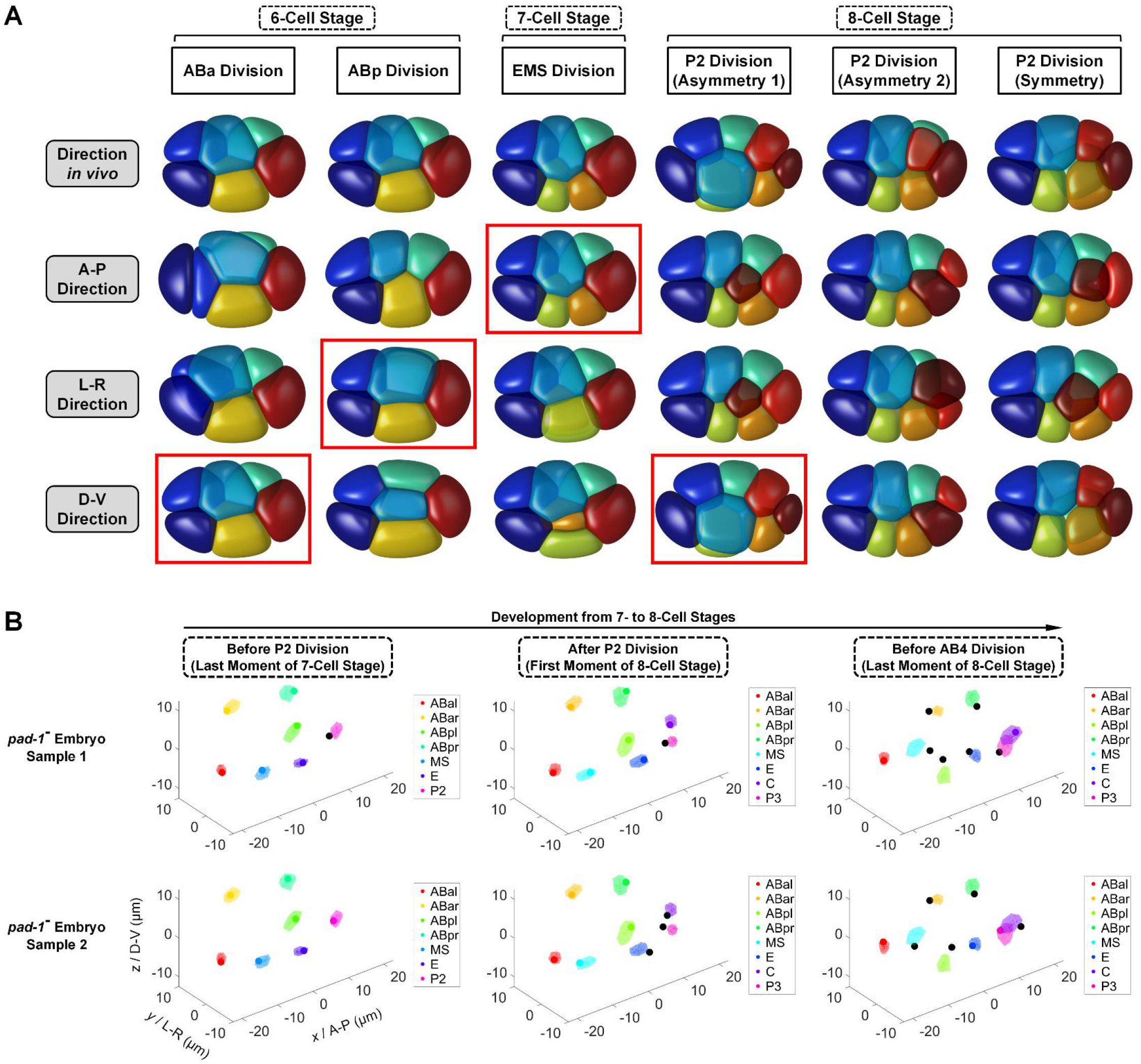
The Impact of Cell Division Orientation on Selecting the Developmental Path. (A) Simulation for 6- to 8-cell stages with cell division orientations aligned to the three orthogonal body axes. All the division orientations of ABa (1^st^ column), ABp (2^nd^ column), EMS (3^rd^ column) and P2 (4^th^, 5^th^, 6^th^ columns with different volume segregation ratio) are set along the experimental (1^st^ row), anterior-posterior (*x* I A-P; 2^nd^ row), leftright (*y* / L-R; 3^rd^ row) and dorsal-ventral (*z* / D-V; 4^th^ row) directions, respectively. The 3D structures whose cell-cell contact map is the same as the one simulated with experimental orientation, are highlighted with red rectangles. (B) Cell-arrangement progression of two *pad-1* RNAi-treated embryos from 7- to 8-cell stages, revealing reproducible positional defects in P2 division orientation and consequent global positional misarrangement. Each color represents one specific cell identity, denoted in the legend on right; the misarranged cells in RNAi-treated embryos are illustrated with black points, while the normal cells are illustrated with original color according to the legend. For each cell, a region formed by nuclei positions from 222×0.95 ≈ 210 independent wild-type embryos is illustrated for visual comparison. Data of both wild-type and RNAi-treated embryos is obtained from a previously established dataset (Guan et al., 2019).

To verify this hypothesis and uncover the possible underlying mechanism specifically controlling P2 division, we screen 1818 RNAi-treated embryos (758 genes, at least 2 replicates for each) with a 4D morphogenesis reference, which is formed by nuclei positions of 222 *C. elegans* wild-type embryos and provided in our previous work (Guan et al., 2019). Notably, we identify a gene, *pad-1* (abbreviation of <u>pa</u>tterning <u>d</u>efective <u>1</u>), whose corresponding RNAi-treated embryos have normal structure at the last moment of 7-cell stage but have significantly deviated division orientation in P2, eventually resulting in a global positional defect as soon as the end of 8-cell stage (*n* = 2, confidence degree = 0.99; Figure 5B). Previous research proposed that silence of *pad-1* would lead to failed embryonic patterning since gastrulation (onset from 26- to 28-cell stages), which however, has little impact on cell fate specification (Guipponi et al., 2000). Here, we bring forward the initial defective timing of *pad-1* RNAi-treated embryos to 7- to 8-cell stages, by detecting cellular misposition with nucleus-based statistical tools established before. We suggest that this defect is likely due to the erroneous division orientation in P2, which immediately leads to a significant global misarrangement in cell positions at the end of 8-cell stage. The detailed function of *pad-1* is elusive and worth of investigation by *in vivo* experiments, especially how it controls P2 division and leads to morphogenetic chaos when perturbation is performed. To summarize, the simulation and gene screening results suggest the impact of both volume segregation direction and ratio on selecting the developmental path of embryo morphology.

### The Cell-Cell Attraction Matrix Provides High-Dimensional Diversity and Regulatory Potential for the Developmental Paths

Cell-cell attraction has been proposed to coordinate cell patterning during early embryogenesis (Nance et al., 2003; Yamamoto and Kimura, 2017; Guan et al., 2020). As some membrane-based adhesive molecules like HMR-1 have a delay in recovery during cytokinesis, we approximate cell-cell attraction with an autonomic binary matrix, which is summarized as relatively strong attraction between non-sister cells and weak attraction between sisters. Nevertheless, as a general mechanical concept of intercellular interaction, this attraction can also be achieved by other biophysical components (e.g., gap junction and filopodial protrusion) (Starich et al., 2003; Altun et al., 2009; Pohl and Bao, 2010) and regulated by some fundamental activities (e.g., cell polarization and signaling pathway) (Nance et al., 2003; Armenti and Nance, 2012; Klompstra et al., 2015). Considering that the simulation without any attraction motif can directly reconstruct the structural evolution of 6- and 7-cell stages but not the 8-cell stage, which however, can be reproduced when single motif on ABpl-E contact is added (Figures 3A–3C, 4A, S5, S6 and S7A-S7B), we choose the 8-cell stage as the object for further research. Here, for simplicity, we only study the effect of single motif added on the attraction matrix default. All the 17 kinds of motif between two specific cells contacted are perturbed one by one separately, from weak to strong (*σ*’ = 0.2 → *σ* = 0.9) or on the contrary (*σ* = 0.9 → *σ*’ = 0.2). All the simulations are first run until time point 350000 (Figure S10A), then the ones with obvious 3D structures are extended to time point 450000 (i.e., ABpl-MS, ABpl-E and C-E motifs) (Figure S10B). The evolutionary sequence of cell-cell contact map is used to classify the developmental paths. The simulations reveal 11 different types of developmental path, 5 different types of terminal structure and 26 different types of cell-cell contact map (Topologies 1 ~ 26 in Figures 6A, 6B and S11). All the paths are first bifurcated from the original 8-cell structure established by P2 division (Topology 1 in Figures 6A and S11), even though the ABar-ABpr motif could transiently destablize their contact, which is broken and recovered within 12200 time steps (Topology 21 in Figures 6A and S11). Interestingly, some paths can merge with or separate from each other, for instance, the terminal structure of attraction matrix default can be reached by the ones with ABal-ABar, ABal-ABpl, ABal-MS, ABar-ABpl or ABpr-C motif (Topology 4 in Figures 6A and S11), through different evolutionary sequences.

**Figure 6.**
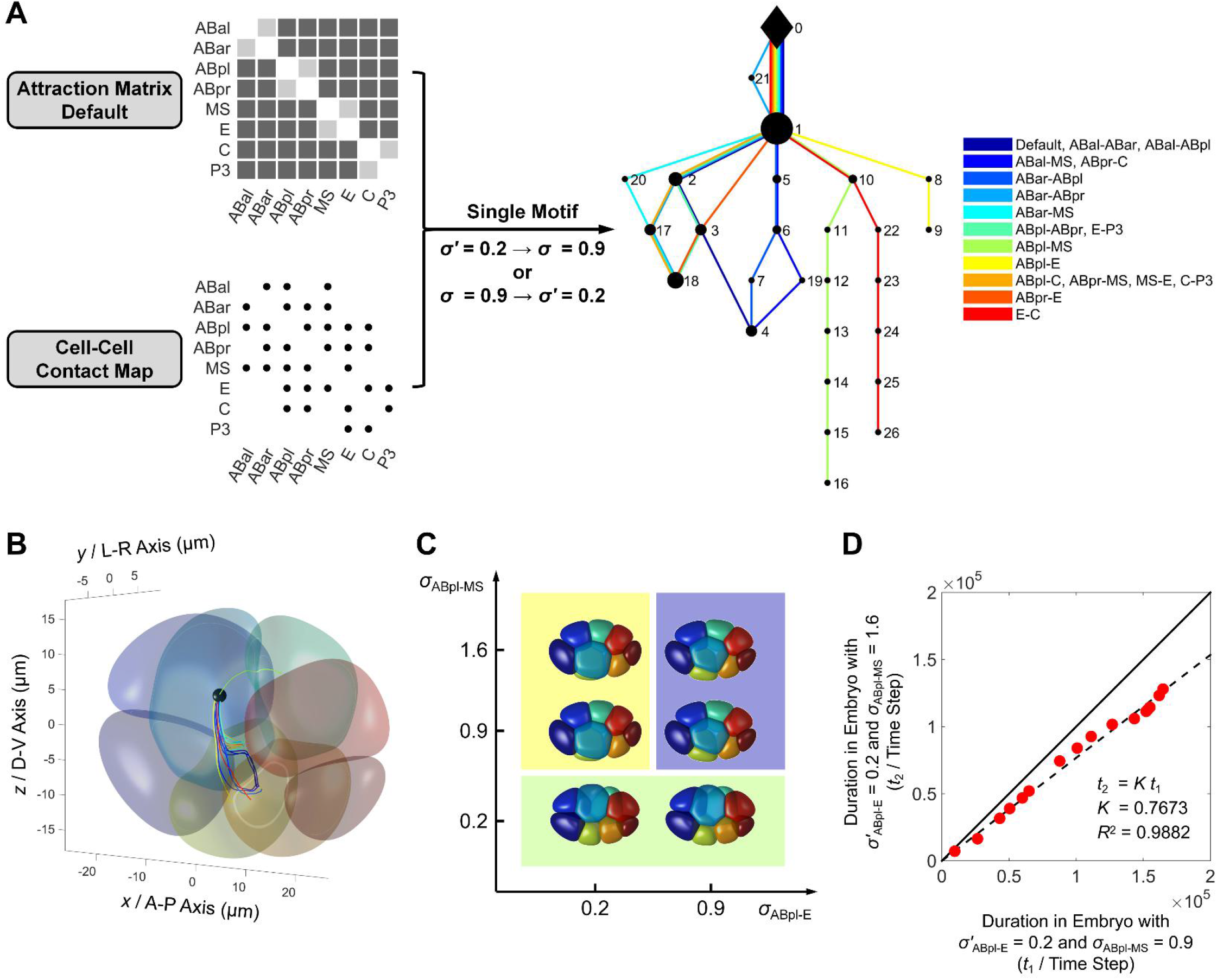
High-Dimensional Diversity and Regulatory Potential of Developmental Paths Provided by Cell-Cell Attraction Matrix and Extra Motifs. (A) A total of 17 single motifs (*σ*’ = 0.2 → *σ* = 0.9 or *σ* = 0.9 → *σ*’ = 0.2) based on the cell-cell contact map at 8-cell stage (bottom left corner) are applied on the cell-cell attraction matrix default (top left corner) sequentially for simulation. The evolutionary sequence of cell-cell contact map is used to classify the developmental paths, illustrated on the right. Each color represents a unique developmental path originating from top and ending in bottom, as indicated in legends; the black diamond denotes the cell-cell contact map at the last moment of 7-cell stage (Topology 0 in Figure S11); the black points denote the different cell-cell contact maps at 8-cell stage (Topologies 1 ~ 26 in Figure S11), with size positively correlated to the number of developmental paths passing through. (B) The migration trajectory of ABpl’s mass center when the 17 single motifs are added sequentially. The initial 8-cell structure is illustrated semi-transparently for visual comparison; the colors representing different cell identities are the same as those used in Figure 3C. The color representing trajectories with different motifs are the same as those used in Figure 6A. (C) The three types of developmental paths differentiated when different combinations of *σ*_ABpl-E_ and *σ*_ABpl-MS_ are applied (*σ*_ABpl-E_ = 0.2, 0.9 and *σ*_ABpl-MS_ = 0.2, 0.9, 1.6), highlighted with light yellow, purple and green background. The 3D structures at their second quasi-steady state are illustrated. (D) Match of migration rate between embryos with *σ*’_ABpl-E_ = 0.2, *σ*_ABpl-MS_ = 0.9 and *σ*’_ABpl-E_ = 0.2, *σ*_ABpl-MS_ = 1.6. A total of 16 critical time points selected by extreme points of the curves of average velocity and its change rate are used to compare the migration rate in the two embryos (Figure S12; Table S10).

### Low Attraction in ABpl-E and High Attraction in ABpl-MS Serve as Stabilizer and Accelerator Respectively

In the scanning of attraction motif, there are 5 different types of terminal structures including two failing to reach threedimensional (Topologies 4 and 18 in Figures 6A and S10A) and three succeeding (Topologies 9, 16 and 26 in Figures 6A and S10A), with respect to their structures at time point 350000 (duration ≈ 9.96 min *in vivo*). Among them, the ABpl-E motif protects the 3D embryo structure most constantly, which has the longest interval with unchanged cell-cell contact map (duration = 150500 time steps *in silico* ≈ 8.99 min *in vivo*) (Table S9), making it an excellent stabilizer for both the 3D embryo structure and cell-cell contact map.

Apart from the 3D structures driven by ABpl-MS, ABpl-E and C-E motifs, there are 5 motifs leading to the same 2D structure as the attraction matrix default, which exhibits mirror symmetry about LR-DV plane (Figures 6A and S10A). Therefore, those motifs are supposed to be hardly essential for cell-arrangement progression at 8-cell stage. The remaining 9 motifs generate another flattened structure with mirror symmetry about AP-LR plane. It shows that the geometrical and mechanical structures of embryo always tend to a terminal state with spatial symmetry, and the one with ABpl-E motif coincidently acquires considerable symmetry about AP-DV plane so that it becomes robust against external compression along L-R axis (Figure 3C).

Previous study discovered that the anteroventral surface of ABpl consists of many protrusions, making a strong bonding and dragging to the MS cell below (Pohl and Bao, 2010). This actively strengthened interaction is ignored in the motif scanning above, because the attraction between ABpl and MS is already set to be relatively strong (*σ*_ABpl-MS_ = 0.9). To uncover the potential function of an even more strengthened attraction between them, we try to increase its *σ* value and conduct series of simulations under *σ*_ABpl-E_ = 0.2, 0.9 and *σ*_ABpl-MS_ = 0.2, 0.9, 1.6. Surprisingly, there are three distinct types of developmental paths regarding the 8-cell structures at their second quasi-steady state (Figure 6C). The developmental path *in vivo* can be reproduced only when ABpl-E attraction is weak (*σ*’_ABpl-E_ = 0.2) and ABpl-MS attraction is strong (*σ*_ABpl-MS_ = 0.9 and 1.6). When ABpl-E attraction is stronger (*σ*_ABpl-E_ = 0.9), the developmental path would be the same to the one with attraction matrix default, i.e., leading to a flattened 2D structure. On the other hand, ABpl will stay in the dorsal if ABpl-MS attraction is too weak (*σ*_ABpl-MS_ = 0.2), no matter *σ*ABpl-E = 0.2 or 0.9, indicating that the attraction between ABpl and MS should be over an underlying threshold and is crucial for ABpl to migrate anteroventrally. As long as the ABpl-MS attraction is strong enough to drag ABpl downward, increasing the *σ* value can even accelerate the whole migratory process. Since the dynamics of average velocity and its change rate are almost linearly scalable between embryos with *σ*_ABpl-MS_ = 0.9 and 1.6 (Figure S12; Table S10), we track the extreme points of both the curves and compare the durations cost to reach them. Clearly, the embryo with *σ*_ABpl-MS_ = 1.6 grows around a quarter faster than the one with *σ*_ABpl-MS_ = 0.9, supporting the acceleratory role of ABpl-MS attraction (Figure 6D).

### The Structural Planarization is a Common Defective Phenotype in Compressed Embryo, which could be Induced by Incorrect Cell Division Timing, Cell Division Orientation and Cell-Cell Attraction Matrix

Apparently, a characteristic tendency of structural planarization exists at all the 6-, 7- and 8-cell stages if the system is allowed to relax with enough time (Figures 4A, 4C, S6, S7A, S9, S10A and S10B). According to our simulations, there are three ways to prevent the multicellular system from losing its 3D structure. The first is to initiate cell division(s) when the embryo reaches its quasi-steady state with temporally minimal kinetic energy, which is right before the collapse timing (Figures 4A, 4C, S6, S7A and S9). The second one is to carefully select the cell division orientation, which consists of both volume segregation direction and ratio (Figure 5A). The final one is to add motifs like the weak attraction in ABpl-E contact, so that the 3D structure might be protected persistently by appropriate geometrical and mechanical symmetry (Figures S10A and S10B). Next, we ask if this spatial phenotype is caused by the eggshell compression along L-R axis, where the compression ratio is roughly 0.5 in both simulation and experiment (Figure 1B) (Cao et al., 2019).

To this end, we generate an axisymmetric eggshell with major axis and total volume as same as the ones in compressed embryo. Simulation is carried out with totally same rules and inputs of system parameters, resulting in sustainable stable 3D structures at all the 6-, 7- and 8-cell stages, no matter how long the systems proceed (Figures S13A, S13B, S14, S15, S16A and S16B; Movies S9-S12). The embryo achieves considerable spatial stability and correct cell-cell contact map, which is no more dependent on the cell division timing or the ABpl-E motif (Figure S13B). To further verify the structural planarization predicted by model, we first evaluate the normal embryo width at the last time points of 6-, 7- and 8-cell stages respectively, by calculating the maximum distance in *y* / L-R direction among all the cell nuclei (Figures S17A-S17C). Subsequently, we culture three *C. elegans* wild-type embryos and perform laser ablation onto their P2 cells during 4-cell stage, using methods described before except that the bleaching time is tuned to 90 seconds (Chen et al., 2018). Then, the cell cycle of P2 arrests and the duration of 7-cell stage is lengthened from 3.27 ± 0.87 min to 10.33 ± 0.81 min, while the duration of 6-cell stage remains still. After laser ablation on P2, the embryo width at the last time point of the extended 7-cell stage, but not the 6-cell stage, is significantly narrower than the normal ones (*n*_perturbed_ = 3, *n*_normal_ = 222, *p* ≈ 0.00887, one-tailed Wilcoxon rank-sum test) (Figures S17A and S17B), revealing a structural planarization induced by improper cell division timing. Furthermore, considering that this spatial defect is sensitive to multiple physical factors and commonly exists in systems with all kinds of perturbation, we then look into the RNAi-treated embryos from previous dataset and see which genes can result in this phenotype after interference. A total of 1818 RNAi-treated embryo (758 genes, at least 2 replicates for each) are tested using the same method (Guan et al., 2019). Strikingly, a total of 101 genes were filtered out with significantly narrower embryo width during 6- to 8-cell stages (*n*_perturbed_ ≥ 2, *n*_normal_ = 222, *p* < 0.05, one-tailed Wilcoxon rank-sum test), which are involved in a number of important biological processes including cell polarity (*mex-1, par-1*), cellcycle progression (*cdk-7, chk-2*), Wnt signaling (*kin-19, gsk-3*), Notch signaling (*lag-1, glp-1*) and so on (Tables S11-S13) (Guedes and Priess, 1997; Noatynska and Gotta, 2012; Wallenfang and Seydoux, 2002; Stolz and Bastians, 2010; Walston et al., 2004; Neves and Priess, 2005). Besides, the *pad-1* gene, which has deviated division orientation in P2 after RNAi treatment, would also decrease the embryo width at 8-cell stage (Figure 5B; Table S13). To sum up, the external compression onto embryo can lead to a tendency of structural planarization and lost contacts, which however, can be rescued by proper cell division timing, cell division orientation and cell-cell attraction matrix. It was previously assumed that this lateral compression exists naturally, in particular to the starved or old animals which have very limited and inflexible space for the developing embryos inside them (Jelier et al., 2016). Artificial compression is usually executed by slide mounting in experiments, so that an embryo can be imaged within the pre-determined depth of view field (Murray et al., 2006; Murray et al., 2012; Chen et al., 2018; Cao et al., 2019). Apart from other active mechanisms found to resist mechanical perturbation (Jelier et al., 2016), here we propose the cell division timing, cell division orientation and cell-cell attraction matrix as three key regulatory factors in physical level.

## DISCUSSION

How animal develops from a single zygote into a stereotypic self-organized pattern has been a fascinating problem for decades. Little is known about the fundamental strategies and principles concerning how different dimensions of developmental properties are coordinated in concert to drive the embryogenesis and morphogenesis. In this work, we established a data-driven phase field model to describe the mechanical interactions and constrictions at the cellular level, including cell surface tension, cell-eggshell and cell-cell repulsion, cell-cell attraction and cell volume constriction. In combination of 4D morphological data obtained from live *C. elegans* embryos, we successfully reproduced the structural evolution from 1- to 8-cell stage, in terms of cell location, cell morphology and cell-cell contact relationship. Simulations with variable parameters supported a set of physical factors that are critical for the precise and robust morphogenesis, i.e., cell division timing, cell division orientation and cell-cell attraction matrix. Last but not least, we predicted that a spatially-defective phenotype termed “planarization” would occur easily when a compressed embryo suffers perturbation, such as delayed cell division timing (Figures 4A, 4C, S6, S7A and S9), deviated cell division orientation (Figure 5A) and inappropriate cell-cell attraction matrix (Figures S10A and S10B). This phenotype is validated by both P2-ablated and RNAi-treated embryos (Figure S17; Tables S11-S13).

Accurate control on cell division timing and cell division orientation is crucial to ensure the cell-level precision and robustness during morphogenesis (Fickentscher et al., 2013; Fickentscher et al., 2016; Tian et al., 2020). According to our simulations, the multicellular embryo structure becomes less robust and easy to be planarized when suffering external compression. Consequently, the correct cell-cell contacts and communications cannot be established persistently. Nevertheless, delicate regulations on cell division timing, cell division orientation and cell-cell attraction matrix can rescue the system from losing the target structure. First, the cell division(s) is activated at a quasi-steady state with the slowest global velocity (Figures 4A, 4C, S6, S8 and S9). Second, the division orientation of a cell determines the initial contact relationship between its daughters and the other cells (Figure 5A; Table S2; Movies S13 and S14). Third, the new cell-cell contact map and its effective attraction matrix specify the system’s developmental path (Figures 6A, 6B, S10A and S11). Drawing lessons from “chemoconnectome” for mapping chemical transmission (Deng et al., 2019), here we introduce the concept of “mechanoconnectome” to represent this contact-preserved developmental behavior. The mechanoconnectome is defined as the conserved contact map among specific cells, which keeps evolving and driving embryonic morphogenesis by cell-cell mechanical interactions. This mechanoconnectome makes the spatial development self-organized by the genetically programmed cell division timing, cell division orientation and cell-cell attraction matrix. Consequently, the cell-cell contact map then serves as a foundation for reproducible and robust establishment of functional cell-cell communications. More specifically, our simulations show that different cell-cell contacts have unequal weights in affecting the embryo system. One outstanding example is the ABpl cell and ABpl-E motif at 8-cell stage. Regarding the cell-cell contact numbers, ABpl always contacts with the maximum number of cells (i.e., 5 contacts at 6-cell stage and 6 contacts at 7- and 8-cell stages) (Figures 3A–3C). Since ABpl has this unique geometrical feature, it could play a pivotal role in morphogenesis under regulated cortical actomyosin contractility and filopodial protrusion (Pohl and Bao, 2010). This then provokes a profound question, whether the biological programs are developed or designed to adapt the geometrical and mechanical characteristics? Probably, the *C. elegans* embryo differentially coordinates ABpl with great effort because ABpl is the center of embryo which can lead collective motion much easier than the others.

Combined with the recent progress in data reconstruction on cell morphology and cell position (Cao et al., 2019), we are now capable of building up a data-driven phase field model that is merged, corrected and verified with *in vivo* data for the early development of *C. elegans* embryo. There are still several problems that remain to be solved in the future. First, we simulated the spatial development and searched the solution space by simplification and exhaustion, for instance, in the identification of the 17 single attraction motifs at 8-cell stage. However, as the cell number increases with the *C. elegans* development, it will become computationally too expensive if we want to enumerate all possible stable developmental paths. This problem might be solved by the recently proposed computational approach of constructing the solution landscape to identify all possibilities (Yin et al., 2020). Second, the morphogenesis of a compressed embryo may involve more subtle and complicated regulations such as contraction and polarization when the number of cells is large (Lee et al., 2006; Tao et al., 2020). To simulate more cells in a compressed embryo, it may require extensions of the current minimal phase field model with additional biochemical or biophysical mechanisms.

The proposed phase-field approach in this paper may be applied to reconstruct other multicellular systems, such as *Arabidopsis thaliana* and *Drosophila melanogaster* (Fernandez et al., 2010; Stegmaier et al., 2016). It may also be used in synthetic systems (Cameron et al., 2014). Very recently, a number of multicellular systems were constructed artificially, with acquired practical functions such as self-organization and autonomic motion (Toda et al., 2018; Kriegmana et al., 2020). Theoretical methods capable of describing cell-cell interactions, cell deformation and cell motion are needed. Our generic computational framework could shed light on simulations of *in vivo* multicellular systems and bio-system design.

## STAR*METHODS

- LEAD CONTACT AND MATERIALS AVAILABILITY
- METHOD DETAILS

- Data Collection of Cell-Resolved Developmental Properties from Previous *In Vivo* Imaging Experiments
- Establishment of Computational Framework based on Phase Field Model
- DATA AND CODE AVAILABILITY

## SUPPLEMENTAL INFORMATION

Supplemental Information includes 19 figures, 13 tables and 14 movies.

## AUTHOR CONTRIBUTIONS

L.Z. and C.T. conceived and coordinated the study; X.K. and G.G. performed computation, data analysis and wrote the manuscript; M.K.W. and L.Y.C. performed embryo curation and imaging; Z.Z. provided reagents and experimental methods; L.Z. and C.T. provided theoretical guidance and revised the paper.

## ACKNOWLEDGMENTS

We thank Lei-Han Tang, An-Chang Shi, Xiaojing Yang and Feng Liu for the helpful discussions, and Kai Kang for his assistance in data visualization. This work was partially supported by the National Natural Science Foundation of China (11861130351), the Ministry of Science and Technology of China (2015CB910300), the Royal Society Newton Advanced Fellowship, the Hong Kong Research Grants Council (HKBU12100118, HKBU12100917, and HKBU12123716) and the HKBU Interdisciplinary Research Cluster Fund.

## DECLARATION OF INTERESTS

The authors declare no competing interests.

## STAR*METHODS

### LEAD CONTACT AND MATERIALS AVAILABILITY

Further information and requests for resources and reagents should be directed to and will be fullfilled by the Lead Contact, Lei Zhang.

### METHOD DETAILS

#### Data Collection of Cell-Resolved Developmental Properties from Previous *In Vivo* Imaging Experiments

Here, we collect imaging data as well as its processed results (i.e., cell segmentation, cell tracing, cell lineaging) of three groups of *C. elegans* wild-type embryos from previous datasets, for different purposes of us (Table S1) (Guan et al., 2019; Cao et al., 2019). The first group is a wild-type embryo sample with GFP marker ubiquitously expressed and localized in cell nuclei, which was imaged since 1-cell stage. The second group is formed by 13 samples with only nucleus marker (GFP) imaged since 4-cell stage. The third group consists of 4 samples with both membrane (mCherry) and nucleus (GFP) markers imaged since 4-cell stage, providing extra 3D time-lapse cell morphology information. All the embryos used in this work were cultured in room temperature within 20 ~ 22 °C, imaged using confocal microscope with temporal resolution of ~1.5 minutes, then traced and lineaged using StarryNite and AceTree softwares (Figure 1A) (Murray et al., 2006). The detailed usages of the 18 embryo samples are listed in Table S1.

Using the embryo data obtained above, cell division orientation, cell volume and eggshell shape are quantified and inputted into simulation as predetermined parameter values (Figure 1B; Table S2). Note that as cell morphology is obtained since the last co-existence moments of ABa, ABp, EMS and P2 (i.e., 4-cell stage), the volume of AB is calculated as the sum of volumes of its daughters ABa and ABp, the volume of P1 is calculated as the sum of volumes of its daughters EMS and P2, and the volume of initial zygote P0 is the sum of the inferred volumes of its daughters AB and P1. Cell morphology from 1- to 8-cell stages including cell location, cell shape, cell surface area, and cell-cell contact relationship and area, will be used to test and verify our phase field model (Tables S3-S6). In total, there are 5, 12, 15, 15 conserved contacts and 1, 4, 7, 10 conserved non-contacts (i.e., reproducibly found in all the 4 embryo samples) identified between specific cells at 4-, 6-, 7- and 8-cell stages, respectively. It should be pointing out that the stages with exact cell numbers are achieved by an invariant division sequence in embryos of *C. elegans* as well as its closely-related species like *C. briggsae* (i.e., P0 → AB → P1 → AB2 → EMS → P2 → AB4…), which has been tested in over 200 wild-type embryos (Guan et al., 2019; Zhao et al., 2008; Memar et al., 2019).

#### Establishment of Computational Framework based on Phase Field Model

##### Mechanical Interactions and Constrictions

Phase-field approach, a numerical technique based on a diffuse-interface description, tracks evolution of a phase or species concentration by a set of phase fields instead of explicit and direct tracking of the sharp interfaces. Here, we develop a data-driven phase-field approach to model the morphological and morphogenetic dynamics of a multicellular system. Boundary of the *i*-th cell is represented and tracked by a phase field *ϕ_i_*(*r,t*), *i* = 1, ···, *N* with *ϕ_i_* ∈ [0,1], defined on a computational domain Ω, where *N* denotes the total amount of cells. The interior of cell *i* is *ω_i_* ∈ Ω where *ϕ_i_* = 1, while the complementary domain *ω_i_′* ∈ Ω with *ϕ_i_* = 0 represents exterior of the cell. Thus, the cell membrane is defined as the narrow transition layer in between them. Apart from the cells, an additional phase field *ϕ*_e_ is employed to track the boundary of eggshell and restrict the range of cell movement, where *ϕ*_e_ = 1 represents outside of eggshell and *ϕ*_e_ = 0 represents inside.

The distribution of phase field, namely the deformation and motion of a cell, is determined by a couple of forces imposed including cell surface tension, cell-eggshell and cell-cell repulsion, cell-cell attraction and the homogeneous pressure that constrains cell volume to be constant. These interactions and constrictions can be expressed in both forms of free energy *E* and acting force ***F***.

Firstly, surface energy of the *i*-th cell is defined with Ginzburg-Landau free energy (Lowengrub et al., 2009).

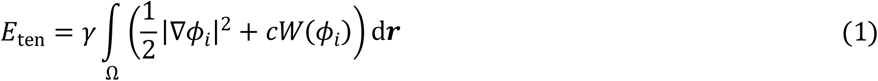

where *γ* is the cell surface tension and *c* is a positive coefficient that determines the thickness of transition layer between interior and exterior of cells; *W*(*ϕ*) = *ϕ*^2^(*ϕ* – 1)^2^ is a double-well potential with minima at *ϕ* = 1 and *ϕ* = 0 which describes the tendency of the phase field to 1 or 0. Minimizing the surface energy *E*_ten_ leads to minimal surface area of a cell, in other words, the cell shape tends to be spherical. A cell with larger *γ* acquires stronger capability to maintain its shape as a sphere. Force field of the *i*-th cell’s surface tension ***F***_ten_ is derived by taking variational derivative on surface energy *E*_ten_ in the form of line density.

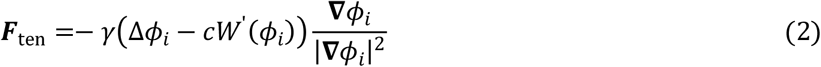

Secondly, cell movements are limited in the interior of eggshell and affected by other cells. Cell-eggshell and cell-cell repulsive potential *E*_rep_ of the *i*-th cell is taken into account and determined as below (Shao et al., 2010).

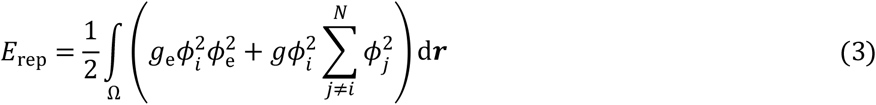

where *g*_e_ and *g* are positive coefficients, denoting the strength of cell-eggshell and cell-cell repulsive energy, respectively. Minimizing the repulsive energy *E*_rep_ can reduce the overlap between phase fields and separate the cells apart. Repulsive force ***F***_rep_ is derived by taking variational derivative with respect to *ϕ_i_* in the form of line density.

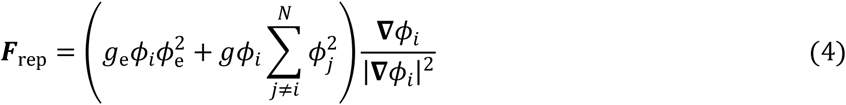

Thirdly, for the possible attraction between cells such as effects of adhesive protein on cell membrane and gap junction between cells (Yamamoto and Kimura, 2017; Costa et al., 1998; Starich et al., 2003; Altun et al., 2009), we model it by introducing an advective item ***F***_atr_ to represent attractive interaction along the normal directions of interface (Löber et al., 2015).

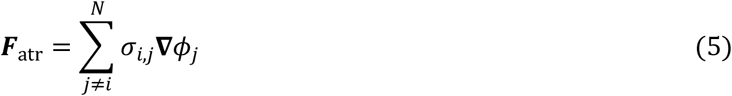

where *σ_i,j_* is a non-negative coefficient and positively associated with the attraction intensity between the *i*-th cell and *j*-th cell.

Fourthly, in an ideal situation, cell volume is constant during cell deformation and motion. Hence, a volume constraint is needed and introduced.

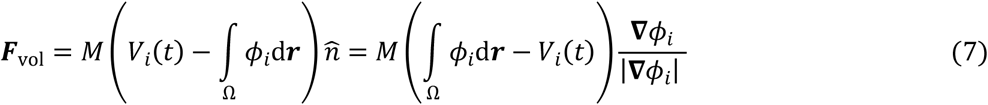

where *M* is a positive coefficient which denotes the volume constraint strength and 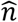 is the unit normal vector at interface which orients inward; *V_i_*(*t*) denotes the prescribed volume for the *i*-th cell, which remains roughly unchanged during cell cycles and updates after cell division since the cell volume is substantially reduced during mitosis.

Inside the overdamped viscous environment of embryo, the resultant of cell surface tension (Equation 2), cell-eggshell and cell-cell repulsion (Equation 4), cell-cell attraction (Equation 5) and cell volume constriction (Equation 7), is always balanced with the viscosity ***f***, which is dominantly determined by a cell’s velocity ***u*** (Fickentscher et al., 2013).

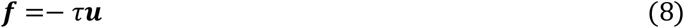

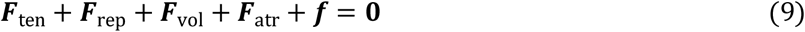

Where *τ* is the viscosity coefficient of the embryo’s inner environment. Finally, evolution of all the phase fields *φ_i_* follows 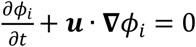, which can be consequently transformed into Equation 10.

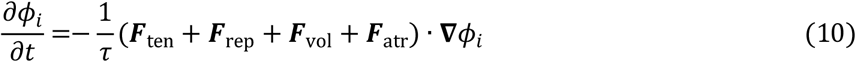

##### Cell Division

We choose a simplified mathematical description for cell division which is implemented as instantaneous bisection of phase field *ϕ_i_* by a plane ***n*** · (***r*** – ***r**_c_*) – *b* = 0. Location and direction of the splitting plane are determined by the cell volume segregation direction and ratio obtained from *in vivo* measurements. Division of the *i*-th cell is processed following Equations 11 and 12 (Akiyama et al., 2018).

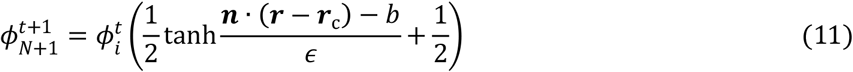

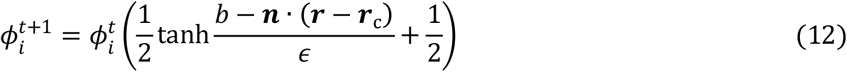

where 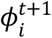 and 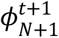 are the phase fields of two daughter cells at their first appearance moment; ***n*** is the unit vector along the division orientation. 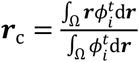 is the mass center of mother cell; *b* is a constant obtained by minimizing 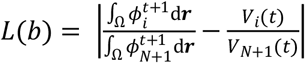, which controls the volume segregation ratio between two daughter cells; *ϵ* controls the width of interface between two daughter cells. It’s worth noting that both the cell division order and cell division orientation is kept in line with the experimental observations when simulating the real scenes (Tables S2-S6), and then we disturb them to discover their roles in embryonic morphogenesis.

##### Parameter Setting

We use a 256 × 256 × 128 rectangular grid as computational domain with grid size d*l* = 0.2508 μm and time step *h* = 0.1. For comparison, the major and minor semi-axes of observed eggshell are 27.5837 μm (*x*) and 18.3477 μm (*z*) respectively. As the eggshell in imaging experiments was compressed about 16.7942 μm narrower in total along the leftright axis (*y*) by slide mounting, we set a cutoff on the left-right axis (*y*) based on experimental values (effective half width = 9.9506 μm) and then construct a compressed eggshell as boundary (Figure 1B) (Cao et al., 2019).

To build up a minimal model which has the least physical constraints but outlines the most significant characteristics of a developing embryo, we complicate the system step by step. We first neglect the intercellular attraction (i.e., *σ_i,j_* = 0), then only three sets of independent physical coefficients are left: cell surface tension (*γ*), transition layer thickness (*c*) and cell- eggshell repulsive energy (*g*_e_). For the other system parameters, we assign constant values as *g* = 1.6 (mechanical unit), *M* = 0.0012, *τ* = 2.62 and *ϵ* = 2^−52^ throughout all the simulations. Sensitivity and comparison analysis are carried out on the physical coefficients *γ, c* and *g*_e_ to determine their optimal values, by evaluation methods as the following.

1. Maximizing the deformation of cell compared with a standard sphere (*α*), according to Equations 13 and 14.

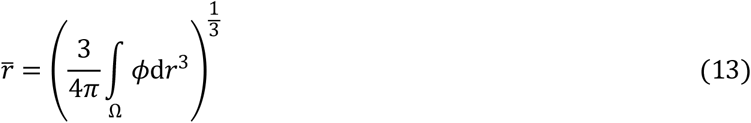

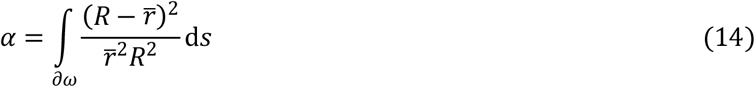

where 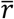 is the average radius of a cell obtained from its equivalent volume; *∂ω* is the cell surface determined by *ϕ* = 0.5; *R* is the distance between panel element ds and the cell’s mass center 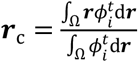.
2. Maximizing the phase-field gradient at interfaces (|∇*φ*|_max_). This requirement sharpens the cell boundary and draws a clear distinction between cells.
3. Minimizing the average positional variation between simulation and experiment, according to Equation 15.

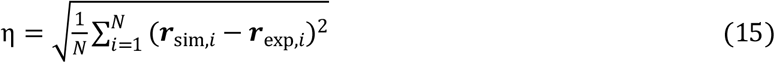

where ***r***_sim,*i*_ and ***r***_exp,*i*_ represent the 3D position of cell *i* in simulation and experiment respectively; *η* is the average positional variation.

These optimization operations result in a parameter combination of *γ* = 0.25, *c* = 1.0 and *g*_e_ = 16, which are used in all the simulations in this work (Figures S18A, S18B and S19). Note that the embryo structure is unsensitive to the exact value of *g*_e_, thus we assume that the stiffness of eggshell is one magnitude larger than that of cell, namely *g*_e_ = 10*g* = 16 (Figure S19) (Guan et al., 2020). Importantly, all the attraction coefficients between cells (*σ_i,j_*) are set as zero at the start and assigned with different values in the later simulations, as a high-dimensional factor to diversify the development paths.

### DATA AND CODE AVAILABILITY

All the data and code used in this paper are available upon reasonable request.

## Notes

### Competing Interest Statement

The authors have declared no competing interest.

### Summary of Updates

Author information, supplemental file and manuscript format are revised.

