## Supplemental Information for "Computable Early *C. elegans* Embryo with a Data-driven Phase Field Model"

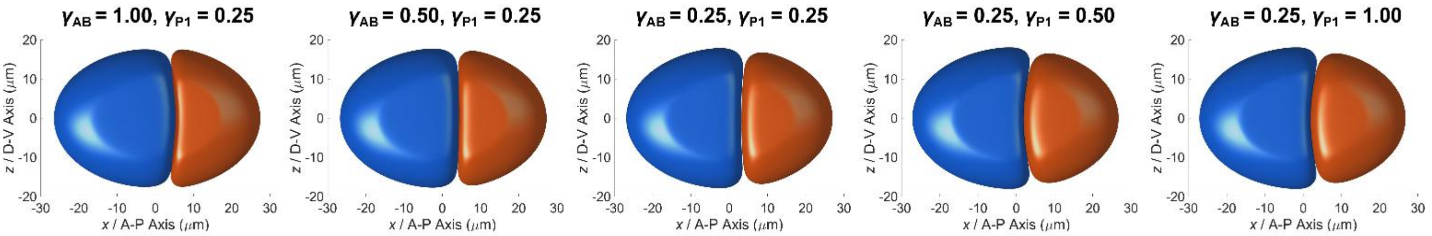


**Figure S1. Relationship between cell surface tension ($\gamma$) and the curvature of interface between cells.** In the 5 simulations from left to right, the ratio between $\gamma_{AB}$ and $\gamma_{P1}$ is changed from 1.00:0.25 to 0.25:1.00; blue cell, AB; orange cell, P1.

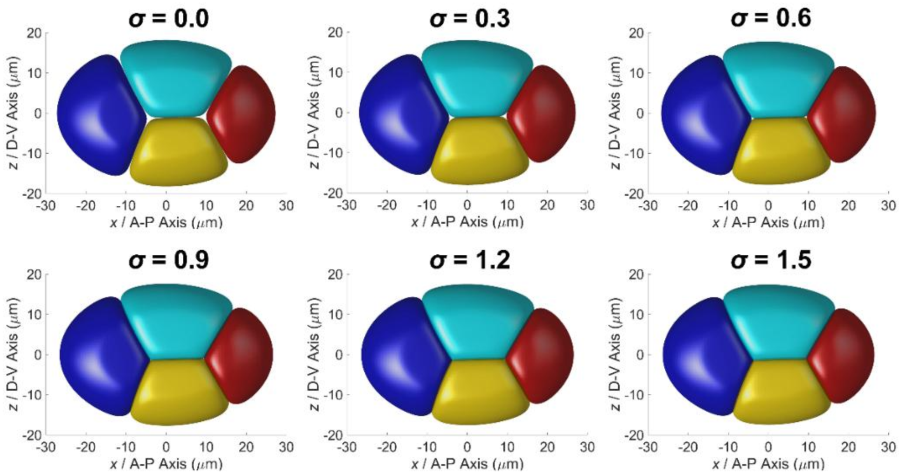


**Figure S2. Relationship between global attraction ($\sigma$) and the hollows at junction points among cells.** In the 6 simulations from top left corner to bottom right corner, $\sigma$ is changed from 0.0 to 1.5, equally applied onto all the 5 cell-cell contacts; blue cell, ABa; cyan cell, ABp; yellow cell, EMS; red cell, P2.

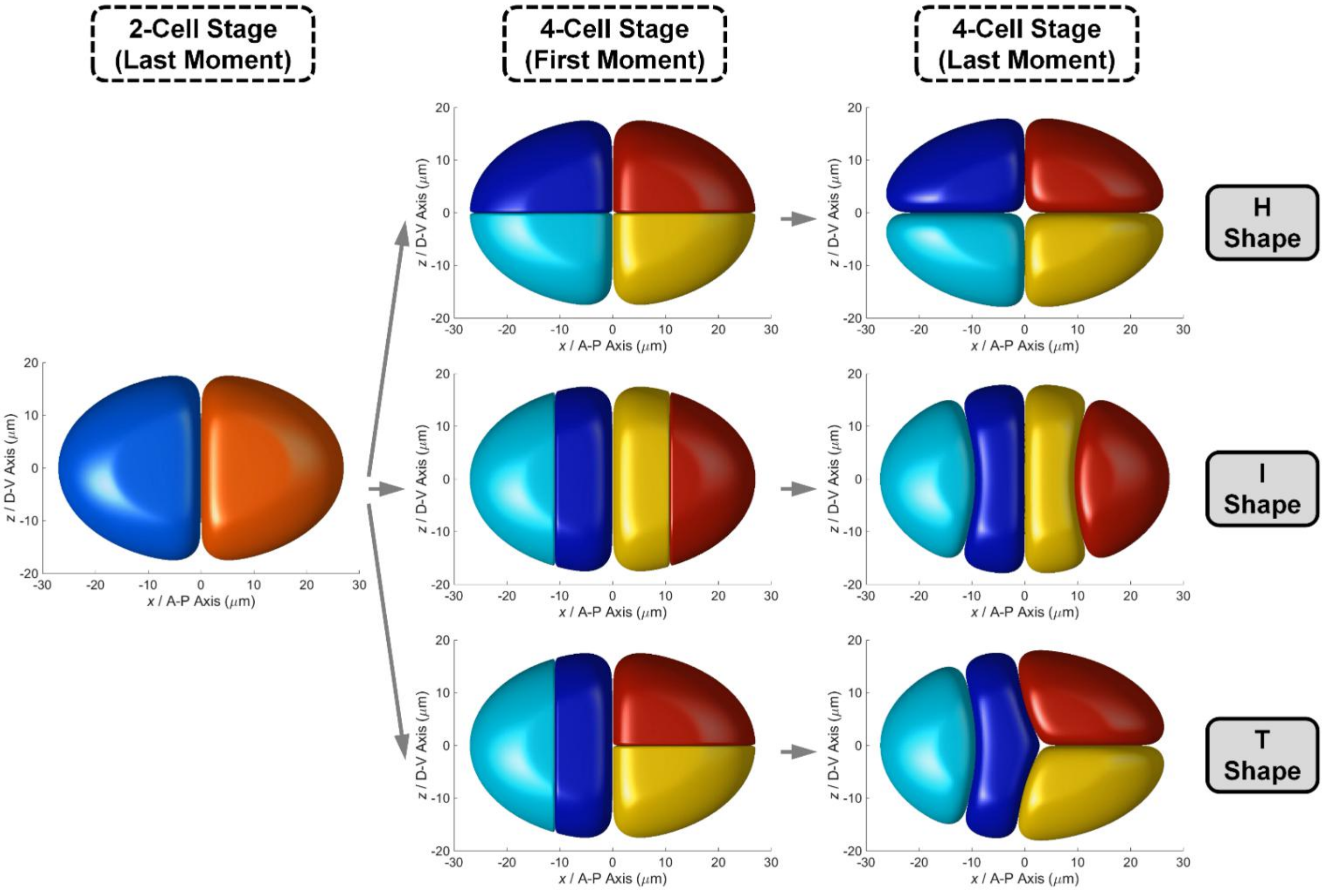

**Figure S3. Diversity of embryo morphology at 4-cell stage, generated by different cell division orientations.**

(A) "H" shape, with P0 dividing parallel to A-P axis, AB and P1 dividing synchronously and parallel to D-V axis.

(B) "I" shape, with P0 dividing parallel to A-P axis, AB and P1 dividing synchronously and parallel to A-P axis.

(C) "T" shape, with P0 dividing parallel to A-P axis, AB and P1 dividing synchronously but parallel to A-P and D-V axes respectively.

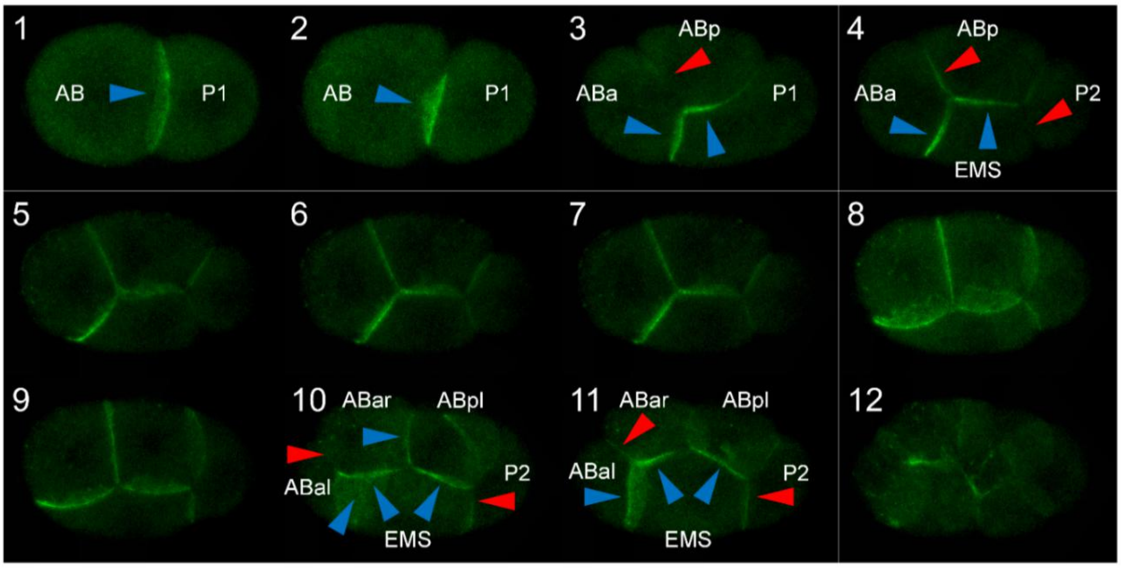


**Figure S4. Distribution of membrane-attached E-cadherin HMR-1 from 2- to 6-cell stages.** The interfaces with distinguishably high accumulation are indicated by blue arrows and the ones with relatively low accumulation are indicated by red arrows. The imaging time point of each figure (1 ~ 12) is denoted in its top left corner, respectively; figures 1 ~ 4, formation of 2- to 4-cell structures; figures 10 ~ 11, formation of 6-cell structure. Some interfaces near the focal planes are not identified or illustrated due to their blurry fluorescence, such as the contact interfaces of ABpl-ABpr, ABpl-P2 and ABpr-P2.

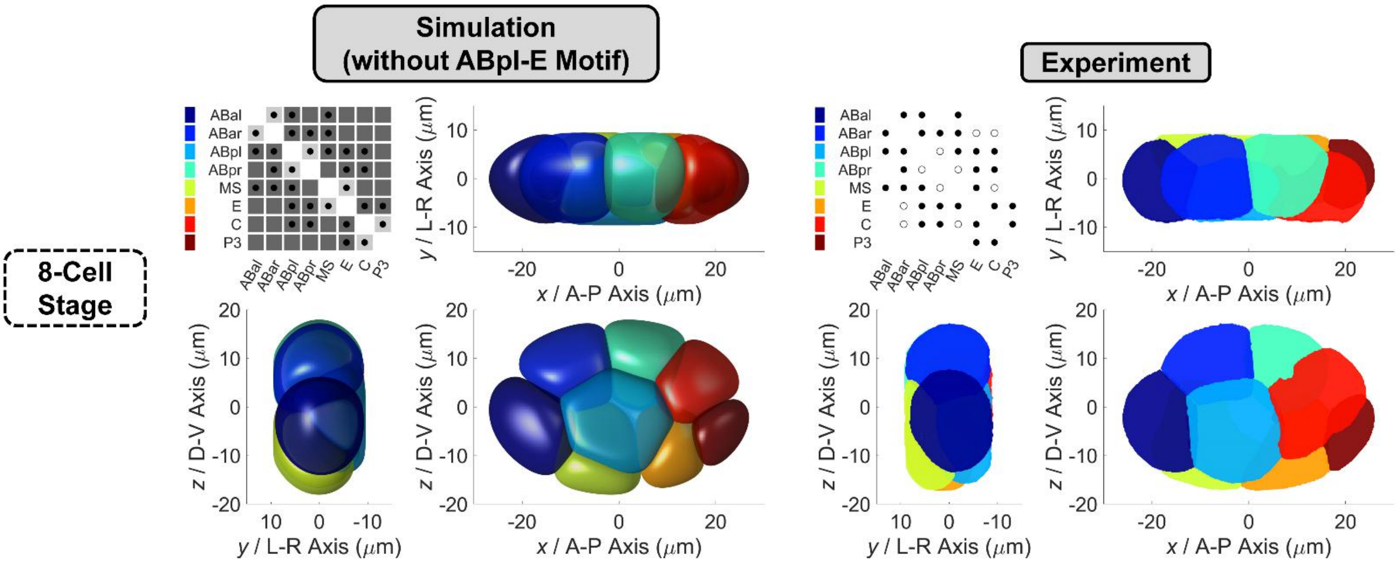


**Figure S5. Comparison of compressed embryo morphology between simulation and experiment at 8-cell stage, without attraction motif on ABpl-E contact, i.e., $\sigma_{\mathrm{ABpl\text{-}E}} = 0.9$.** Embryo morphology in simulation and experiment is respectively illustrated on the left and right in three orthogonal observation directions, while a cell-cell contact map is placed in the top left conners. About the map in simulation, dark gray and light gray shades denote strong attraction ($\sigma$ = 0.9) and weak attraction ($\sigma$' = 0.2) respectively, while black dots represent the contacted cell pairs. About the map in experiment, black dots represent the conserved contacted cell pairs which are reproducible in all the 4 embryo samples, while empty circles represent the unconserved contacted cell pairs which exist in at least one embryo sample, but not all of them (Cao et al., 2019). The relationship between cell identity and color is listed next to the contact map.

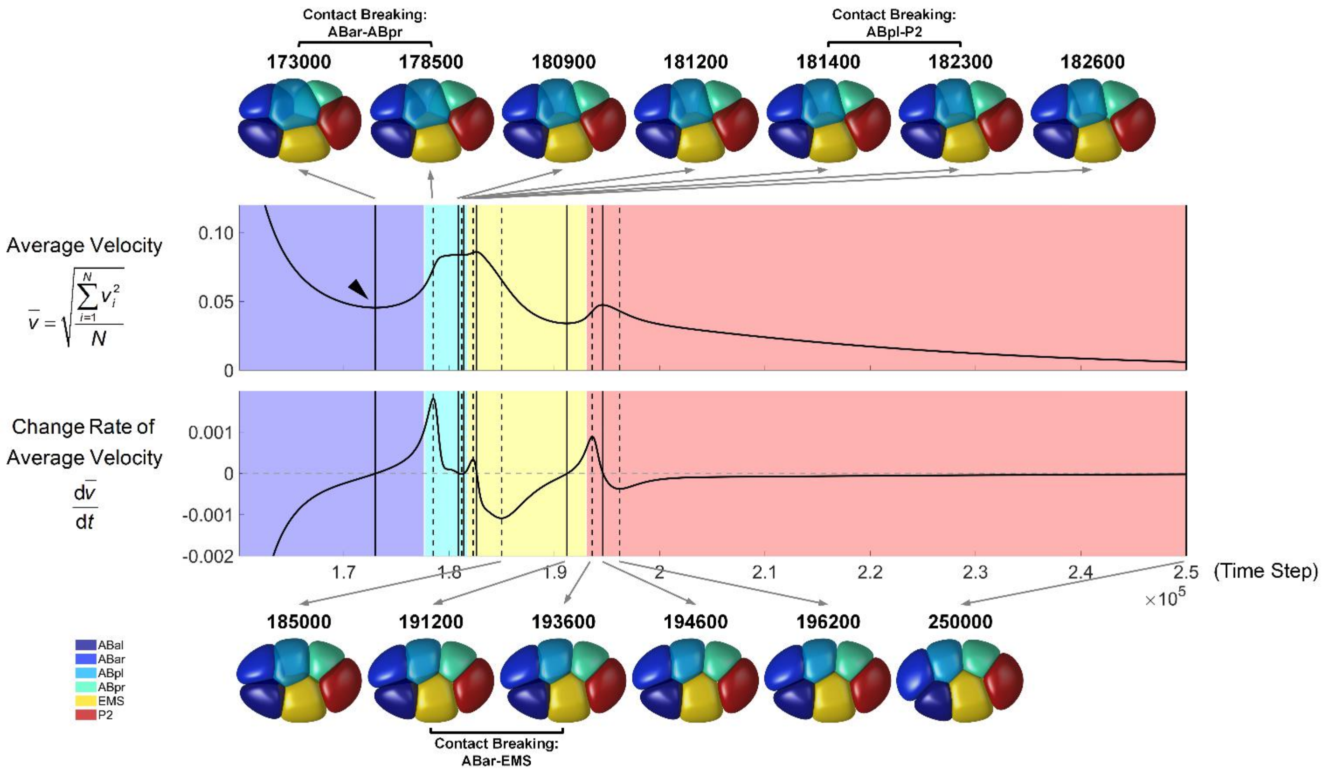


**Figure S6. Morphological evolution of a compressed embryo during 6-cell stage, with simulation time long enough for the whole system to reach mechanical equilibrium.** The curves of average velocity (upper) and its change rate (lower) are illustrated side by side. The solid and dashed vertical black lines denote the extreme points in the curves of average velocity and its change rate, respectively. The time point of the first quasi-steady state is indicated by a black triangle. The 3D structures at those time points in both curves are illustrated on top and bottom, pointed by gray arrows originating from their corresponding lines. The last structure in the bottom right is the system's terminal state approaching mechanical equilibrium. The change of cell-cell contact map is illustrated by different colors in the background, while the detail is written between two consecutive structures. The relationship between cell identity and color is listed in the bottom left corner.

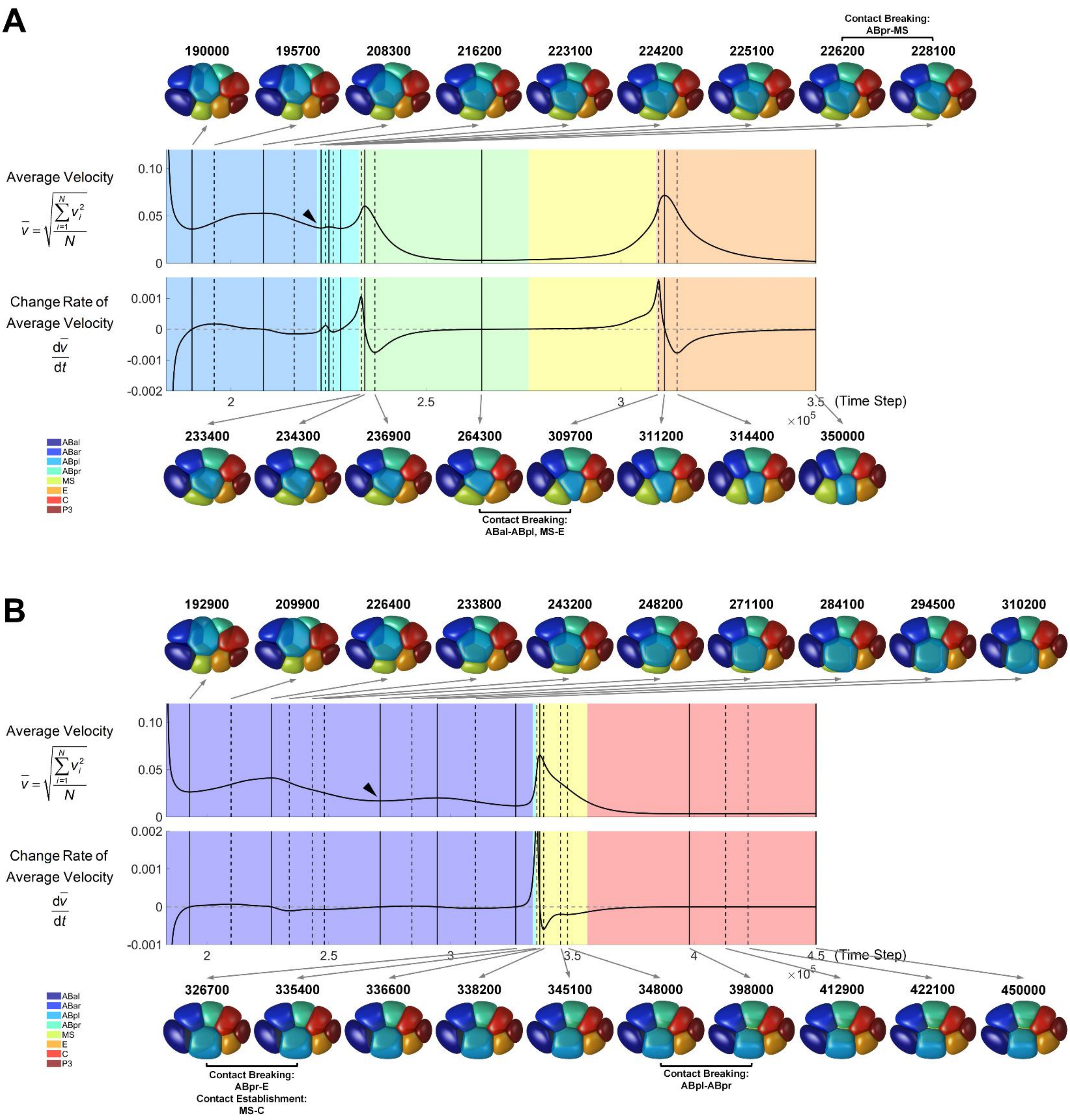


**Figure S7. Morphological evolution of a compressed embryo during 8-cell stage, with simulation time long enough for the whole system to reach mechanical equilibrium.**

(A) The upper panel, without attraction motif on ABpl-E contact, i.e., $\sigma_{\text{ABpl-E}} = 0.9$.

(B) The lower panel, with attraction motif on ABpl-E contact, i.e., $\sigma'_{\text{ABpl-E}} = 0.2$.

For each panel, the curves of average velocity (upper) and its change rate (lower) are illustrated side by side. The solid and dashed vertical black lines denote the extreme points in the curves of average velocity and its change rate, respectively. The time point of the second quasi-steady state is indicated by a black triangle. The 3D structures at those time points in both curves are illustrated on top and bottom, pointed by gray arrows originating from their corresponding lines. The last structure in the bottom right is the system's terminal state approaching mechanical equilibrium. The change of cell-cell contact map is illustrated by different colors in the background, while the detail is written between two consecutive structures. The relationship between cell identity and color is listed in the bottom left corner.

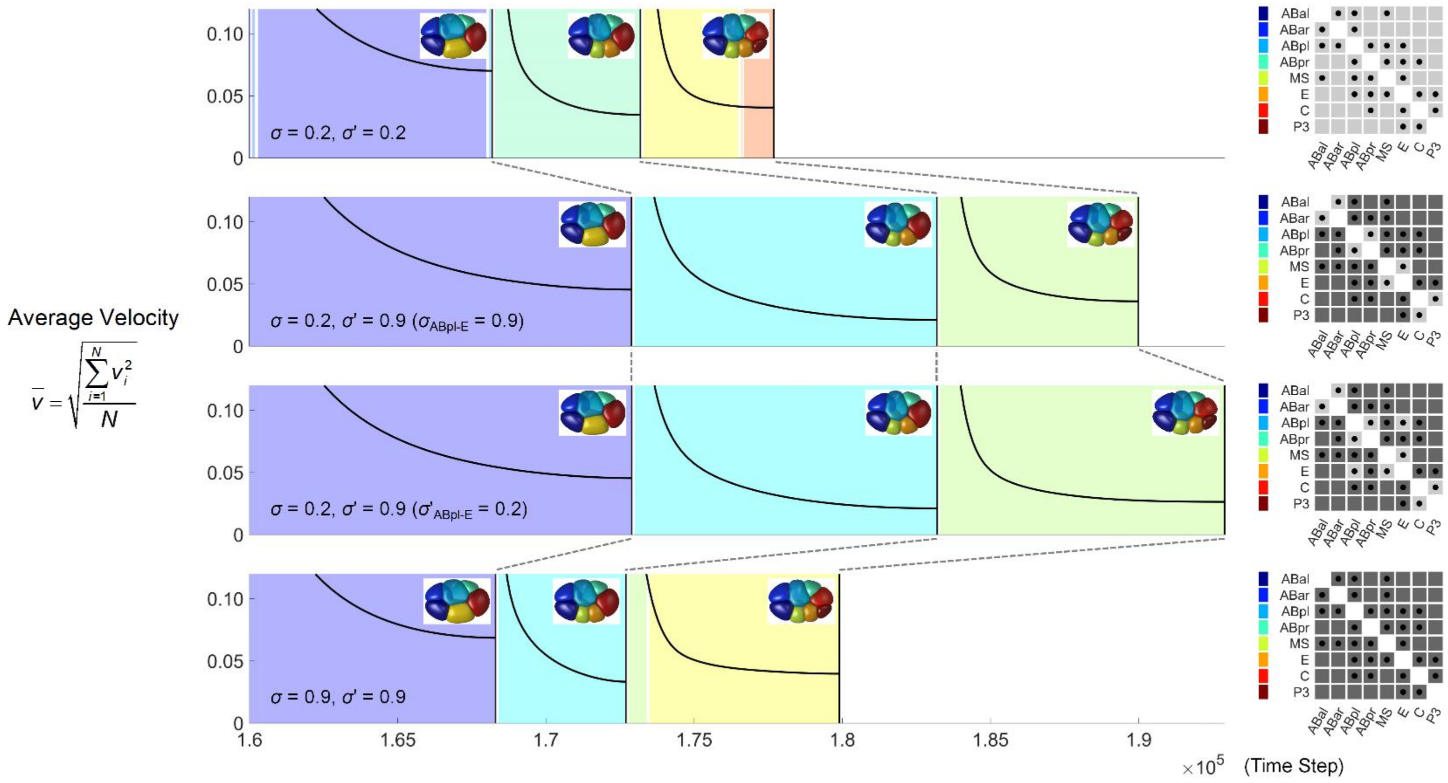


**Figure S8. Morphological evolution of a compressed embryo from 4- to 8-cell stages, with cell division timing controlled by quasi-steady state.** The 4 simulations are performed using globally weak attraction ($\sigma$ = 0.2, $\sigma$' = 0.2; 1st row), asymmetric attraction without ABpl-E motif ($\sigma$ = 0.9, $\sigma$' = 0.2, $\sigma_{\text{ABpl-E}}$ = 0.9; 2nd row), asymmetric attraction with ABpl-E motif ($\sigma$ = 0.9, $\sigma$' = 0.2, $\sigma'_{\text{ABpl-E}}$ = 0.2; 3rd row) and globally strong attraction ($\sigma$ = 0.9, $\sigma$' = 0.9; 4th row). The curves of average velocity are illustrated side by side. The 3D structures at the first quasi-steady state of 6-, 7- and 8-cell stages are illustrated near their corresponding time points. The change of cell-cell contact map is illustrated by different colors in the background. The cell-cell attraction matrix and contact map at 8-cell stage is illustrated on right; dark gray and light gray shades denote strong attraction ($\sigma$ = 0.9) and weak attraction ($\sigma$' = 0.2) respectively, while black dots represent the contacted cell pairs. The relationship between cell identity and color is listed next to the contact map.

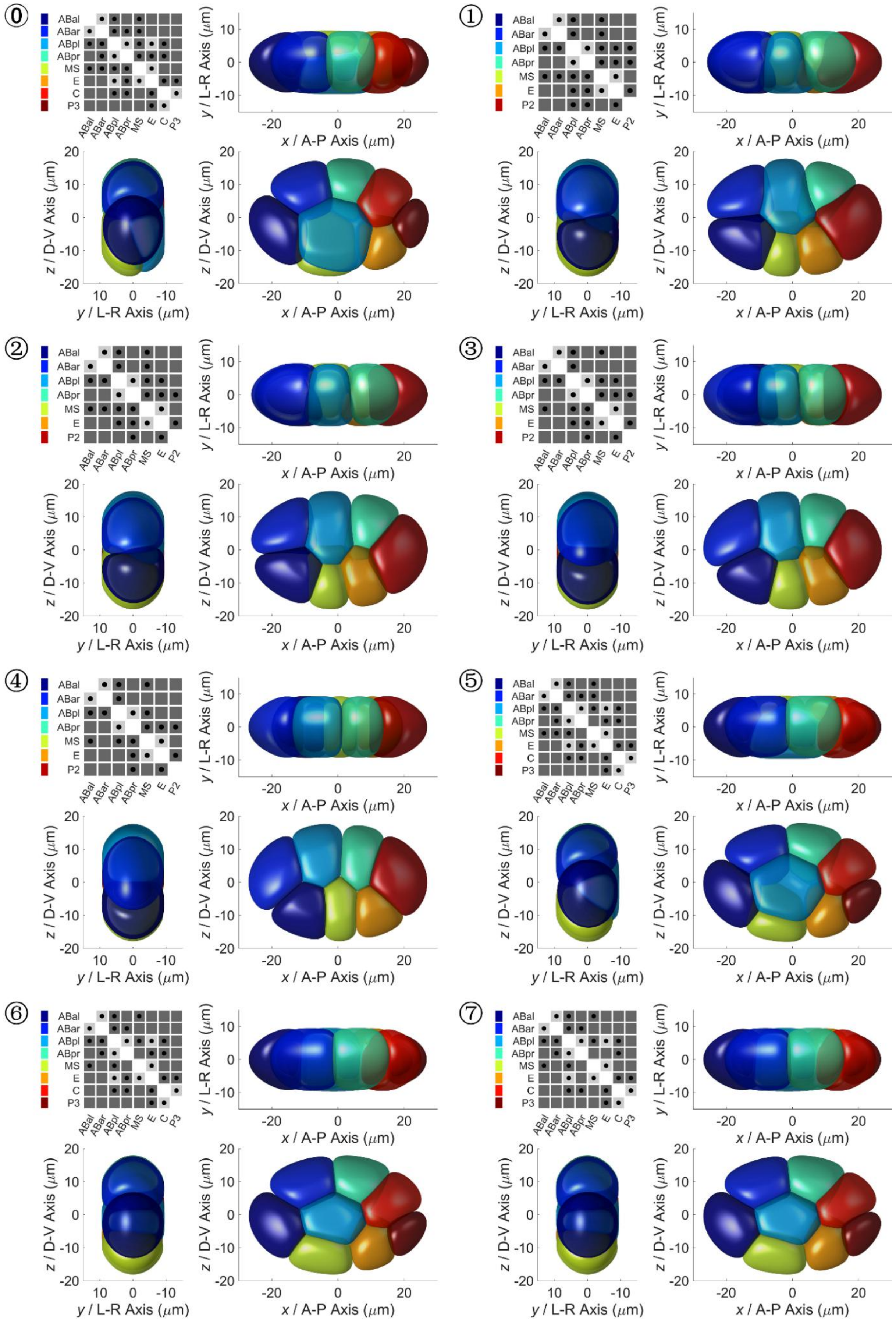


**Figure S9. Eight different developmental paths under perturbation on cell division timing.** The structures at quasi-steady state are labeled by ⓪, ①, ②, ③, ④, ⑤, ⑥, ⑦ from top left corner to bottom right corner, corresponding to Figure 4C. In each panel, embryo morphology in simulation is illustrated in three orthogonal observation directions, while a cell-cell contact map is placed in the top left conner. About the map in simulation, dark gray and light gray shades denote strong attraction ($\sigma$ = 0.9) and weak attraction ($\sigma$' = 0.2) respectively, while black dots represent the contacted cell pairs. The relationship between cell identity and color is listed in the bottom left corner.

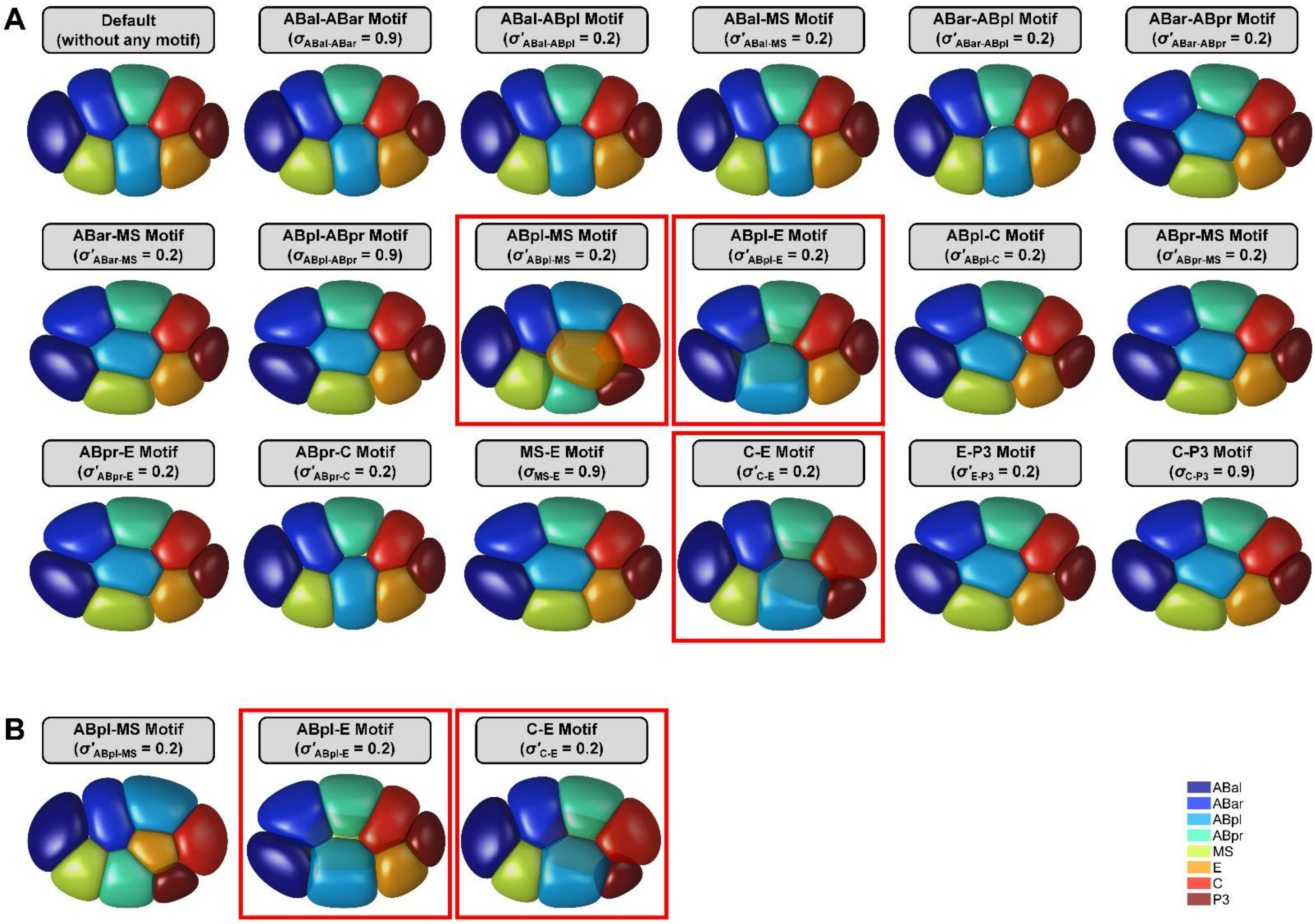


**Figure S10. Simulation for 8-cell stage with single motif added on cell-cell attraction matrix.**

(A) 3D structures at time point 350000. The ones with ABpl-MS, ABpl-E and C-E motifs are three-dimensional and highlighted with red rectangles.

(B) 3D structures at time point 450000. The ones with ABpl-E and C-E motifs are three-dimensional and highlighted with red rectangles.

The relationship between cell identity and color is listed in the bottom left corner.

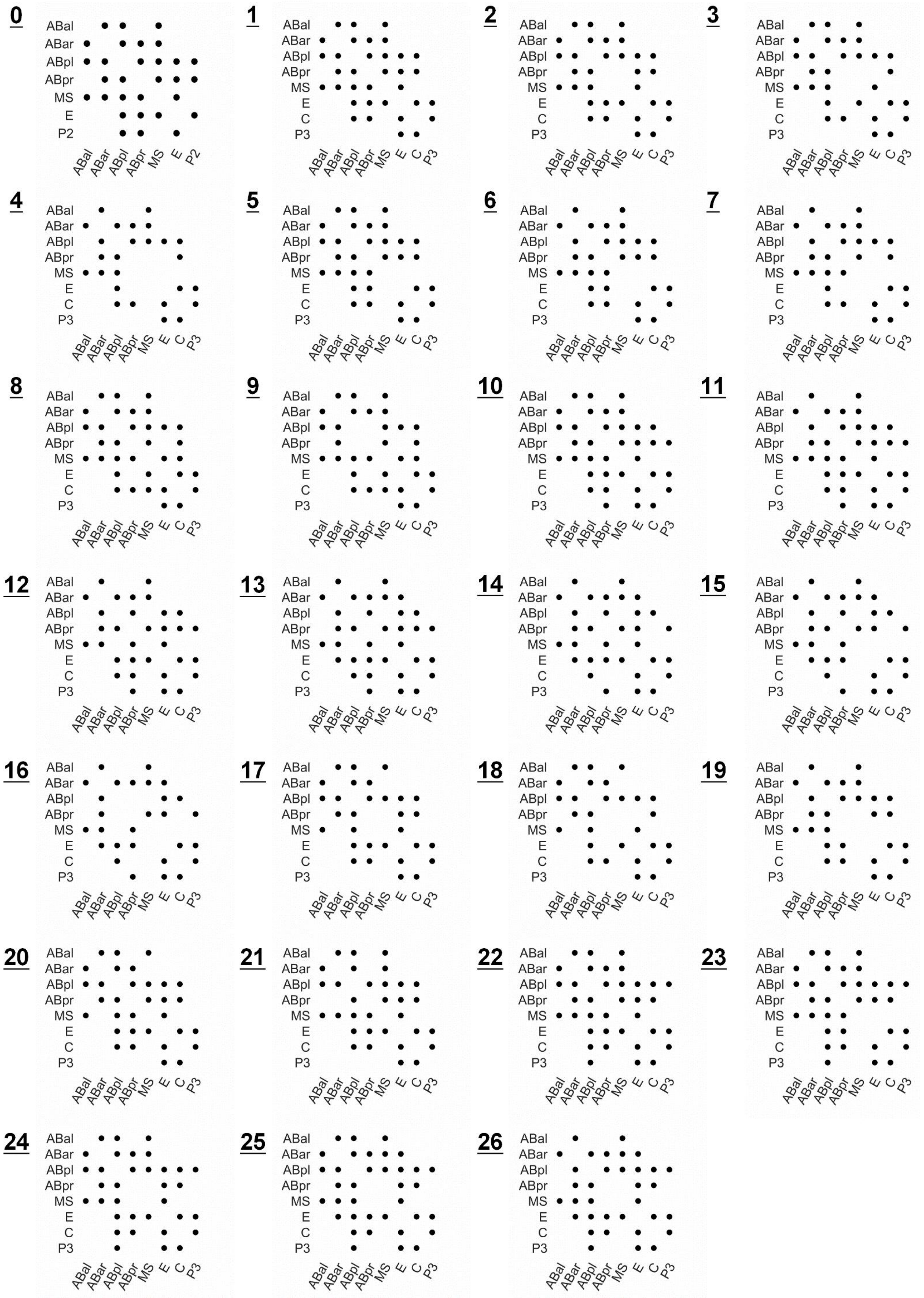

**Figure S11. Independent cell-cell contact map identified in the 17 simulations with single motif added on cell-cell attraction matrix.** The Topologies 0 ~ 26 corresponding to Figure 6A are listed from top left corner to bottom right corner. For each map, black dots represent the contacted cell pairs.

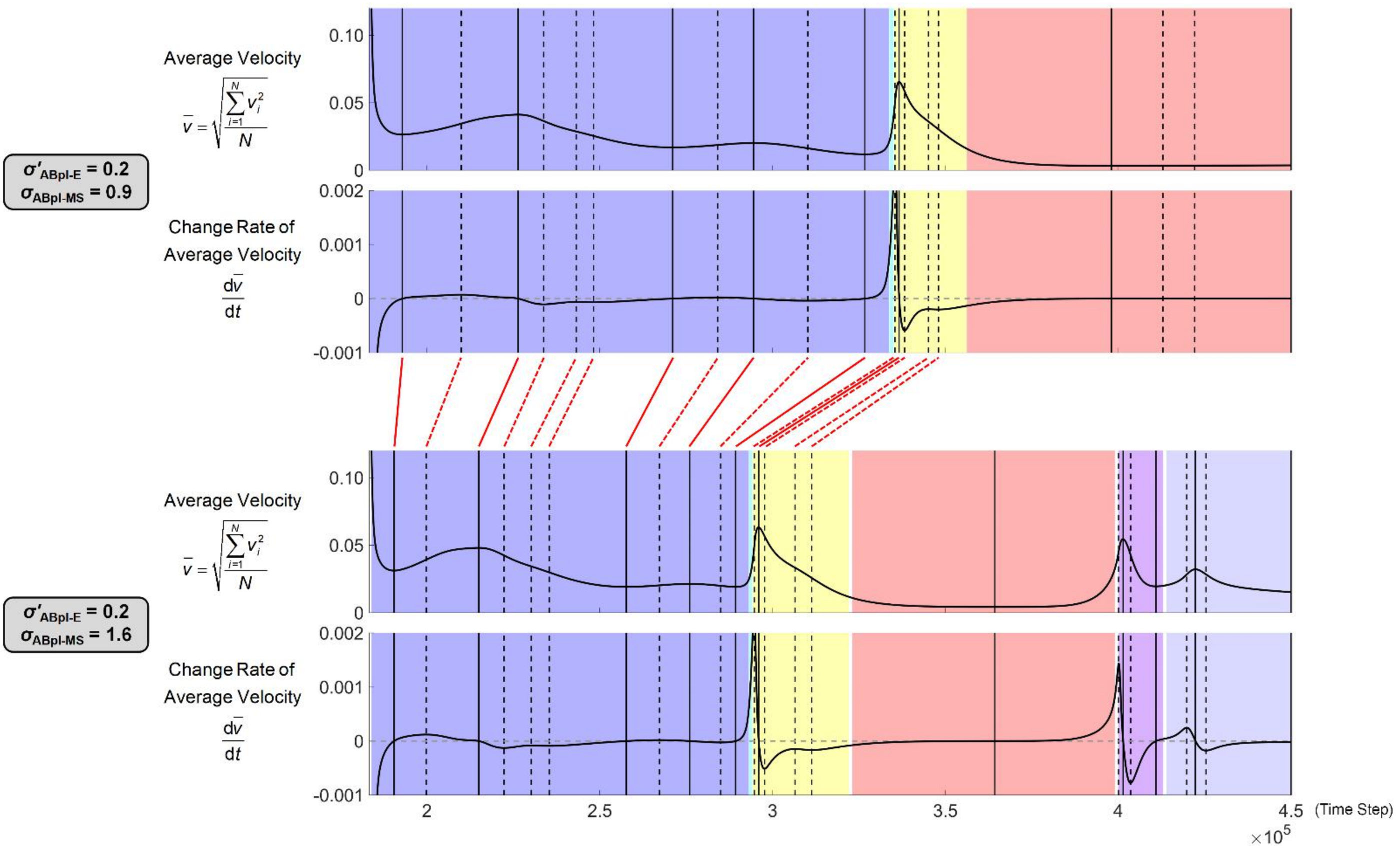


**Figure S12. Curves of average velocity and its change rate in embryos with $\sigma'_{ABpl\text{-}E}$ = 0.2, $\sigma_{ABpl\text{-}MS}$ = 0.9 (upper) and $\sigma'_{ABpl\text{-}E}$ = 0.2, $\sigma_{ABpl\text{-}MS}$ = 1.6 (lower) at 8-cell stage.** A total of 16 pairs of critical time points before mechanical equilibrium, which are defined by extreme points of the curves of average velocity and its change rate, are connected using solid and dashed red lines across the two panels (embryos).

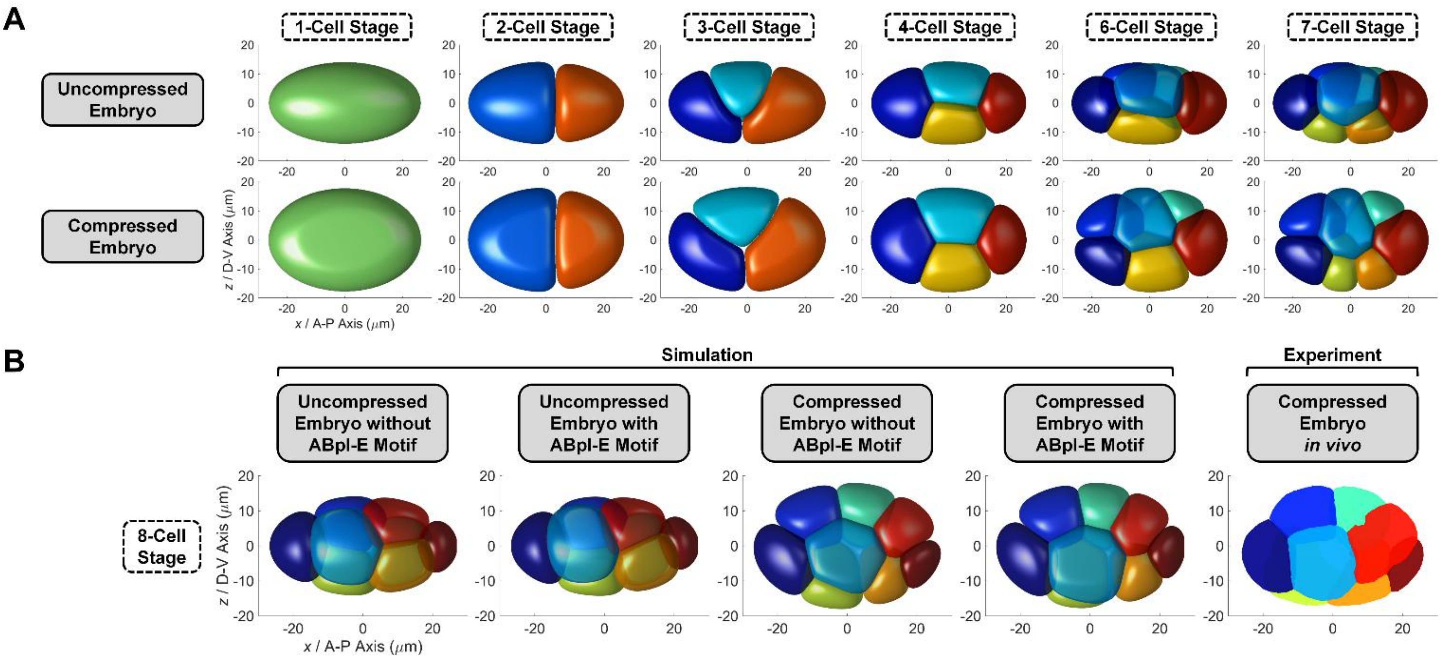


**Figure S13. Comparison between simulations on uncompressed and compressed embryos.**

(A) Embryo morphology of uncompressed (1st row) and compressed (2nd row) embryos from 1- to 7-cell stages, without any attraction motif added.

(B) Embryo morphology in uncompressed (1st and 2nd columns) and compressed (3rd and 4th columns) embryos at 8-cell stage, with or without ABpl-E motif, revealing that ABpl-E motif is essential for the compressed embryo but not for the uncompressed one.

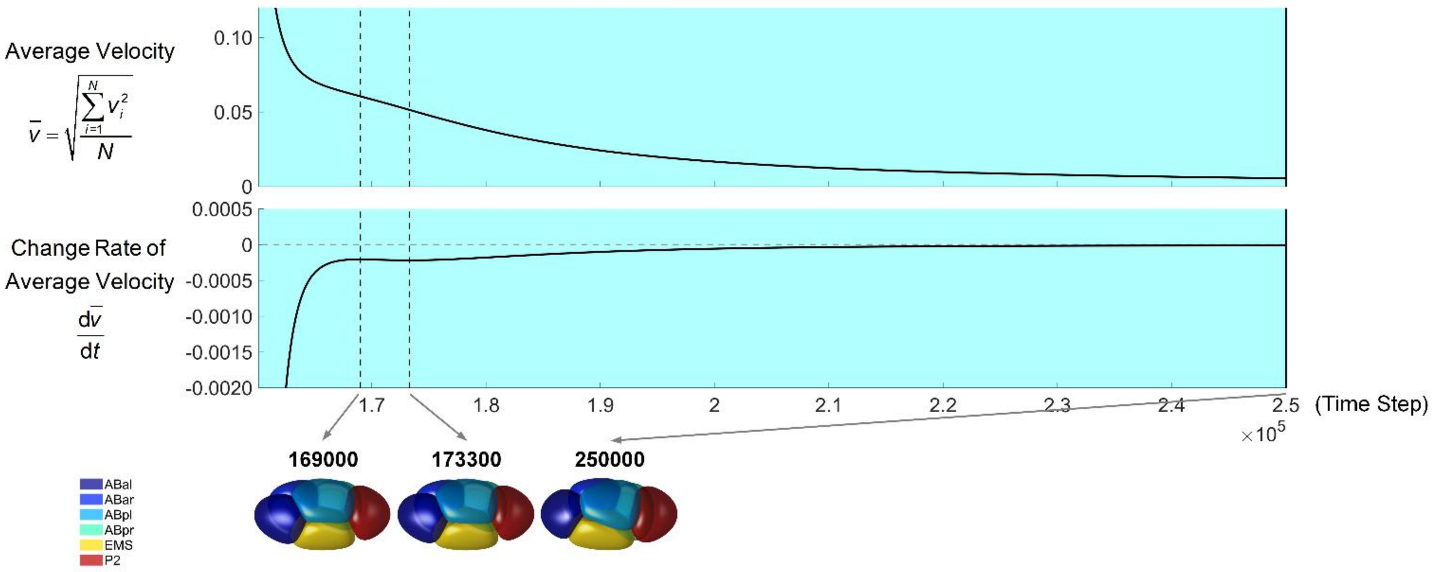


**Figure S14. Morphological evolution of an uncompressed embryo during 6-cell stage, with simulation time long enough for the whole system to reach mechanical equilibrium.** The curves of average velocity (upper) and its change rate (lower) are illustrated side by side. The solid and dashed vertical black lines denote the extreme points in the curves of average velocity and its change rate, respectively. The 3D structures at those time points in both curves are illustrated on bottom, pointed by gray arrows originating from their corresponding lines. The last structure in the bottom right is the system's terminal state approaching mechanical equilibrium. The relationship between cell identity and color is listed in the bottom left corner.

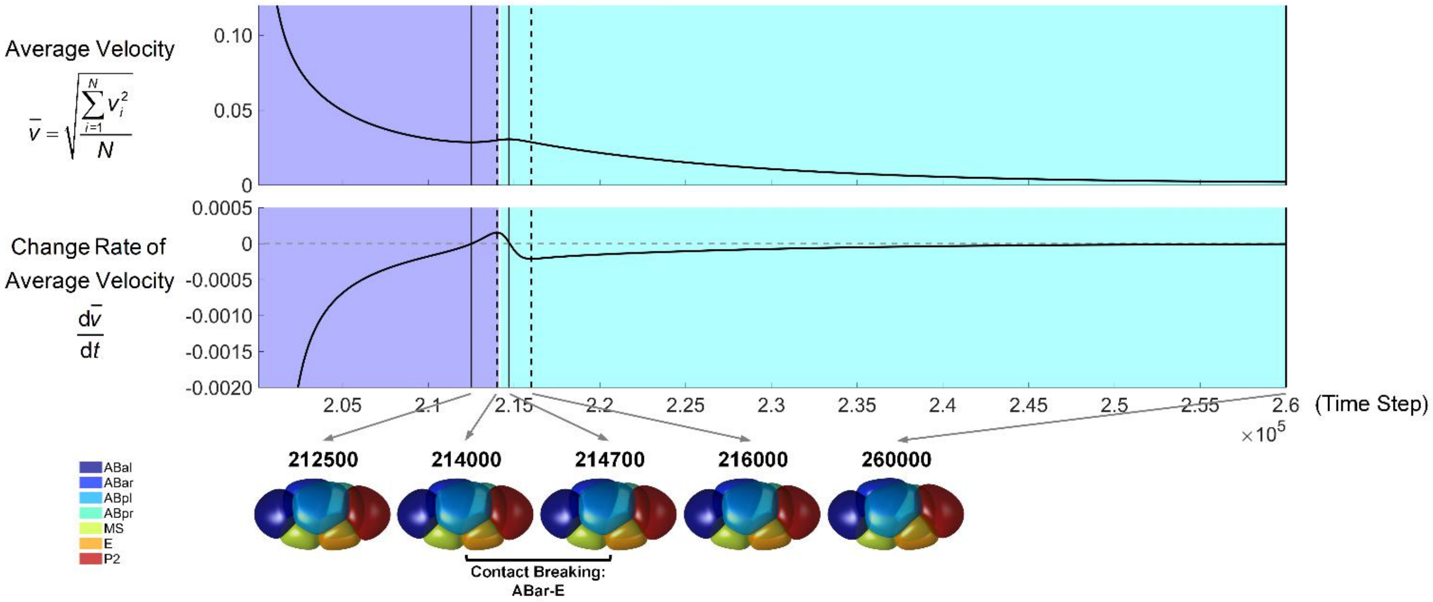


**Figure S15. Morphological evolution of an uncompressed embryo during 7-cell stage, with simulation time long enough for the whole system to reach mechanical equilibrium.** The curves of average velocity (upper) and its change rate (lower) are illustrated side by side. The solid and dashed vertical black lines denote the extreme points in the curves of average velocity and its change rate, respectively. The 3D structures at those time points in both curves are illustrated on bottom, pointed by gray arrows originating from their corresponding lines. The last structure in the bottom right is the system's terminal state approaching mechanical equilibrium. The change of cell-cell contact map is illustrated by different colors in the background, while the detail is written between two consecutive structures. The relationship between cell identity and color is listed in the bottom left corner.

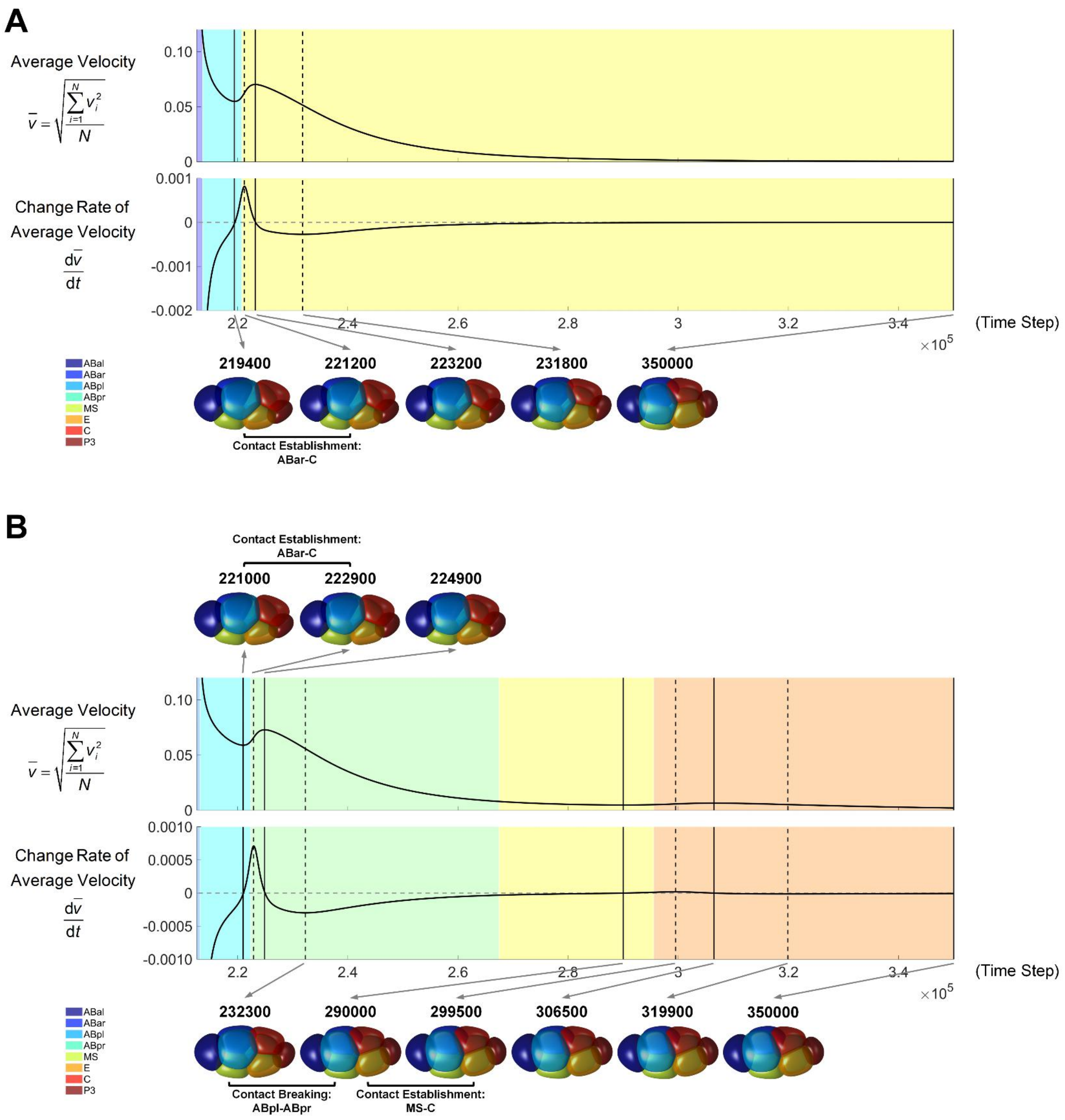


**Figure S16. Morphological evolution of an uncompressed embryo during 8-cell stage, with simulation time long enough for the whole system to reach mechanical equilibrium.**

(A) The upper panel, without attraction motif on ABpl-E contact, i.e., $\sigma_{\text{ABpl-E}} = 0.9$.

(B) The lower panel, with attraction motif on ABpl-E contact, i.e., $\sigma'_{\text{ABpl-E}} = 0.2$.

For each panel, the curves of average velocity (upper) and its change rate (lower) are illustrated side by side. The solid and dashed vertical black lines denote the extreme points in the curves of average velocity and its change rate, respectively. The 3D structures at those time points in both curves are illustrated on top and bottom, pointed by gray arrows originating from their corresponding lines. The last structure in the bottom right is the system's terminal state approaching mechanical equilibrium. The change of cell-cell contact map is illustrated by different colors in the background, while the detail is written between two consecutive structures. The relationship between cell identity and color is listed in the bottom left corner.

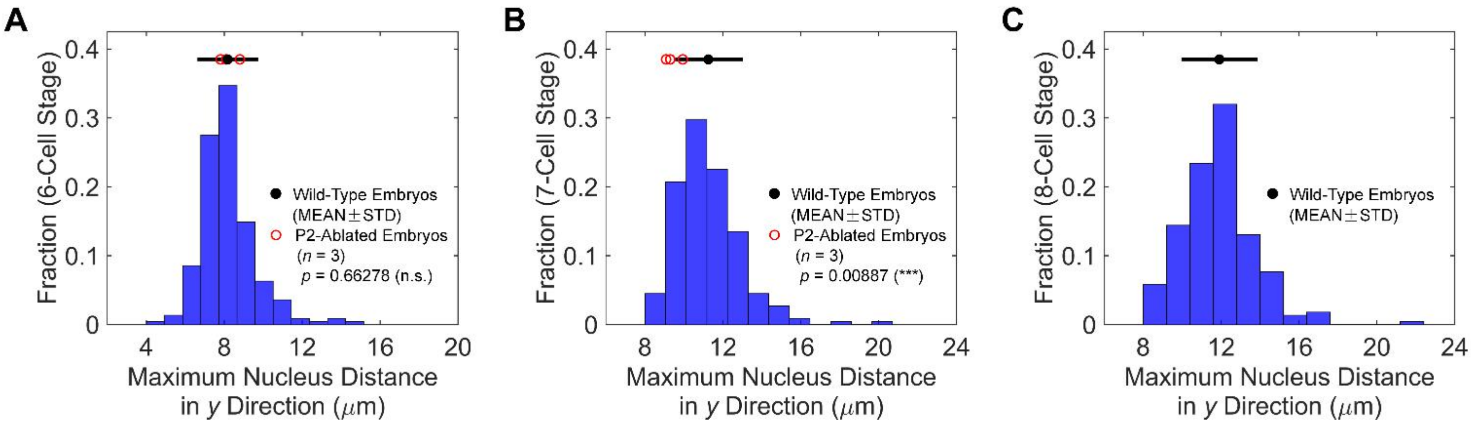


**Figure S17. Embryo width (maximum nucleus distance in *y* direction) of wild-type and P2-ablated embryos.**

(A) Distribution at the last time point of 6-cell stage.

(B) Distribution at the last time point of 7-cell stage.

(C) Distribution at the last time point of 8-cell stage.

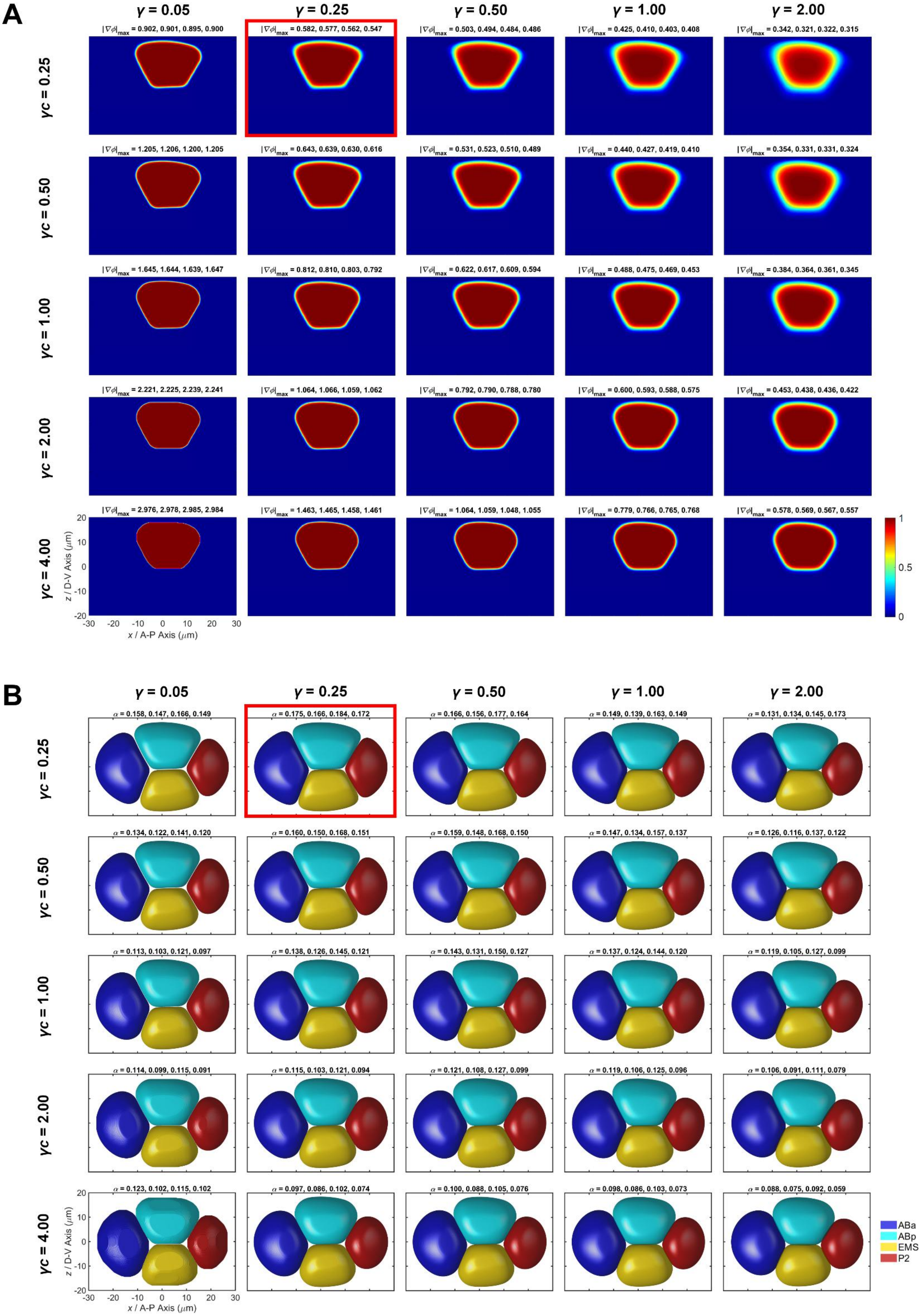
A
γ = 0.05
γ = 0.25
γ = 0.50
γ = 1.00
γ = 2.00
γc = 0.25
|∇φ|max = 0.902, 0.901, 0.895, 0.900
|∇φ|max = 0.582, 0.577, 0.562, 0.547
|∇φ|max = 0.503, 0.494, 0.484, 0.486
|∇φ|max = 0.425, 0.410, 0.403, 0.408
|∇φ|max = 0.342, 0.321, 0.322, 0.315
γc = 0.50
|∇φ|max = 1.205, 1.206, 1.200, 1.205
|∇φ|max = 0.643, 0.639, 0.630, 0.616
|∇φ|max = 0.531, 0.523, 0.510, 0.489
|∇φ|max = 0.440, 0.427, 0.419, 0.410
|∇φ|max = 0.354, 0.331, 0.331, 0.324
γc = 1.00
|∇φ|max = 1.645, 1.644, 1.639, 1.647
|∇φ|max = 0.812, 0.810, 0.803, 0.792
|∇φ|max = 0.622, 0.617, 0.609, 0.594
|∇φ|max = 0.488, 0.475, 0.469, 0.453
|∇φ|max = 0.384, 0.364, 0.361, 0.345
γc = 2.00
|∇φ|max = 2.221, 2.225, 2.239, 2.241
|∇φ|max = 1.064, 1.066, 1.059, 1.062
|∇φ|max = 0.792, 0.790, 0.788, 0.780
|∇φ|max = 0.600, 0.593, 0.588, 0.575
|∇φ|max = 0.453, 0.438, 0.436, 0.422
γc = 4.00
|∇φ|max = 2.976, 2.978, 2.985, 2.984
|∇φ|max = 1.463, 1.465, 1.458, 1.461
|∇φ|max = 1.064, 1.059, 1.048, 1.055
|∇φ|max = 0.779, 0.766, 0.765, 0.768
|∇φ|max = 0.578, 0.569, 0.567, 0.557
z / D-V Axis (μm)
x / A-P Axis (μm)
20 10 0 -10 -20
-30 -20 -10 0 10 20 30
1 0.5 0
B
γ = 0.05
γ = 0.25
γ = 0.50
γ = 1.00
γ = 2.00
γc = 0.25
α = 0.158, 0.147, 0.166, 0.149
α = 0.175, 0.166, 0.184, 0.172
α = 0.166, 0.156, 0.177, 0.164
α = 0.149, 0.139, 0.163, 0.149
α = 0.131, 0.134, 0.145, 0.173
γc = 0.50
α = 0.134, 0.122, 0.141, 0.120
α = 0.160, 0.150, 0.168, 0.151
α = 0.159, 0.148, 0.168, 0.150
α = 0.147, 0.134, 0.157, 0.137
α = 0.126, 0.116, 0.137, 0.122
γc = 1.00
α = 0.113, 0.103, 0.121, 0.097
α = 0.138, 0.126, 0.145, 0.121
α = 0.143, 0.131, 0.150, 0.127
α = 0.137, 0.124, 0.144, 0.120
α = 0.119, 0.105, 0.127, 0.099
γc = 2.00
α = 0.114, 0.099, 0.115, 0.091
α = 0.115, 0.103, 0.121, 0.094
α = 0.121, 0.108, 0.127, 0.099
α = 0.119, 0.106, 0.125, 0.096
α = 0.106, 0.091, 0.111, 0.079
γc = 4.00
α = 0.123, 0.102, 0.115, 0.102
α = 0.097, 0.086, 0.102, 0.074
α = 0.100, 0.088, 0.105, 0.076
α = 0.098, 0.086, 0.103, 0.073
α = 0.088, 0.075, 0.092, 0.059
z / D-V Axis (μm)
x / A-P Axis (μm)
20 10 0 -10 -20
-30 -20 -10 0 10 20 30
ABa
ABp
EMS
P2

**Figure S18. Sensitivity and comparison analysis on composite parameters $\gamma$ (0.05, 0.25, 0.50, 1.00, 2.00) and $\gamma c$ (0.25, 0.50, 1.00, 2.00, 4.00) with $g_e$ = 16.0.** The optimum fitting result ($\gamma = 0.25$, $c = 1.0$) is highlighted by red rectangles.

(A) Phase-field distribution at 4-cell stage, with interface transition quantified by maximum gradient $|\nabla \Phi|_{max}$.

(B) Embryo morphology pattern at 4-cell stage, with cell deformation quantified by coefficient $\alpha$ (Equations 13 and 14).

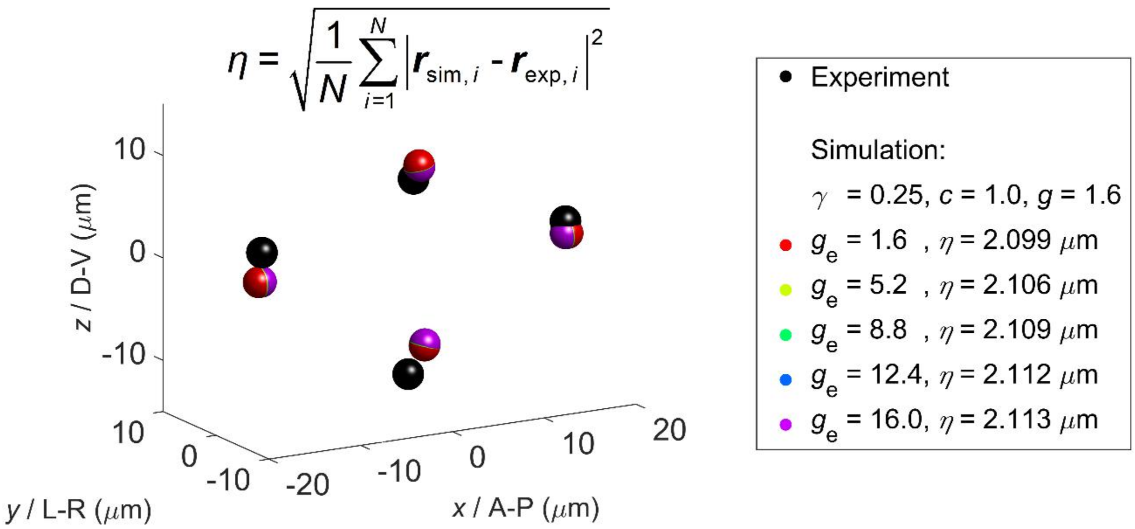


**Figure S19. Sensitivity and comparison analysis on parameter $g_e$ (1.6, 5.2, 8.8, 12.4, 16.0) with $\gamma$ = 0.25 and $c$ = 1.0.** The average positional variation $\eta$ is used to evaluate the deviation between simulation and experiment structures (Equation 15).

**Table S1. Information of the embryos collected from datasets produced previously.**

| Group | Embryo Number | Original Reference of Strain | Original Reference of Dataset | Nucleus Marker | Membrane Marker | Usage |
|---|---|---|---|---|---|---|
| 1 | 1 | Murray et al., 2012 (RW10425) | Guan et al., 2019 | √ | × | 1. Division Orientation of the Cells before 4-Cell Stage (i.e., P0, AB, P1) |
| 2 | 13 | Murray et al., 2012 (RW10112) | Cao et al., 2019 | √ | √ | 1. Division Orientation of the Cells since 4-Cell Stage (i.e., ABa, ABp, EMS, P2) |
| 3 | 4 | Cao et al., 2019 (ZZY0655) | Cao et al., 2019 | √ | √ | 1. Division Orientation of the Cells since 4-Cell Stage (i.e., ABa, ABp, EMS, P2) 2. Cell Morphology of the Cells since 4-Cell Stage (i.e., ABa, ABp, EMS, P2, ABal, ABar, ABpl, ABpr, MS, E, C, P3) |

**Table S2. Volume and division orientation of the cells up to 8-cell stage.**

| Cell Identity | Cell Volume (μm$^3$) | Cell Division Orientation (Normalized, [*x*, *y*, *z*]) |
|---|---|---|
| P0 | 20107.335 | '-0.992,-0.123,0.010' |
| AB | 11938.845 | '0.767,0.078,0.637' |
| P1 | 8168.490 | '0.954,0.040,0.297' |
| ABa | 6020.879 | '0.313,0.684,0.659' |
| ABp | 5917.966 | '0.376,0.913,0.160' |
| EMS | 4690.054 | '0.978,-0.196,-0.073' |
| P2 | 3478.436 | '0.255,0.155,-0.954' |
| ABal | 2879.988 | '0.424,0.412,-0.807' |
| ABar | 3010.672 | '0.478,-0.839,0.260' |
| ABpl | 3240.075 | '0.689,0.354,-0.632' |
| ABpr | 2642.929 | '0.349,0.123,-0.929' |
| MS | 2389.470 | '0.920,0.062,0.388' |
| E | 2080.275 | '0.938,-0.131,0.320' |
| C | 2365.737 | '0.976,0.103,0.191' |
| P3 | 1116.712 | '-0.266,0.201,-0.943' |
| ABala | 1269.628 | '0.700,-0.625,0.347' |
| ABalp | 1518.182 | '0.771,-0.564,-0.294' |
| ABara | 1380.479 | '0.960,-0.119,0.255' |
| ABarp | 1616.351 | '0.965,-0.144,-0.221' |
| ABpla | 1859.459 | '0.995,0.050,-0.081' |
| ABplp | 1353.634 | '0.971,-0.115,0.211' |
| ABpra | 1389.208 | '0.939,-0.229,-0.257' |
| ABprp | 1230.518 | '0.974,-0.223,0.041' |

Note: For P0, AB and P1, cell volume is calculated with the ones of their progenies (i.e., ABa, ABp, EMS, P2). Cell division orientations of P0, AB and P1 are measured using 1 wild-type embryo imaged since 1-cell stage (Table S1) (Guan et al., 2019). For the others appearing since 4-cell stage, cell volumes are the averages obtained from 4 wild-type embryos with membrane marker, while cell division orientations are the averages obtained from all the 17 wild-type embryos with nucleus marker (Table S1) (Cao et al., 2019).

**Table S3. Cell surface area, cell-cell contact relationship and area at 4-cell stage (Figure 2B and 2C).**

| 'Cell Identity' | | 'ABa' | 'ABp' | 'EMS' | 'P2' |
|---|---|---|---|---|---|
| | 'Surface Area' | 30818 | 31141 | 31341 | 23667 |
| | | 32121 | 32105 | 32590 | 23450 |
| | | 29655 | 29935 | 29408 | 22699 |
| | | 29104 | 32103 | 28542 | 21670 |
| 'ABa' | 30818 | 0 | 6591 | 5718 | 0 |
| | 32121 | 0 | 6275 | 5075 | 0 |
| | 29655 | 0 | 5859 | 4215 | 0 |
| | 29104 | 0 | 5902 | 4692 | 0 |
| 'ABp' | 31141 | 6591 | 0 | 4472 | 5421 |
| | 32105 | 6275 | 0 | 5361 | 4957 |
| | 29935 | 5859 | 0 | 4876 | 4316 |
| | 32103 | 5902 | 0 | 4417 | 4523 |
| 'EMS' | 31341 | 5718 | 4472 | 0 | 2620 |
| | 32590 | 5075 | 5361 | 0 | 1845 |
| | 29408 | 4215 | 4876 | 0 | 2372 |
| | 28542 | 4692 | 4417 | 0 | 2385 |
| 'P2' | 23667 | 0 | 5421 | 2620 | 0 |
| | 23450 | 0 | 4957 | 1845 | 0 |
| | 22699 | 0 | 4316 | 2372 | 0 |
| | 21670 | 0 | 4523 | 2385 | 0 |

Note: Cell surface area is quantified by the total number of pixels surrounding a cell, while cell-cell contact area is quantified by the total number of pixels adjacent to both cells (sample size = 4; spatial resolution ≈ 0.225 μm / pixel in three orthogonal coordinates). "NaN" represents a cell which has its own nucleus but hasn't completely finished cytokinesis with its sister, which together share the same membrane boundary. "0" means that the two independent cells don't contact with each other at all. The cell-cell contacts with no "0" is reproducible in all the 4 segmented wild-type embryo samples (Table S1) (Cao et al., 2019).

**Table S4. Cell surface area, cell-cell contact relationship and area at 6-cell stage (Figure 3A).**

| 'Cell Identity' | | 'ABal' | 'ABar' | 'ABpl' | 'ABpr' | 'EMS' | 'P2' |
|---|---|---|---|---|---|---|---|
| | 'Surface Area' | 18213 | 21100 | 22378 | 17609 | 26983 | 23360 |
| | | NaN | NaN | NaN | NaN | 29589 | 21621 |
| | | 17076 | 20312 | 23168 | 17280 | 24715 | 21648 |
| | | 16279 | 18435 | 22993 | 17502 | 24186 | 19801 |
| 'ABal' | 18213 | 0 | 893 | 2162 | 0 | 3830 | 0 |
| | NaN | NaN | NaN | NaN | NaN | NaN | NaN |
| | 17076 | 0 | 1107 | 1667 | 0 | 2716 | 0 |
| | 16279 | 0 | 1553 | 986 | 0 | 2821 | 0 |
| 'ABar' | 21100 | 893 | 0 | 4502 | 2761 | 2238 | 0 |
| | NaN | NaN | NaN | NaN | NaN | NaN | NaN |
| | 20312 | 1107 | 0 | 4702 | 1711 | 1729 | 0 |
| | 18435 | 1553 | 0 | 4030 | 1188 | 1403 | 0 |
| 'ABpl' | 22378 | 2162 | 4502 | 0 | 590 | 2528 | 2182 |
| | NaN | NaN | NaN | NaN | NaN | NaN | NaN |
| | 23168 | 1667 | 4702 | 0 | 906 | 3292 | 1181 |
| | 22993 | 986 | 4030 | 0 | 2231 | 2584 | 1184 |
| 'ABpr' | 17609 | 0 | 2761 | 590 | 0 | 1826 | 3604 |
| | NaN | NaN | NaN | NaN | NaN | NaN | NaN |
| | 17280 | 0 | 1711 | 906 | 0 | 1527 | 3122 |
| | 17502 | 0 | 1188 | 2231 | 0 | 1420 | 2639 |
| 'EMS' | 26983 | 3830 | 2238 | 2528 | 1826 | 0 | 3240 |
| | 29589 | NaN | NaN | NaN | NaN | 0 | 2236 |
| | 24715 | 2716 | 1729 | 3292 | 1527 | 0 | 3066 |
| | 24186 | 2821 | 1403 | 2584 | 1420 | 0 | 2934 |

| | | | | | | | |
|---|---|---|---|---|---|---|---|
| 'P2' | 23360 | 0 | 0 | 2182 | 3604 | 3240 | 0 |
| | 21621 | NaN | NaN | NaN | NaN | 2236 | 0 |
| | 21648 | 0 | 0 | 1181 | 3122 | 3066 | 0 |
| | 19801 | 0 | 0 | 1184 | 2639 | 2934 | 0 |

Note: Cell surface area is quantified by the total number of pixels surrounding a cell, while cell-cell contact area is quantified by the total number of pixels adjacent to both cells (sample size = 4; spatial resolution ≈ 0.225 μm / pixel in three orthogonal coordinates). "NaN" represents a cell which has its own nucleus but hasn't completely finished cytokinesis with its sister, which together share the same membrane boundary. "0" means that the two independent cells don't contact with each other at all. The cell-cell contacts with no "0" is reproducible in all the 4 segmented wild-type embryo samples (Table S1) (Cao et al., 2019).

### Table S5. Cell surface area, cell-cell contact relationship and area at 7-cell stage (Figure 3B).

| 'Cell Identity' | | 'ABal' | 'ABar' | 'ABpl' | 'ABpr' | 'MS' | 'E' | 'P2' |
|---|---|---|---|---|---|---|---|---|
| | 'Surface Area' | 18081 | 21185 | 23654 | 18836 | 18411 | 16192 | 19575 |
| | | 18542 | 21088 | 25031 | 18631 | NaN | NaN | 19947 |
| | | 17037 | 20914 | 24066 | 18077 | 16810 | 13463 | 20158 |
| | | 16475 | 18717 | 23418 | 18987 | NaN | NaN | 18167 |
| 'ABal' | 18081 | 0 | 1397 | 2972 | 0 | 3328 | 0 | 0 |
| | 18542 | 0 | 1917 | 2210 | 0 | NaN | NaN | 0 |
| | 17037 | 0 | 1704 | 2110 | 0 | 2873 | 0 | 0 |
| | 16475 | 0 | 1356 | 1513 | 0 | NaN | NaN | 0 |
| 'ABar' | 21185 | 1397 | 0 | 3770 | 3438 | 1988 | 0 | 0 |
| | 21088 | 1917 | 0 | 3937 | 2694 | NaN | NaN | 0 |
| | 20914 | 1704 | 0 | 4953 | 2360 | 1384 | 0 | 0 |
| | 18717 | 1356 | 0 | 4280 | 1975 | NaN | NaN | 0 |
| 'ABpl' | 23654 | 2972 | 3770 | 0 | 960 | 1662 | 2316 | 1847 |
| | 25031 | 2210 | 3937 | 0 | 1614 | NaN | NaN | 1628 |
| | 24066 | 2110 | 4953 | 0 | 1622 | 1892 | 906 | 1454 |
| | 23418 | 1513 | 4280 | 0 | 2628 | NaN | NaN | 1476 |
| 'ABpr' | 18836 | 0 | 3438 | 960 | 0 | 1540 | 1976 | 2599 |
| | 18631 | 0 | 2694 | 1614 | 0 | NaN | NaN | 3014 |
| | 18077 | 0 | 2360 | 1622 | 0 | 1153 | 1135 | 3374 |
| | 18987 | 0 | 1975 | 2628 | 0 | NaN | NaN | 2608 |
| 'MS' | 18411 | 3328 | 1988 | 1662 | 1540 | 0 | 455 | 0 |
| | NaN | NaN | NaN | NaN | NaN | NaN | NaN | NaN |
| | 16810 | 2873 | 1384 | 1892 | 1153 | 0 | 797 | 0 |
| | NaN | NaN | NaN | NaN | NaN | NaN | NaN | NaN |
| 'E' | 16192 | 0 | 0 | 2316 | 1976 | 455 | 0 | 2846 |
| | NaN | NaN | NaN | NaN | NaN | NaN | NaN | NaN |
| | 13463 | 0 | 0 | 906 | 1135 | 797 | 0 | 2422 |
| | NaN | NaN | NaN | NaN | NaN | NaN | NaN | NaN |
| 'P2' | 19575 | 0 | 0 | 1847 | 2599 | 0 | 2846 | 0 |
| | 19947 | 0 | 0 | 1628 | 3014 | NaN | NaN | 0 |
| | 20158 | 0 | 0 | 1454 | 3374 | 0 | 2422 | 0 |
| | 18167 | 0 | 0 | 1476 | 2608 | NaN | NaN | 0 |

Note: Cell surface area is quantified by the total number of pixels surrounding a cell, while cell-cell contact area is quantified by the total number of pixels adjacent to both cells (sample size = 4; spatial resolution ≈ 0.225 μm / pixel in three orthogonal coordinates). "NaN" represents a cell which has its own nucleus but hasn't completely finished cytokinesis with its sister, which together share the same membrane boundary. "0" means that the two independent cells don't contact with each other at all. The cell-cell contacts with no "0" is reproducible in all the 4 segmented wild-type embryo samples (Table S1) (Cao et al., 2019).

**Table S6. Cell surface area, cell-cell contact relationship and area at 8-cell stage (Figure 3C).**

| 'Cell Identity' | | 'ABal' | 'ABar' | 'ABpl' | 'ABpr' | 'MS' | 'E' | 'C' | 'P3' |
|---|---|---|---|---|---|---|---|---|---|
| | 'Surface Area' | 18405 | 19431 | 19462 | 16561 | 20681 | 18184 | 19945 | 9994 |
| | | 18315 | 19500 | 21850 | 17480 | 23034 | 15782 | 19032 | 10357 |
| | | 17130 | 18899 | 19348 | 16548 | 21096 | 14890 | 19749 | 9441 |
| | | 15950 | 17385 | 19825 | 16355 | 17064 | 15405 | 17163 | 9163 |
| 'ABal' | 18405 | 0 | 1679 | 2795 | 0 | 2998 | 0 | 0 | 0 |
| | 18315 | 0 | 1766 | 2692 | 0 | 2524 | 0 | 0 | 0 |
| | 17130 | 0 | 1855 | 2642 | 0 | 1979 | 0 | 0 | 0 |
| | 15950 | 0 | 1315 | 2297 | 0 | 2557 | 0 | 0 | 0 |
| 'ABar' | 19431 | 1679 | 0 | 2170 | 2795 | 2367 | 14 | 760 | 0 |
| | 19500 | 1766 | 0 | 2610 | 3192 | 1558 | 0 | 3 | 0 |
| | 18899 | 1855 | 0 | 2448 | 3042 | 1340 | 0 | 202 | 0 |
| | 17385 | 1315 | 0 | 2603 | 2719 | 861 | 0 | 0 | 0 |
| 'ABpl' | 19462 | 2795 | 2170 | 0 | 0 | 2119 | 1770 | 2011 | 0 |
| | 21850 | 2692 | 2610 | 0 | 195 | 4173 | 1664 | 1094 | 0 |
| | 19348 | 2642 | 2448 | 0 | 0 | 3736 | 816 | 1323 | 0 |
| | 19825 | 2297 | 2603 | 0 | 612 | 2633 | 1361 | 1547 | 0 |
| 'ABpr' | 16561 | 0 | 2795 | 0 | 0 | 897 | 1646 | 2911 | 0 |
| | 17480 | 0 | 3192 | 195 | 0 | 1224 | 1153 | 3760 | 0 |
| | 16548 | 0 | 3042 | 0 | 0 | 1247 | 937 | 3439 | 0 |
| | 16355 | 0 | 2719 | 612 | 0 | 0 | 855 | 3162 | 0 |
| 'MS' | 20681 | 2998 | 2367 | 2119 | 897 | 0 | 2407 | 0 | 0 |
| | 23034 | 2524 | 1558 | 4173 | 1224 | 0 | 1891 | 11 | 0 |
| | 21096 | 1979 | 1340 | 3736 | 1247 | 0 | 2045 | 212 | 0 |
| | 17064 | 2557 | 861 | 2633 | 0 | 0 | 898 | 0 | 0 |
| 'E' | 18184 | 0 | 14 | 1770 | 1646 | 2407 | 0 | 2246 | 2408 |
| | 15782 | 0 | 0 | 1664 | 1153 | 1891 | 0 | 1038 | 2206 |
| | 14890 | 0 | 0 | 816 | 937 | 2045 | 0 | 2108 | 2037 |
| | 15405 | 0 | 0 | 1361 | 855 | 898 | 0 | 1677 | 2099 |
| 'C' | 19945 | 0 | 760 | 2011 | 2911 | 0 | 2246 | 0 | 1866 |
| | 19032 | 0 | 3 | 1094 | 3760 | 11 | 1038 | 0 | 2291 |
| | 19749 | 0 | 202 | 1323 | 3439 | 212 | 2108 | 0 | 1840 |
| | 17163 | 0 | 0 | 1547 | 3162 | 0 | 1677 | 0 | 1688 |
| 'P3' | 9994 | 0 | 0 | 0 | 0 | 0 | 2408 | 1866 | 0 |
| | 10357 | 0 | 0 | 0 | 0 | 0 | 2206 | 2291 | 0 |
| | 9441 | 0 | 0 | 0 | 0 | 0 | 2037 | 1840 | 0 |
| | 9163 | 0 | 0 | 0 | 0 | 0 | 2099 | 1688 | 0 |

Note: Cell surface area is quantified by the total number of pixels surrounding a cell, while cell-cell contact area is quantified by the total number of pixels adjacent to both cells (sample size = 4; spatial resolution ≈ 0.225 μm / pixel in three orthogonal coordinates). "NaN" represents a cell which has its own nucleus but hasn't completely finished cytokinesis with its sister, which together share the same membrane boundary. "0" means that the two independent cells don't contact with each other at all. The cell-cell contacts with no "0" is reproducible in all the 4 segmented wild-type embryo samples (Table S1) (Cao et al., 2019).

**Table S7. Comparison between simulation ($\sigma$ = $\sigma$' = 0.0, 0.3, 0.6, 0.9, 1.2, 1.5) and experiment at 4-cell stage (Figure 2B).**

| $\sigma$ | | Area in Simulation $S_{sim}$ (Pixel Number*Spatial Resolution$^2$, μm$^2$) | | | | | ABS{($S_{sim,\,i}$ - $S_{exp,\,i}$)/($S_{exp,\,i}$)} | | | | | MEAN{ABS{($S_{sim,\,i}$ - $S_{exp,\,i}$)/($S_{exp,\,i}$)}} |
|---|---|---|---|---|---|---|---|---|---|---|---|---|
| | | Contact | | | | Surface | Contact | | | | Surface | |
| | | ABa | ABp | EMS | P2 | | ABa | ABp | EMS | P2 | | |
| 0.0 | ABa | 0.00 | 123.50 | 82.63 | 0.00 | 1494.62 | NaN | 0.625 | 0.686 | NaN | 0.030 | 0.3641 |
| | ABp | 123.50 | 0.00 | 45.15 | 111.05 | 1546.50 | 0.625 | NaN | 0.823 | 0.568 | 0.025 | |
| | EMS | 82.63 | 45.15 | 0.00 | 93.25 | 1317.05 | 0.686 | 0.823 | NaN | 0.244 | 0.146 | |
| | P2 | 0.00 | 111.05 | 93.25 | 0.00 | 1006.85 | NaN | 0.568 | 0.244 | NaN | 0.130 | |
| 0.3 | ABa | 0.00 | 244.67 | 188.39 | 0.00 | 1492.99 | NaN | 0.257 | 0.285 | NaN | 0.031 | 0.2054 |
| | ABp | 244.67 | 0.00 | 158.08 | 206.00 | 1547.63 | 0.257 | NaN | 0.382 | 0.198 | 0.024 | |
| | EMS | 188.39 | 158.08 | 0.00 | 171.98 | 1316.48 | 0.285 | 0.382 | NaN | 0.395 | 0.147 | |
| | P2 | 0.00 | 206.00 | 171.98 | 0.00 | 1006.41 | NaN | 0.198 | 0.395 | NaN | 0.131 | |
| 0.6 | ABa | 0.00 | 291.27 | 229.77 | 0.00 | 1494.37 | NaN | 0.115 | 0.128 | NaN | 0.030 | 0.1705 |
| | ABp | 291.27 | 0.00 | 204.87 | 245.87 | 1550.96 | 0.115 | NaN | 0.199 | 0.043 | 0.022 | |
| | EMS | 229.77 | 204.87 | 0.00 | 212.92 | 1322.83 | 0.128 | 0.199 | NaN | 0.727 | 0.142 | |
| | P2 | 0.00 | 245.87 | 212.92 | 0.00 | 1008.24 | NaN | 0.043 | 0.727 | NaN | 0.129 | |
| 0.9 | ABa | 0.00 | 323.21 | 257.94 | 0.00 | 1500.41 | NaN | 0.018 | 0.021 | NaN | 0.026 | 0.1559 |
| | ABp | 323.21 | 0.00 | 233.54 | 269.95 | 1557.19 | 0.018 | NaN | 0.087 | 0.051 | 0.018 | |
| | EMS | 257.94 | 233.54 | 0.00 | 236.62 | 1329.69 | 0.021 | 0.087 | NaN | 0.919 | 0.138 | |
| | P2 | 0.00 | 269.95 | 236.62 | 0.00 | 1012.01 | NaN | 0.051 | 0.919 | NaN | 0.126 | |
| 1.2 | ABa | 0.00 | 339.50 | 272.09 | 0.00 | 1503.74 | NaN | 0.031 | 0.033 | NaN | 0.024 | 0.1723 |
| | ABp | 339.50 | 0.00 | 253.35 | 288.44 | 1562.03 | 0.031 | NaN | 0.009 | 0.123 | 0.015 | |
| | EMS | 272.09 | 253.35 | 0.00 | 253.22 | 1330.75 | 0.033 | 0.009 | NaN | 1.054 | 0.137 | |
| | P2 | 0.00 | 288.44 | 253.22 | 0.00 | 1013.20 | NaN | 0.123 | 1.054 | NaN | 0.125 | |
| 1.5 | ABa | 0.00 | 353.58 | 286.05 | 0.00 | 1505.82 | NaN | 0.074 | 0.086 | NaN | 0.022 | 0.1932 |
| | ABp | 353.58 | 0.00 | 260.77 | 294.60 | 1561.53 | 0.074 | NaN | 0.020 | 0.147 | 0.015 | |
| | EMS | 286.05 | 260.77 | 0.00 | 260.83 | 1333.08 | 0.086 | 0.020 | NaN | 1.115 | 0.136 | |
| | P2 | 0.00 | 294.60 | 260.83 | 0.00 | 1015.28 | NaN | 0.147 | 1.115 | NaN | 0.123 | |

| | Area in Experiment $S_{exp}$ (Pixel Number*Spatial Resolution$^2$, μm$^2$) | | | | |
|---|---|---|---|---|---|
| | Contact | | | | Surface |
| | ABa | ABp | EMS | P2 | |
| ABa | 0.00 | 329.25 | 263.36 | 0.00 | 1540.24 |
| ABp | 329.25 | 0.00 | 255.71 | 256.91 | 1585.63 |
| EMS | 263.36 | 255.71 | 0.00 | 123.30 | 1542.56 |
| P2 | 0.00 | 256.91 | 123.30 | 0.00 | 1157.87 |

Note: MEAN{} denotes the function that calculates the average value of samples; ABS{} denotes the function that calculates the absolute value of sample.

**Table S8. Comparison between simulation ($\sigma$ = 0.9, $\sigma$' = 0.0, 0.2, 0.4, 0.6, 0.8) and experiment at 4-cell stage (Figure 2C).**

| $\sigma$' | | Area in Simulation $S_{sim}$ (Pixel Number*Spatial Resolution$^2$, μm$^2$) | | | | | ABS{($S_{sim, i}$ - $S_{exp, i}$)/($S_{exp, i}$)} | | | | | MEAN{ABS{($S_{sim, i}$ - $S_{exp, i}$) /($S_{exp, i}$)}} |
|---|---|---|---|---|---|---|---|---|---|---|---|---|
| | | Contact | | | | Surface | Contact | | | | Surface | |
| | | ABa | ABp | EMS | P2 | | ABa | ABp | EMS | P2 | | |
| 0.0 | ABa | 0.00 | 318.56 | 279.76 | 0.00 | 1502.48 | NaN | 0.032 | 0.062 | NaN | 0.025 | 0.1429 |
| | ABp | 318.56 | 0.00 | 245.61 | 275.17 | 1571.34 | 0.032 | NaN | 0.039 | 0.071 | 0.009 | |
| | EMS | 279.76 | 245.61 | 0.00 | 27.42 | 1321.01 | 0.062 | 0.039 | NaN | 0.778 | 0.144 | |
| | P2 | 0.00 | 275.17 | 27.42 | 0.00 | 1011.63 | NaN | 0.071 | 0.778 | NaN | 0.126 | |
| 0.2 | ABa | 0.00 | 323.52 | 277.43 | 0.00 | 1505.75 | NaN | 0.017 | 0.053 | NaN | 0.022 | 0.0629 |
| | ABp | 323.52 | 0.00 | 237.88 | 275.73 | 1571.27 | 0.017 | NaN | 0.070 | 0.073 | 0.009 | |
| | EMS | 277.43 | 237.88 | 0.00 | 130.23 | 1326.73 | 0.053 | 0.070 | NaN | 0.056 | 0.140 | |
| | P2 | 0.00 | 275.73 | 130.23 | 0.00 | 1013.71 | NaN | 0.073 | 0.056 | NaN | 0.125 | |
| 0.4 | ABa | 0.00 | 320.63 | 268.38 | 0.00 | 1501.60 | NaN | 0.026 | 0.019 | NaN | 0.025 | 0.0991 |
| | ABp | 320.63 | 0.00 | 233.67 | 273.78 | 1564.36 | 0.026 | NaN | 0.086 | 0.066 | 0.013 | |
| | EMS | 268.38 | 233.67 | 0.00 | 171.04 | 1323.77 | 0.019 | 0.086 | NaN | 0.387 | 0.142 | |
| | P2 | 0.00 | 273.78 | 171.04 | 0.00 | 1010.88 | NaN | 0.066 | 0.387 | NaN | 0.127 | |
| 0.6 | ABa | 0.00 | 320.25 | 263.22 | 0.00 | 1500.53 | NaN | 0.027 | 0.001 | NaN | 0.026 | 0.1286 |
| | ABp | 320.25 | 0.00 | 232.16 | 273.47 | 1559.89 | 0.027 | NaN | 0.092 | 0.064 | 0.016 | |
| | EMS | 263.22 | 232.16 | 0.00 | 204.87 | 1322.64 | 0.001 | 0.092 | NaN | 0.662 | 0.143 | |
| | P2 | 0.00 | 273.47 | 204.87 | 0.00 | 1011.00 | NaN | 0.064 | 0.662 | NaN | 0.127 | |
| 0.8 | ABa | 0.00 | 321.45 | 258.69 | 0.00 | 1500.85 | NaN | 0.024 | 0.018 | NaN | 0.026 | 0.1506 |
| | ABp | 321.45 | 0.00 | 231.91 | 272.78 | 1557.32 | 0.024 | NaN | 0.093 | 0.062 | 0.018 | |
| | EMS | 258.69 | 231.91 | 0.00 | 228.01 | 1326.79 | 0.018 | 0.093 | NaN | 0.849 | 0.140 | |
| | P2 | 0.00 | 272.78 | 228.01 | 0.00 | 1011.38 | NaN | 0.062 | 0.849 | NaN | 0.127 | |

| | Area in Experiment $S_{exp}$ (Pixel Number*Spatial Resolution$^2$, μm$^2$) | | | | |
|---|---|---|---|---|---|
| | Contact | | | | Surface |
| | ABa | ABp | EMS | P2 | |
| ABa | 0.00 | 329.25 | 263.36 | 0.00 | 1540.24 |
| ABp | 329.25 | 0.00 | 255.71 | 256.91 | 1585.63 |
| EMS | 263.36 | 255.71 | 0.00 | 123.30 | 1542.56 |
| P2 | 0.00 | 256.91 | 123.30 | 0.00 | 1157.87 |

Note: MEAN{} denotes the function that calculates the average value of samples; ABS{} denotes the function that calculates the absolute value of sample.

**Table S9. Preservation time of contact map in simulation with attraction motif at 8-cell stage (Figures 6A, 6B, S10A and S10B).**

| Attraction Motif at 8-Cell Stage | | Preservation Time of Contact Map (Time Step) | Corresponding Time in Experiment (min) |
|---|---|---|---|
| Default | / | 38900 | 2.32 |
| ABal-ABar | ($\sigma$ABal-ABar = 0.9) | 45300 | 2.71 |
| ABal-ABpl | ($\sigma$'ABal-ABpl = 0.2) | 40600 | 2.43 |
| ABal-MS | ($\sigma$'ABal-MS = 0.2) | 98900 | 5.91 |
| ABar-ABpl | ($\sigma$'ABar-ABpl = 0.2) | 53100 | 3.17 |
| ABar-ABpr | ($\sigma$'ABar-ABpr = 0.2) | 30800 | 1.84 |
| ABar-MS | ($\sigma$'ABar-MS = 0.2) | 11200 | 0.67 |
| ABpl-ABpr | ($\sigma$ABpl-ABpr = 0.9) | 38200 | 2.28 |
| ABpl-MS | ($\sigma$'ABpl-MS = 0.2) | 20800 | 1.24 |
| ABpl-E | ($\sigma$'ABpl-E = 0.2) | 150500 | 8.99 |
| ABpl-C | ($\sigma$'ABpl-C = 0.2) | 3500 | 0.21 |
| ABpr-MS | ($\sigma$'ABpr-MS = 0.2) | 6900 | 0.41 |
| ABpr-E | ($\sigma$'ABpr-E = 0.2) | 24700 | 1.48 |
| ABpr-C | ($\sigma$'ABpr-C = 0.2) | 73200 | 4.37 |
| MS-E | ($\sigma$MS-E = 0.9) | 46300 | 2.77 |
| E-P3 | ($\sigma$'E-P3 = 0.2) | 35700 | 2.13 |
| C-E | ($\sigma$'C-E = 0.2) | 9200 | 0.55 |
| C-P3 | ($\sigma$C-P3 = 0.9) | 31800 | 1.90 |

**Table S10. Corresponding timing between simulations with $\sigma'_{ABpl\text{-}E} = 0.2$, $\sigma_{ABpl\text{-}MS} = 0.9$ and $\sigma'_{ABpl\text{-}E} = 0.2$, $\sigma_{ABpl\text{-}MS} = 1.6$ at 8-cell stage (Figures 6C, 6D, and S12).**

| Attraction Motifs at 8-Cell Stage | $\sigma'_{ABpl\text{-}E} = 0.2$, $\sigma_{ABpl\text{-}MS} = 0.9$ | $\sigma$'ABpl-E = 0.2, $\sigma$ABpl-MS = 1.6 |
|---|---|---|
| | 192900 | 190500 |
| | 209900 | 199800 |
| | 226400 | 215000 |
| | 233800 | 222300 |
| | 243200 | 230200 |
| | 248200 | 235400 |
| | 271100 | 257700 |
| | 284100 | 267300 |
| | 294500 | 276000 |
| | 310200 | 285000 |
| | 326700 | 289300 |
| | 335400 | 294700 |
| | 336600 | 296000 |
| | 338200 | 297700 |
| | 345100 | 306500 |
| | 348000 | 311300 |

**Table S11. Genes with "planarization" phenotype at 6-cell stage after perturbation.**

| Gene Name | Embryo Width (μm) | *p*-value (One-Tailed Wilcoxon Rank-Sum Test) | Measured Values |
|---|---|---|---|
| wild-type' | 8.175 ± 1.468' | / | / |
| *glp-1'* | 7.070 ± 0.000' | 0.04201 | 7.070,7.070' |
| *cdk-12'* | 6.363 ± 1.143' | 0.04505 | 5.555,7.171' |
| *ceh-18'* | 7.019 ± 0.071' | 0.03915 | 7.070,6.969' |
| *hmg-1.2'* | 6.969 ± 0.143' | 0.03557 | 7.070,6.868' |
| *egl-27'* | 6.705 ± 0.293' | 0.00381 | 6.666,6.666,6.390,7.100' |
| *fkh-2'* | 5.959 ± 0.143' | 0.01047 | 6.060,5.858' |
| *spr-1'* | 6.868 ± 0.286' | 0.03148 | 6.666,7.070' |
| *ham-1'* | 7.117 ± 0.757' | 0.01200 | 8.383,6.969,7.100,6.745,6.390' |
| *ego-1'* | 6.248 ± 0.803' | 0.01689 | 6.816,5.680' |
| *spe-5'* | 6.106 ± 0.100' | 0.01226 | 6.035,6.177' |
| *rars-2'* | 5.928 ± 1.858' | 0.04663 | 7.242,4.615' |
| *nkcc-1'* | 5.928 ± 1.356' | 0.01599 | 6.887,4.970' |
| *unc-115'* | 6.071 ± 0.351' | 0.01209 | 5.822,6.319' |
| *pfd-1'* | 6.859 ± 1.245' | 0.02396 | 6.319,6.035,8.520,7.810,5.609' |
| *mrp-5'* | 6.852 ± 0.050' | 0.03034 | 6.887,6.816' |
| *kin-19'* | 6.319 ± 0.256' | 0.00424 | 6.106,6.603,6.248' |
| *prp-31'* | 6.532 ± 0.000' | 0.02235 | 6.532,6.532' |
| *C24H12.4'* | 6.567 ± 0.753' | 0.03110 | 6.035,7.100' |
| *cir-1'* | 6.692 ± 0.837' | 0.01428 | 7.597,7.100,5.680,6.390' |
| *dnj-12'* | 6.958 ± 0.201' | 0.04771 | 6.816,7.100' |
| *cdc-73'* | 6.922 ± 0.424' | 0.01734 | 7.100,6.816,7.384,6.390' |
| *F52A8.5'* | 6.780 ± 0.452' | 0.03389 | 6.461,7.100' |
| *F57B9.3'* | 6.745 ± 0.502' | 0.03961 | 6.390,7.100' |
| *rga-5'* | 6.745 ± 0.502' | 0.02959 | 7.100,6.390' |
| *let-504'* | 6.390 ± 0.000' | 0.01643 | 6.390,6.390' |
| *gop-3'* | 6.863 ± 0.820' | 0.03730 | 7.810,6.390,6.390' |
| *fib-1'* | 6.319 ± 0.100' | 0.01689 | 6.248,6.390' |
| *F59C6.5'* | 6.638 ± 0.653' | 0.03307 | 6.177,7.100' |
| *C16C10.8'* | 6.390 ± 1.812' | 0.03803 | 5.680,5.041,8.449' |
| *scpl-1'* | 6.745 ± 0.502' | 0.04826 | 7.100,6.390' |
| *F37C12.3'* | 6.603 ± 0.703' | 0.04152 | 6.106,7.100' |
| *ttll-5'* | 6.141 ± 0.351' | 0.01392 | 5.893,6.390' |
| *dnc-5'* | 7.195 ± 0.434' | 0.04560 | 7.668,6.816,7.100' |
| *tomm-20'* | 6.213 ± 0.753' | 0.01599 | 5.680,6.745' |
| *phf-5'* | 6.745 ± 0.502' | 0.03227 | 7.100,6.390' |
| *chp-1'* | 6.568 ± 0.753' | 0.03110 | 7.100,6.035' |
| *cdk-7'* | 6.390 ± 0.000' | 0.01807 | 6.390,6.390' |
| *cit-1.2'* | 6.993 ± 0.151' | 0.03961 | 6.887,7.100' |
| *cya-1'* | 6.851 ± 0.351' | 0.03472 | 6.603,7.100' |
| *wee-1.1'* | 7.305 ± 0.867' | 0.03100 | 7.668,8.094,6.603,6.390,6.390,6.390,7.597,8.023, 8.591' |

**Table S12. Genes with "planarization" phenotype at 7-cell stage after perturbation.**

| Gene Name | Embryo Width (μm) | *p*-value (One-Tailed Wilcoxon Rank-Sum Test) | Measured Values |
|---|---|---|---|
| wild-type' | 11.257 ± 1.695' | / | / |
| *mex-1'* | 9.935 ± 1.231' | 0.03668 | 11.110,11.413,9.191,8.733,9.230' |
| *gad-1'* | 10.266 ± 1.655' | 0.02757 | 12.625,12.726,11.999,11.644,11.005,7.952,9.088, 8.449,9.585,9.372,9.656,9.088' |
| *glp-1'* | 9.393 ± 0.143' | 0.02860 | 9.292,9.494' |
| *par-1'* | 8.029 ± 0.214' | 0.00807 | 7.878,8.181' |
| *grh-1'* | 9.864 ± 0.875' | 0.02477 | 10.403,10.807,9.017,9.230' |
| *lag-1'* | 10.187 ± 0.964' | 0.01114 | 8.080,11.009,9.940,9.869,11.573,10.437,10.650, 10.295,10.650,9.372' |
| *blmp-1'* | 9.292 ± 0.143' | 0.02101 | 9.393,9.191' |
| *lpd-2'* | 8.787 ± 0.429' | 0.01180 | 9.090,8.484' |
| *egl-27'* | 9.797 ± 0.904' | 0.01644 | 10.504,9.090,10.650,8.946' |
| *fkh-2'* | 8.181 ± 0.143' | 0.00807 | 8.282,8.080' |
| *lsl-1'* | 9.393 ± 1.571' | 0.04951 | 8.282,10.504' |
| *spr-1'* | 9.797 ± 0.143' | 0.04518 | 9.696,9.898' |
| *rps-9'* | 8.855 ± 0.948' | 0.00202 | 7.777,8.686,8.875,10.082' |
| *ZK792.5'* | 9.514 ± 0.502' | 0.02754 | 9.869,9.159' |
| *ego-1'* | 8.839 ± 1.255' | 0.01941 | 9.727,7.952' |
| *spe-5'* | 8.023 ± 0.301' | 0.00807 | 7.810,8.236' |
| *rfc-3'* | 9.195 ± 0.251' | 0.01993 | 9.017,9.372' |
| *aagr-2'* | 9.620 ± 0.452' | 0.03744 | 9.940,9.301' |
| *F19F10.9'* | 9.159 ± 0.402' | 0.02046 | 8.875,9.443' |
| *ifa-1'* | 9.478 ± 0.151' | 0.03238 | 9.585,9.372' |
| *T19D2.2'* | 8.378 ± 1.105' | 0.01083 | 9.159,7.597' |
| *dhfr-1'* | 9.727 ± 0.201' | 0.04264 | 9.869,9.585' |
| *nkcc-1'* | 7.490 ± 1.255' | 0.00831 | 8.378,6.603' |
| *nxf-1'* | 9.691 ± 0.753' | 0.04895 | 10.224,9.159' |
| *pole-2'* | 8.733 ± 0.803' | 0.01418 | 9.301,8.165' |
| *F13H8.9'* | 9.230 ± 0.502' | 0.02303 | 9.585,8.875' |
| *unc-115'* | 9.514 ± 0.502' | 0.02754 | 9.159,9.869' |
| *cul-1'* | 9.287 ± 0.456' | 0.00081 | 9.656,8.662,9.088,9.230,9.798' |
| *pfd-1'* | 9.542 ± 1.919' | 0.01696 | 9.585,8.378,12.638,7.597,9.514' |
| *cwn-2'* | 10.052 ± 0.435' | 0.00802 | 10.224,10.295,9.585,9.372,10.650,10.082,10.153' |
| *dsl-1'* | 9.656 ± 0.402' | 0.04785 | 9.940,9.372' |
| *gsk-3'* | 9.715 ± 1.668' | 0.01523 | 8.946,12.425,10.650,9.940,7.810,8.520' |
| *kin-19'* | 9.088 ± 1.280' | 0.01236 | 9.443,10.153,7.668' |
| *lin-44'* | 8.946 ± 1.406' | 0.02425 | 9.940,7.952' |
| *sma-2'* | 9.053 ± 0.251' | 0.01606 | 8.875,9.230' |
| *rae-1'* | 9.159 ± 1.205' | 0.03198 | 8.307,10.011' |
| *ntl-9'* | 8.910 ± 1.356' | 0.01993 | 7.952,9.869' |
| *sel-9'* | 9.230 ± 0.803' | 0.02363 | 8.662,9.798' |
| *rpn-5'* | 9.230 ± 1.607' | 0.04364 | 10.366,8.094' |
| *B0361.6'* | 9.123 ± 0.151' | 0.01606 | 9.017,9.230' |
| *cdc-73'* | 9.691 ± 1.230' | 0.02500 | 11.360,9.727,9.230,8.449' |
| *rfc-1'* | 9.301 ± 0.904' | 0.02896 | 8.662,9.940' |
| *fars-3'* | 9.265 ± 0.050' | 0.01915 | 9.230,9.301' |
| *gex-3'* | 9.123 ± 0.151' | 0.01606 | 9.017,9.230' |
| *let-504'* | 7.810 ± 0.000' | 0.00759 | 7.810,7.810' |
| *cdc-37'* | 8.520 ± 0.502' | 0.01083 | 8.875,8.165' |
| *gop-3'* | 8.425 ± 0.690' | 0.00242 | 9.017,7.668,8.591' |
| *fib-1'* | 8.555 ± 0.351' | 0.01067 | 8.307,8.804' |
| *Y69A2AR.18'* | 8.485 ± 0.351' | 0.01022 | 8.733,8.236' |
| *F59C6.5'* | 9.088 ± 0.201' | 0.01606 | 9.230,8.946' |
| *C16C10.8'* | 8.556 ± 0.954' | 0.01215 | 7.881,9.230' |
| *scpl-1'* | 9.159 ± 0.602' | 0.02186 | 9.585,8.733' |
| *fat-7'* | 9.123 ± 1.155' | 0.02333 | 9.940,8.307' |
| *mrps-26'* | 9.691 ± 0.753' | 0.04785 | 9.159,10.224' |

| | | | |
|---|---|---|---|
| *F37C12.3'* | 9.053 ± 0.251' | 0.01606 | 8.875,9.230' |
| *ttll-5'* | 9.372 ± 0.201' | 0.02686 | 9.514,9.230' |
| *nath-10'* | 9.396 ± 0.782' | 0.01279 | 8.875,10.295,9.017' |
| *dnc-5'* | 9.301 ± 0.309' | 0.00825 | 8.946,9.514,9.443' |
| *tomm-20'* | 8.910 ± 0.050' | 0.01341 | 8.875,8.946' |
| *clu-1'* | 8.946 ± 0.256' | 0.00336 | 9.017,9.159,8.662' |
| *vps-20'* | 7.502 ± 2.746' | 0.00644 | 10.082,4.615,7.810' |
| *chp-1'* | 8.094 ± 2.410' | 0.01941 | 9.798,6.390' |
| *lin-32'* | 9.549 ± 0.452' | 0.02754 | 9.869,9.230' |
| *F32B5.1'* | 9.301 ± 1.004' | 0.03526 | 8.591,10.011' |
| *rac-2'* | 9.656 ± 0.402' | 0.03880 | 9.372,9.940' |
| *gpx-3'* | 9.585 ± 0.502' | 0.03238 | 9.230,9.940' |
| *swsn-6'* | 9.833 ± 0.151' | 0.04840 | 9.940,9.727' |
| *sys-1'* | 9.751 ± 0.041' | 0.01873 | 9.727,9.798,9.727' |
| *cdc-25.1'* | 7.917 ± 1.155' | 0.00950 | 7.100,8.733' |
| *cdk-7'* | 8.875 ± 0.502' | 0.01418 | 9.230,8.520' |
| *chk-2'* | 9.774 ± 0.148' | 0.02087 | 9.656,9.940,9.727' |
| *cit-1.1'* | 9.052 ± 0.251' | 0.01584 | 9.230,8.875' |
| *cya-1'* | 9.514 ± 0.602' | 0.02825 | 9.088,9.940' |
| *F25H5.5'* | 8.473 ± 0.660' | 0.00245 | 8.165,9.230,8.023' |
| *cdc-48'* | 9.656 ± 0.402' | 0.04840 | 9.940,9.372' |
| *cdc-7'* | 9.372 ± 0.000' | 0.02754 | 9.372,9.372' |
| *wee-1.1'* | 10.011 ± 1.175' | 0.00868 | 11.360,12.070,9.230,9.230,9.585,8.307,10.153, 10.650,9.514' |

**Table S13. Genes with "planarization" phenotype at 8-cell stage after perturbation.**

| Gene Name | Embryo Width (μm) | $p$-value (One-Tailed Wilcoxon Rank-Sum Test) | Measured Values |
|---|---|---|---|
| wild-type' | 11.913 ± 1.865' | / | / |
| *mex-1'* | 9.348 ± 0.556' | 0.00073 | 8.585,10.100,9.595,9.230,9.230' |
| *gad-1'* | 11.048 ± 1.545' | 0.03031 | 14.544,13.029,10.224,11.502,11.644,9.514,9.940, 9.372,11.715,10.579,10.792,9.727' |
| *par-1'* | 8.131 ± 0.786' | 0.00964 | 7.575,8.686' |
| *icd-2'* | 9.949 ± 0.071' | 0.03483 | 9.999,9.898' |
| *nhr-2'* | 9.797 ± 0.571' | 0.03483 | 9.393,10.201' |
| *lpd-2'* | 8.989 ± 0.286' | 0.01438 | 8.787,9.191' |
| *fkh-2'* | 8.433 ± 1.214' | 0.01215 | 7.575,9.292' |
| *rps-9'* | 8.149 ± 1.475' | 0.00024 | 6.262,8.181,7.272,10.082,8.946' |
| *ego-1'* | 9.337 ± 1.155' | 0.02333 | 10.153,8.520' |
| *F19F10.9'* | 8.058 ± 0.753' | 0.00922 | 7.526,8.591' |
| *T19D2.2'* | 10.046 ± 0.151' | 0.04214 | 9.940,10.153' |
| *nkcc-1'* | 9.798 ± 0.000' | 0.02896 | 9.798,9.798' |
| *pole-2'* | 9.905 ± 0.251' | 0.03744 | 10.082,9.727' |
| *unc-115'* | 10.082 ± 0.402' | 0.04623 | 10.366,9.798' |
| *cul-1'* | 10.394 ± 0.654' | 0.01329 | 11.147,10.437,10.863,9.514,10.011' |
| *eif-3.E'* | 9.976 ± 0.351' | 0.03973 | 10.224,9.727' |
| *cye-1'* | 9.709 ± 2.067' | 0.02025 | 11.005,11.928,8.094,7.810' |
| *pad-1'* | 9.833 ± 0.151' | 0.03082 | 9.727,9.940' |
| *rfc-1'* | 9.301 ± 0.904' | 0.02046 | 8.662,9.940' |
| *snfc-5'* | 10.508 ± 1.962' | 0.01082 | 12.780,12.780,10.437,9.372,13.987,12.496,9.869, 8.875,9.514,8.307,9.159,8.520' |
| *gex-3'* | 8.946 ± 0.201' | 0.01399 | 9.088,8.804' |
| *let-504'* | 6.319 ± 0.703' | 0.00759 | 5.822,6.816' |
| *cdc-37'* | 9.407 ± 0.151' | 0.02214 | 9.514,9.301' |
| *gop-3'* | 8.283 ± 1.027' | 0.00256 | 8.946,8.804,7.100' |
| *fib-1'* | 8.236 ± 0.402' | 0.00856 | 7.952,8.520' |
| *F59C6.5'* | 9.869 ± 0.201' | 0.03318 | 10.011,9.727' |
| *C16C10.8'* | 8.485 ± 1.456' | 0.01418 | 7.455,9.514' |
| *scpl-1'* | 8.839 ± 0.452' | 0.01215 | 8.520,9.159' |
| *tbc-1'* | 10.118 ± 0.151' | 0.04677 | 10.224,10.011' |
| *fat-7'* | 10.011 ± 0.602' | 0.04623 | 10.437,9.585' |
| *ttll-5'* | 9.549 ± 0.050' | 0.02585 | 9.514,9.585' |
| *nath-10'* | 8.567 ± 0.823' | 0.00294 | 8.449,7.810,9.443' |
| *dnc-5'* | 9.230 ± 0.710' | 0.00539 | 9.940,8.520,9.230' |
| *tomm-20'* | 9.123 ± 0.151' | 0.01606 | 9.230,9.017' |
| *clu-1'* | 10.011 ± 1.060' | 0.02322 | 9.230,9.585,11.218' |
| *snr-4'* | 9.479 ± 0.351' | 0.02186 | 9.230,9.727' |
| *chp-1'* | 7.810 ± 2.309' | 0.01360 | 9.443,6.177' |
| *rac-2'* | 9.833 ± 0.853' | 0.03880 | 9.230,10.437' |
| *gpx-3'* | 9.620 ± 0.050' | 0.02686 | 9.585,9.656' |
| *swsn-6'* | 10.046 ± 0.251' | 0.04214 | 9.869,10.224' |
| *sys-1'* | 10.153 ± 0.426' | 0.02273 | 9.727,10.153,10.579' |
| *chk-2'* | 9.277 ± 0.675' | 0.00597 | 9.301,9.940,8.591' |
| *cit-1.1'* | 9.123 ± 0.251' | 0.01606 | 9.301,8.946' |
| *cya-1'* | 9.265 ± 0.954' | 0.01967 | 8.591,9.940' |
| *F25H5.5'* | 9.443 ± 0.432' | 0.00661 | 9.230,9.940,9.159' |
| *cdc-48'* | 9.265 ± 0.452' | 0.01915 | 8.946,9.585' |
| *wee-1.1'* | 10.958 ± 0.984' | 0.03464 | 12.567,12.070,11.360,9.940,10.295,10.224,11.573, 10.792,9.798' |
| *cul-4'* | 10.082 ± 0.402' | 0.04623 | 9.798,10.366' |

**Movie S1. Phase-field simulation on the structural evolution of a compressed embryo from 1- to 4-cell stages (Figure 2A).**

**Movie S2. Phase-field simulation on the structural evolution of a compressed embryo from 1 to 4-cell stages, which forms "H" shape with P0 dividing parallel to A-P axis, AB and P1 dividing synchronously and parallel to D-V axis (Figure S3).**

**Movie S3. Phase-field simulation on the structural evolution of a compressed embryo from 1 to 4-cell stages, which forms "I" shape with P0 dividing parallel to A-P axis, AB and P1 dividing synchronously and parallel to A-P axis (Figure S3).**

**Movie S4. Phase-field simulation on the structural evolution of a compressed embryo from 1 to 4-cell stages, which forms "T" shape with P0 dividing parallel to A-P axis, AB and P1 synchronously but parallel to A-P and D-V axes respectively. (Figure S3).**

**Movie S5. Phase-field simulation on the structural evolution of a compressed embryo at 6-cell stage (Figures 3A and S6).**

**Movie S6. Phase-field simulation on the structural evolution of a compressed embryo at 7-cell stage (Figures 3B and 4A).**

**Movie S7. Phase-field simulation on the structural evolution of a compressed embryo at 8-cell stage, without attraction motif on ABpl-E contact (Figures S5 and S7A).**

**Movie S8. Phase-field simulation on the structural evolution of a compressed embryo at 8-cell stage, with attraction motif on ABpl-E contact (Figures 3C and S7B).**

**Movie S9. Phase-field simulation on the structural evolution of an uncompressed embryo at 6-cell stage (Figure S14).**

**Movie S10. Phase-field simulation on the structural evolution of an uncompressed embryo at 7-cell stage (Figure S15).**

**Movie S11. Phase-field simulation on the structural evolution of an uncompressed embryo at 8-cell stage, without attraction motif on ABpl-E contact (Figure S16A).**

**Movie S12. Phase-field simulation on the structural evolution of an uncompressed embryo at 8-cell stage, with attraction motif on ABpl-E contact (Figure S16B).**

**Movie S13. Phase-field simulation on the structural evolution of a compressed embryo from 1 to 8-cell stages, with attraction motif on ABpl-E contact at 8-cell stage (Figures 2A and 3A-3C).**

**Movie S14. Phase-field simulation on the structural evolution of an uncompressed embryo from 1 to 8-cell stages, with attraction motif on ABpl-E contact at 8-cell stage (Figures S13A and S13B).**
